# Multivariate analysis reveals shared genetic architecture of brain morphology and human behavior

**DOI:** 10.1101/2021.04.19.440478

**Authors:** R. de Vlaming, Eric A.W. Slob, Philip R. Jansen, Alain Dagher, Philipp D. Koellinger, Patrick J.F. Groenen, Cornelius A. Rietveld

**Affiliations:** School of Business and Economics, Vrije Universiteit Amsterdam, Amsterdam, The Netherlands; Department of Applied Economics, Erasmus School of Economics, Rotterdam, The Netherlands; Erasmus University Rotterdam Institute for Behavior and Biology, Erasmus School of Economics, Rotterdam, The Netherlands; MRC Biostatistics Unit, School of Clinical Medicine, University of Cambridge, Cambridge, UK; Department of Complex Trait Genetics, Center for Neurogenomics and Cognitive Research, Amsterdam Neuroscience, Vrije Universiteit Amsterdam, Amsterdam, The Netherlands; Department of Clinical Genetics, VU Medical Center, Amsterdam UMC, Amsterdam, The Netherlands; Montreal Neurological Institute, McGill University, Montreal, Quebec, Canada; La Follette School of Public Affairs, University of Wisconsin-Madison, WI, USA; Econometric Institute, Erasmus School of Economics, Rotterdam, The Netherlands

## Abstract

Human variation in brain morphology and behavior are related and highly heritable. Yet, it is largely unknown to what extent specific features of brain morphology and behavior are genetically related. Here, we introduce multivariate genomic-relatedness restricted maximum likelihood (MGREML) and provide estimates of the heritability of grey-matter volume in 74 regions of interest (ROIs) in the brain. We map genetic correlations between these ROIs and health-relevant behavioral outcomes including intelligence. We find four genetically distinct clusters in the brain that are aligned with standard anatomical subdivision in neuroscience. Behavioral traits have distinct genetic correlations with brain morphology which suggests trait-specific relevance of ROIs.

## Introduction

Global and regional grey matter volumes are known to be linked to differences in human behavior and mental health^1^. For example, reduced grey matter density has been implicated in a wide range of neurodegenerative diseases and mental illnesses^2,3,4,5^. In addition, differences in grey matter volume have been related to cognitive and behavioral phenotypic traits such as fluid intelligence and personality, although results have not always been replicable^6,7^.

Variation in brain morphology can be measured non-invasively using magnetic resonance imaging (MRI). Large-scale data collection efforts such as the UK Biobank that included both MRI scans and genetic data enabled recent studies to discover the genetic architecture of human variation in brain morphology and to explore the genetic correlations of brain morphology with behavior and health^8,9,10,11,12^. These studies have demonstrated that all features of brain morphology are genetically highly complex traits and their heritable component is mostly due to the combined influence of many common genetic variants, each with a small effect.

A corollary of this insight is that even the largest currently possible genome-wide association studies (GWASs) identify only a small part of the genetic variants underlying the heritable components of brain morphology: the vast majority of their heritability remains missing^8,9,10,11,12,13^. As a consequence, the genetic correlations of regional brain volumes with each other as well as with human behavior and health have remained largely elusive. However, such estimates could advance our understanding of the genetic architecture of the brain for example regarding its structure and plasticity. Similarly, a strong genetic overlap of specific features of brain morphology with mental health would provide clues about the neural mechanisms behind the genesis of disease^14,15,16^.

We developed multivariate genome-based restricted maximum likelihood method (MGREML) to provide a comprehensive map of the genetic architecture of brain morphology. MGREML overcomes several limitations of existing approaches to estimate heritability and genetic correlations from molecular genetic data. In contrast to existing pairwise bivariate approaches, MGREML guarantees internally consistent (i.e., semi-positive definite) genetic correlation matrices and it yields standard errors that reflect the multivariate structure of the data correctly. The software implementation of MGREML is computationally substantially more efficient than traditional bivariate GREML^17,18^. Moreover, we show that MGREML allows for stronger statistical inference than methods that are based on GWAS summary statistics such as bivariate LD score regression (LDSC)^19,20^. In sum, MGREML yields precise estimates of genetic correlations across a large number of traits when existing approaches applied to the same data are either inaccurate, computationally unfeasible, or underpowered.

We leverage the advantages of MGREML by analyzing brain morphology based on MRI-derived grey-matter volumes in 74 regions of interest (ROIs). We also estimate the genetic correlations of these ROIs with global measures of brain volume and eight human behavioral traits that have well-known associations with mental and physical health. The anthropometric measures height and body-mass index (BMI) are also analyzed, because of their relationships with brain size^6,12^. Our analyses are based on data from the UK Biobank brain imaging study^21^.

## Results

### Estimating genetic correlations

Several methods allow the estimation of heritability and genetic correlations from molecular genetic data. One class of these methods is based on GWAS summary statistics^19,20,22^. Another class of methods is based on individual-level data, such as genome-based restricted maximum likelihood (GREML) and variations of this approach^23,24,25,26,27,28^. Methods based on GWAS summary statistics such as LDSC^19,20^ and variants thereof^29^ can leverage the ever increasing sample sizes of GWAS meta- or mega-analyses^30^ and they are computationally efficient once GWAS results have been obtained. These methods benefit from the fact that GWAS summary statistics are often publicly shared^31,32^. However, the computationally more intensive methods based on individual-level data such as GREML are statistically more powerful^33^.

Where GWAS meta-analysis sample sizes for genetically complex traits such as height^34^ or educational attainment^35^ currently exceed 1 million, most datasets contributing to such a meta-analysis are considerably smaller, which put a constraint on the statistical inferences one can obtain using methods based on individual-level data. Due to the high costs of MRI brain scans, GWAS samples for brain imaging genetics are still relatively small compared to GWAS samples for traits that can be measured at low cost (e.g., height and BMI). The UK Biobank brain imaging study (**Online Methods**) is currently by far the largest available sample that includes both MRI scans and genetic data, often surpassing the sample size of most previous studies in neuroscience by an order of magnitude or more^8,9,12^. This UK Biobank data led to recent breakthroughs in imaging genetics, however even the sample size of the currently largest GWAS of brain morphology is too small to yield precise estimates of genetic correlations with LDSC^10,11^.

Irrespective of whether one uses GWAS summary statistics or individual-level data, the use of bivariate methods poses another challenge when computing genetic correlation across more than two traits. When genetic correlations are estimated across more than two traits, the correlation estimates of pairwise combinations of traits are often aggregated into >2 by >2 genetic correlation matrices^19,20,36^. However, this ‘pairwise bivariate’ approach can result in genetic correlation matrices that are not internally consistent (i.e., they describe interrelationships across traits that cannot exist simultaneously). In mathematical terms, the resulting matrices are not positive semi-definite. Although the correlation between two traits can vary between −1 and +1, their correlation with a third trait is naturally bounded. For a set of three traits, positive semi-definiteness holds when the correlation coefficients satisfy the condition 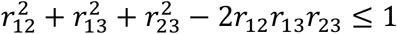. This condition does for instance not hold when pairwise correlations are estimated to be *r*_12_ = 0.9, *r*_13_ = 0.9, and *r*_23_ = 0.2. For example, the genetic correlation matrix in the well-known atlas of genetic correlations is not positive semi-definite^19^. A second consequence of the pairwise bivariate approach is that the standard errors of the resulting genetic correlation matrix do not adequately reflect the multivariate structure of the data.

### MGREML

Our multivariate extension of the GREML method^17,27^ guarantees the internal consistency of the estimated genetic correlation matrix by adopting a factor model for the covariance matrices (**Supplementary Information S1**). This parametrization also ensures that the standard errors of the estimated genetic correlations reflect the multivariate structure of the data correctly. Therefore, methods such as genomic-SEM^37^ that use multivariate genetic correlation matrices as input information may benefit from using results obtained with MGREML. To deal with the computational burden and to make MGREML applicable to large data sets in terms of individuals and traits, we derived efficient expressions for the likelihood function and developed a new optimization algorithm (**Supplementary Information S1**). Runtime analyses described in **Supplementary Information S3** show that MGREML is computationally faster than pairwise bivariate GREML. Increasing the number of individuals *N* in the analyses increases the running time of both pairwise bivariate GREML and MGREML at approximately the same rate, but, when increasing the number of traits *T*, the running time of pairwise bivariate GREML increases disproportionally as compared to MGREML. Moreover, comparison of results obtained with MGREML with results obtained using LDSC shows that standard errors obtained with MGREML are 38.9%-46.0% smaller, illustrating the substantial gains in statistical power afforded by MGREML.

### Analysis of brain morphology

We used MGREML to analyze the heritability of and genetic correlations across 86 traits in 20,190 unrelated ‘white British’ individuals from the UK Biobank (**Fig. 1**, **Online Methods**). The subset of 76 brain morphology traits includes total brain volume (grey and white matter), total grey matter volume, and grey matter volumes in 74 regions of interest (ROIs) in the brain. Relative volumes were obtained by dividing ROI grey matter volumes by total grey matter volume. The full set of heritability estimates is available in **Extended Data Table 1**. **Fig. 2a** and **Fig. 2b** show that SNP-based heritability 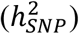 (i.e., the proportion of phenotypic variance which can be explained by autosomal SNPs) is on average highest in the insula, and in the cerebellar and subcortical structures of the brain (average 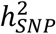 is 33.1%, 32.4%, and 29.5%, respectively) and lowest in the parietal, frontal and temporal lobes of the cortex (average 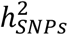 is 21.2%, 21.4%, and 25.2%, respectively). Grouping of the 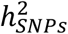 estimates in networks of intrinsic functional connectivity^38^ reveals that ROIs in the heteromodal cortex (frontoparietal, dorsal attention) are less heritable than primary (visual, somatomotor), subcortical and cerebellar regions (**Fig. 3a)**.

**Fig. 1.**
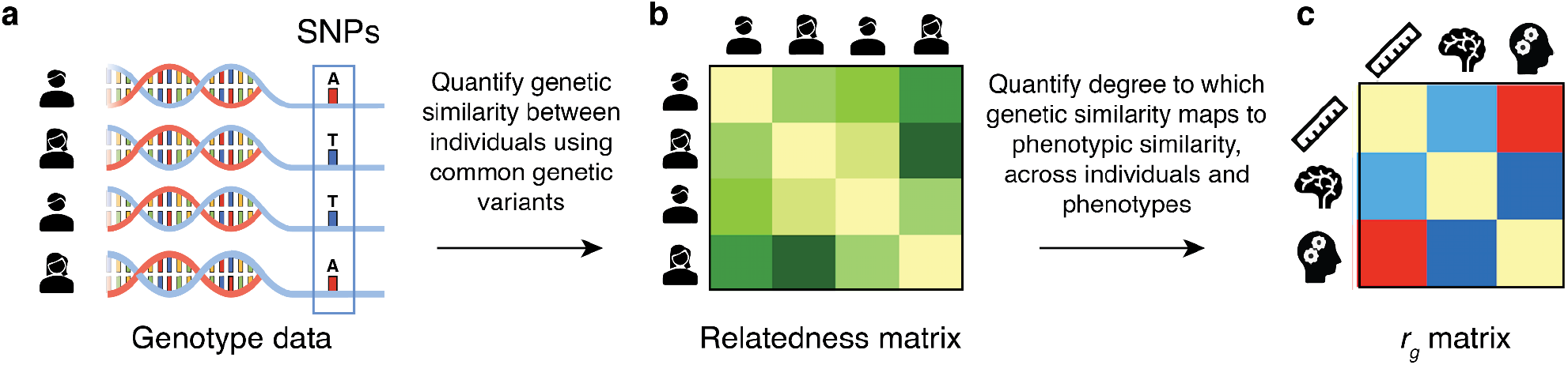
Visualization of multivariate genomic-relatedness restricted maximum likelihood (MGREML). Common genetic variants (single-nucleotide polymorphisms) in the human genome (Panel a) are used to construct a Genomic-Relatedness Matrix (GRM) capturing pairwise genetic similarity between individuals in the sample (Panel b). MGREML uses this GRM to jointly estimate heritabilities of phenotypes and genetic correlations (*r_g_*) across multiple phenotypes (Panel c), by quantifying the degree to which genetic similarity maps to phenotypic similarity (across all individuals and phenotypes in the sample). In our empirical application, 1,384,830 common SNPs are used to analyze the genetic correlations across *T*=86 phenotypes in a sample of *N*=20,190 unrelated individuals.

**Fig. 2.**
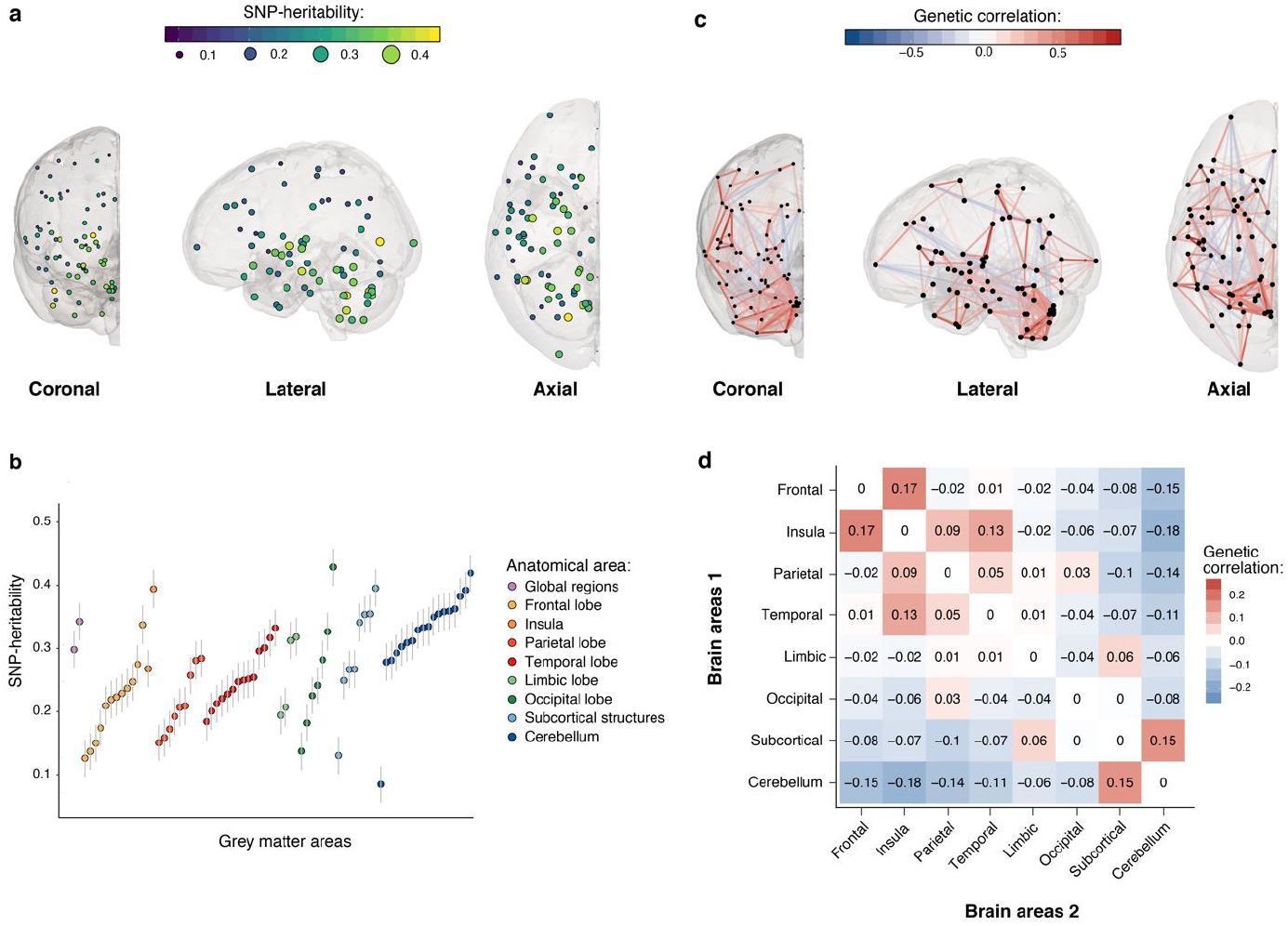
Spatial mapping of SNP-based heritability and genetic correlation estimates obtained using MGREML (*N*=20,190) of relative grey matter volumes in different cortical and subcortical brain areas. **a**. SNP-based heritability of relative grey matter volume mapped to the respective brain region in three dimensions. Each dot represents an area, the color and size represent the heritability of that area. **b.** SNP-based heritability and standard error of relative grey matter volume of each brain region grouped by global anatomical area. **c.** Genetic correlations between cortical and subcortical relative grey matter volumes. The opacity and color represent the strength of the genetic overlap between these two areas (blue vertices represent a negative correlation, red vertices a positive correlation). Only genetic correlations larger than |0.25| are shown. **d.** Average genetic correlations in broad anatomical areas of the brain.

**Fig. 3.**
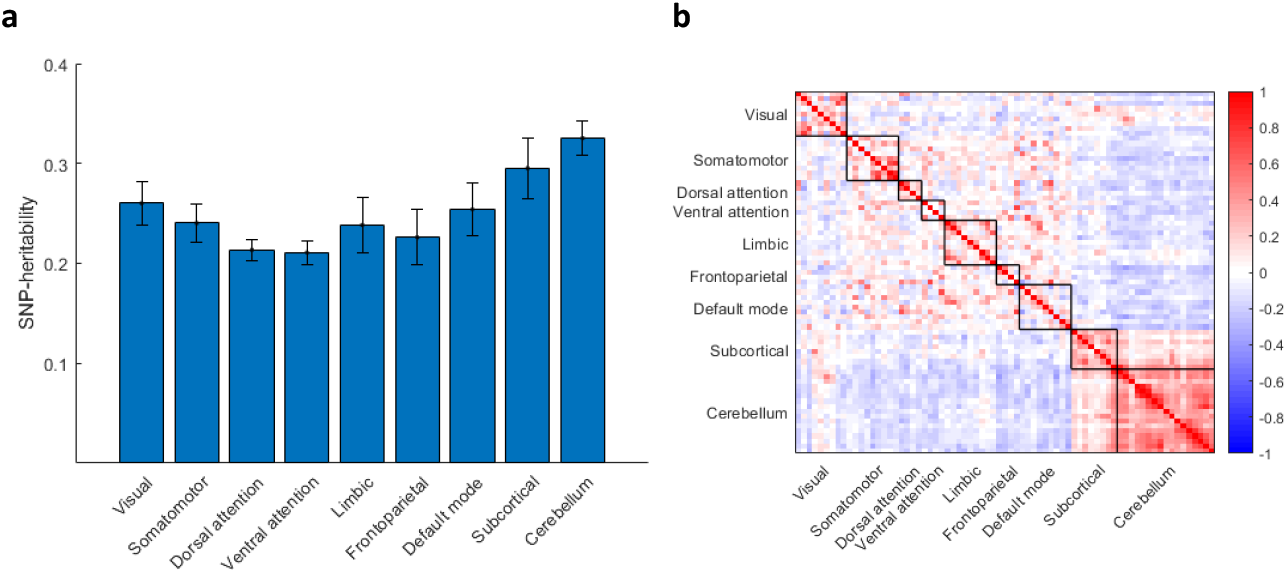
Mapping of SNP-based heritability and genetic correlation estimates obtained using MGREML (*N*=20,190) of relative grey matter volumes in networks of intrinsic functional connectivity. **a**. Average SNP-based heritability of relative grey matter volume in networks of intrinsic functional connectivity. **b.** Genetic correlations in the brain in networks of intrinsic functional connectivity (blue vertices represent a negative correlation, red vertices a positive correlation).

The full set of estimated genetic correlations is available in **Extended Data Table 1**. Using spatial mapping, **Fig. 2c** visualizes the estimated genetic correlations across the relative volumes of the cortical and subcortical brain areas. The largest positive genetic correlations were found between the insular and frontal regions (average *r_g_* = 0.17) and between the cerebellar and subcortical areas (average *r_g_* = 0.15). The largest negative correlations were present between the cerebellar and insular regions (average *r_g_* = −0.18) and between the cerebellar and frontal regions (average *r_g_* = −0.15) (**Fig. 2d**). **Fig. 3b** shows that the genetic correlations are particularly strong within intrinsic connectivity networks, especially the visual, somatomotor, subcortical, and cerebellum networks, possibly because of lower experience-dependent plasticity in these brain regions compared to heteromodal and associative areas^39^. Using Ward’s method for hierarchical clustering^40^, we identify four clusters within the estimated genetic correlations for the 74 ROIs in the brain (**Fig. 4**). The first cluster (*n*=18) includes most of the frontal cortical areas of the brain, the second (*n*=18) the cerebellar cortex, the third (*n*=18) subcortical structures including the brain stem, and the last cluster (*n*=20) contains a mixture of temporal and occipital brain areas.

**Fig. 4.**
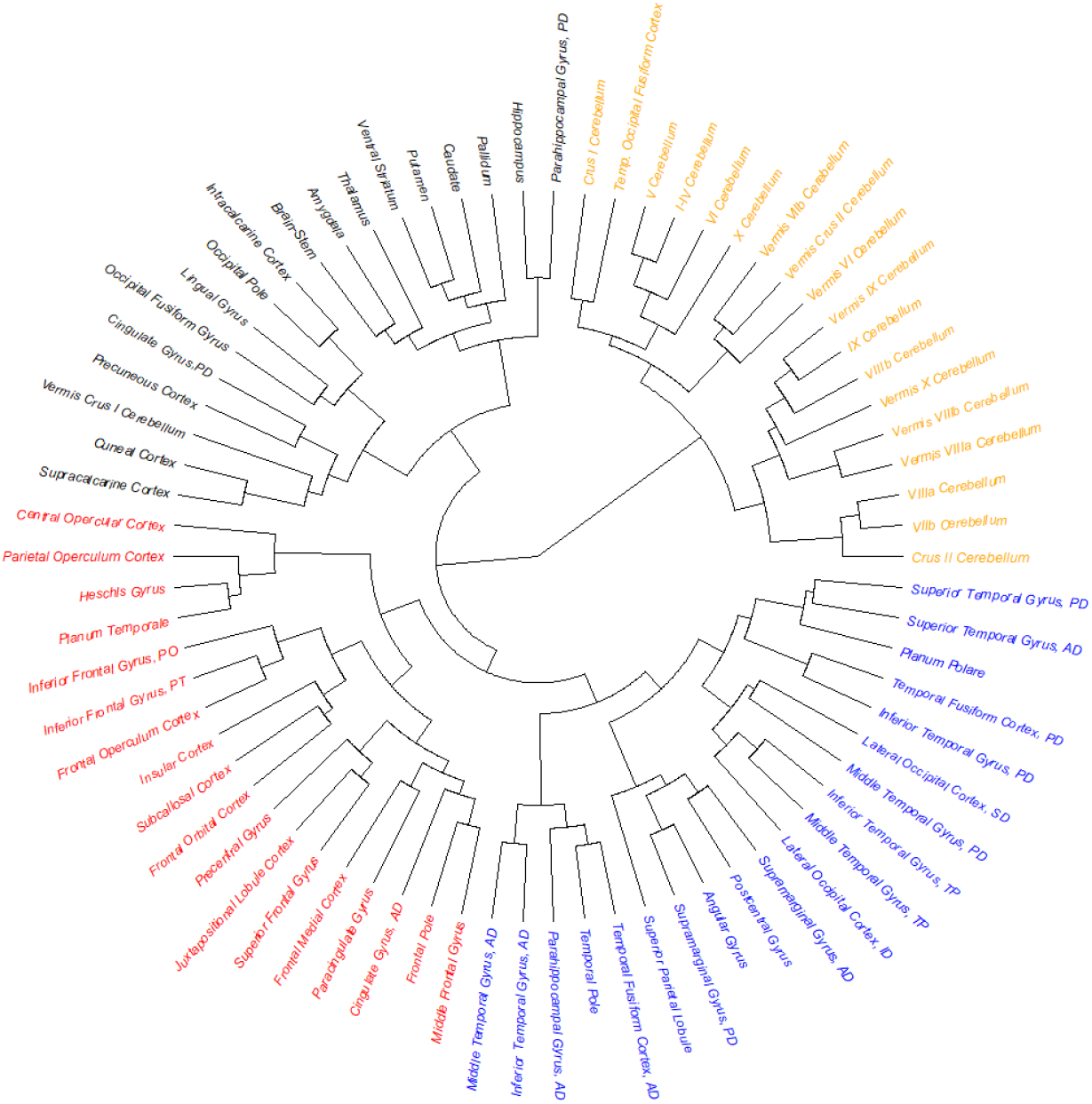
Dendogram of the estimated genetic correlations for the relative grey matter volumes of the 74 regions of interest in the brain. Genetic correlations are estimated using MGREML, and clusters are identified using Ward’s method with a D2 ward for hierarchical clustering. Each color represents a different cluster.

We also used MGREML to estimate the genetic correlations between brain morphology and eight human behavioral traits that are known to be related to health and that have previously been studied in large-scale GWASs, as well as the anthropometric measures height and BMI. Statistically significant correlations are highlighted in **Extended Data Table 1** (Panel C). Spatial maps of the genetic correlation between brain morphology and the behavioral traits are shown in **Fig. 5**. For subjective well-being, we find the strongest genetic correlation with the Middle Frontal Gyrus (**Fig. 5a**, *r_g_* = 0.21), a region that has been linked before to emotion regulation^41^. The genetic correlations of the ROIs with neuroticism (**Fig. 5b**) and depression (**Fig. 5c**) are generally weak and insignificant, perhaps reflecting the coarseness of these phenotypic measures in the UK Biobank data. The strongest genetic correlation with the number of alcoholic drinks consumed per week is with the Lateral Occipital Cortex, superior and inferior divisions (**Fig. 5d**, *r_g_* = 0.23 and *r_g_* = 0.18, respectively). Although the phenotypic correlations between the analyzed ROIs and alcohol consumption are generally negative^42^, these particular brain regions are among those implicated in the affective response to drug cues based on the perception-valuation-action model^43^. For educational attainment and intelligence, the strongest correlations are found in the frontal lobe region (*r_g_* = −0.13 between educational attainment and the Superior Frontal Gyrus, and *r_g_* = 0.16 between intelligence and the Frontal Medial Cortex). **Fig. 5e** and **Fig. 5f** show that the genetic correlation structures estimated for educational attainment and intelligence are largely similar, in line with earlier studies showing the strong genetic overlap between these two traits^44^. Genetic correlations of the ROIs with visual memory (**Fig. 5g**) are insignificant, and the strongest genetic correlation of reaction time is with the Middle Temporal Gyrus, temporooccipital part (**Fig. 5h**, *r_g_* = 0.20). Activity within the middle temporal gyrus has been linked before with reaction time^45^.

**Fig. 5.**
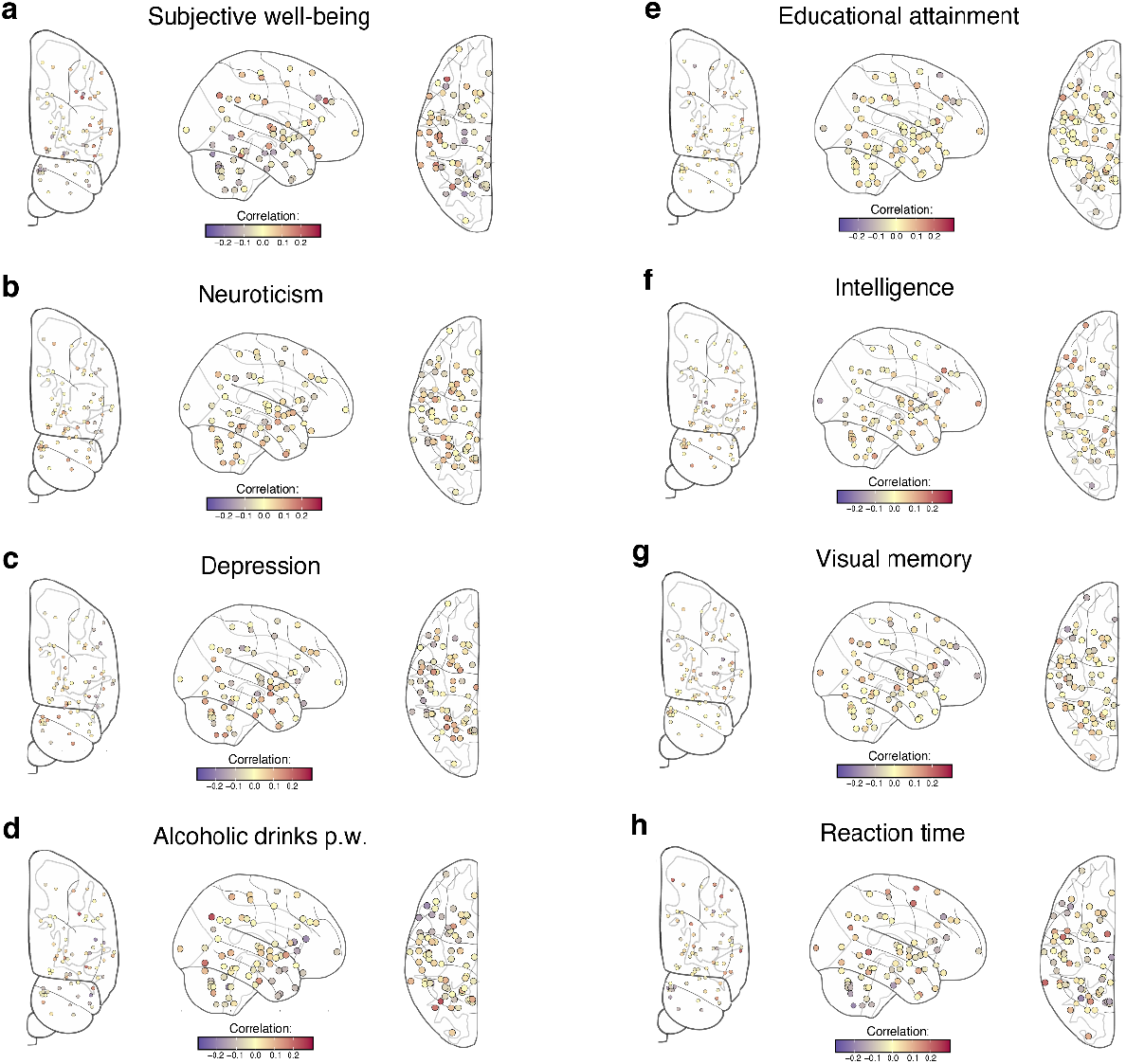
Spatial mapping of genetic correlation estimates obtained using MGREML (*N*=20,190) of relative grey matter volumes of the 74 regions of interest in the brain and 8 behavioral traits. Blue and red points represent negative and positive genetic correlations, respectively. **a**, Subjective well-being. **b**, Neuroticism. **c,** Depression. **d**, Alcoholic drinks per week. **e**, Educational attainment. **f,** Intelligence. **g**, Visual spatial memory. **h**, Reaction time.

Earlier studies suggest that the size of the brain is positively associated with traits such as intelligence^6^, and our use of relative measures may have attenuated the estimation of these relationships in our analyses. When analyzing absolute brain volumes of the ROIs rather than relative brain volumes (i.e., relative to total grey matter volume in the brain), we indeed observe robust positive relationships between the absolute volumes of the ROIs on the one hand and height and intelligence on the other hand (**Extended Data Table 2**). The main differences we observe in the set of estimated correlations across the ROIs are that the genetic correlations within the cerebellum clusters are slightly smaller and that the positive correlations within the subcortical structures are somewhat larger.

## Discussion

We designed MGREML to estimate high dimensional genetic correlation matrices from large-scale individual-level genetic data in a computationally efficient manner while guaranteeing the internal consistency of the estimated genetic correlation matrix. For comparison, we used GWAS and bivariate LDSC^20^ to obtain a genetic correlation matrix with the pairwise bivariate approach using the exact same set of individuals (*N*=20,190) and traits (*T*=86) as in our main analysis (**Extended Data Table 3**). The correlation between the heritability estimates obtained by MGREML and LDSC is 0.95, but the 95% confidence interval of *h*^2^ estimates obtained with MGREML are on average 38.9% smaller than those obtained from LDSC. The 95% confidence intervals of the genetic correlations obtained using MGREML are on average 46.0% smaller compared to those obtained with LDSC, illustrating the advantages of MGREML in terms of statistical power (1,519 versus 1,044 significant correlations at the 5% level). This gain in statistical efficiency is slightly larger than the efficiency gain a recently developed variant of bivariate LDSC was able to achieve^29^. Importantly, the genetic correlation matrix obtained using bivariate LDSC is not positive semi-definite and thus the estimated genetic correlations across traits are not internally coherent. The use of this correlation matrix would pose challenges for multivariate methods such as Genomic-SEM^37^ that are currently based on genetic correlation matrices estimated by LDSC.

Our results show marked variation in the estimated heritability across cortical grey matter volumes, with on average higher heritability estimates in subcortical and cerebellar areas than in cortical areas (**Fig. 2b**). Grouping of by 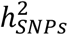 estimates by networks of intrinsic functional connectivity suggests that heritability is particularly low in brain areas with presumed stronger experience-dependent plasticity (**Fig. 3a**). These results suggest that neocortical areas of the brain are under weaker genetic control perhaps reflecting greater environmentally-determined plasticity^39,46^. Furthermore, the estimated genetic correlations suggest the presence of four genetically distinct clusters in the brain (**Fig. 4**). These clusters largely correspond with the conventional subdivision of the brain in different lobes based on anatomical borders^47^. The estimated genetic correlations also provide evidence for a shared genetic architecture of traits between which an association has been observed before in phenotypic studies such as between intelligence and educational attainment^44^. In addition, new genetic correlations were identified between alcohol consumption and cerebellar volume, and between subjective well-being and the temporo-occipital part of the Middle Temporal Gyrus (**Extended Data Table 1**).

To verify that our results are not merely a reflection of the physical proximity of brain regions, we regressed the estimated genetic correlations on the physical distance between the different brain regions. Although this correction procedure decreased the estimated genetic correlations by approximately 25%, the main patterns are still observed. For the same reason, we recreated the dendogram (**Fig. 3**) after aggregating the results for sub regions into an average for the larger region because the optimization procedure of MGREML puts equal weight on each trait and does not account for physical proximity. The results of this robustness check show that the four identified clusters do not merely reflect the number of analyzed measures for a specific brain region.

Estimates of heritability increase our understanding of the relative impact of genetic and environmental variation on traits^13,27^, and estimates of genetic correlation lead to a better understanding of the shared biological pathways between traits^48^. MGREML has been designed to estimate both SNP-based heritability and genetic correlations in a computational efficient and internally consistent manner using individual-level genetic data. Its newly developed optimization algorithm makes it possible to apply MGREML to estimate high dimensional genetic correlation matrices in large datasets such as the UK Biobank.

## Data and code availability

Individual-level genotype and phenotype data are available by application from the UKB Biobank (https://www.ukbiobank.ac.uk/). MGREML is available on https://github.com/devlaming/mgreml as a ready-to-use command-line tool. The GitHub page comes with a full tutorial on the usage of this tool. A MGREML analysis of 86 traits, observed in a sample of 20,190 unrelated individuals (i.e., the dataset we exploit in our empirical application), takes around four hours on a four-core laptop with 16GB of RAM.

## Acknowledgements

UK Biobank has obtained ethical approval from the National Research Ethics Committee (11/NW/0382). This research has been conducted using the UK Biobank Resource under application number 11425. We would like to thank the participants and researchers from UK Biobank Imaging Study who significantly contributed or collected data. This work was carried out on the Dutch national e-infrastructure with the support of SURF Cooperative (NWO Call for Compute Time EINF-403 to E.A.W.S.). P.D.K. and R.d.V. were supported by a European Research Council Consolidator Grant (647648 EdGe to P.D.K.). P.D.K. was also supported by the Office of the Vice Chancellor for Research and Graduate Education at the University of Wisconsin–Madison with funding from the Wisconsin Alumni Research Foundation. C.A.R. was supported by a European Research Council Starting Grant (946647 GEPSI).

## Online Methods

### Statistical framework

In a genome-wide association study (GWAS) of a quantitative trait *y*, the effect of SNP *m* on is modelled as:

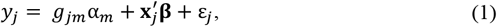

where *y_j_* is the phenotypic value for individual *j*, and *g_jm_* is the genotype for individual *j* for SNP *m* (i.e., a value equal to zero, one, or two, indicating the number of copies of the coded allele the individual carries) with effect α_*m*_ on the phenotype, 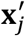 is a 1×*k* vector of control variables with effects **β**, and ε_*j*_ is a normally distributed error term. Instead of one equation per SNP, we can also consider the contribution of all SNPs jointly:

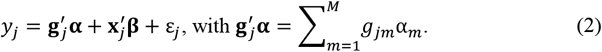

Here, ***g′_j_*** is the 1×*M* vector of genotypes for individual *j* with standardized effects **α**. Assuming the phenotype is either mean-centered and/or an intercept is included in the set of control variables, we can assume, without loss of generality, that SNPs are standardized in accordance with their distribution under Hardy–Weinberg equilibrium. Equation 2 can then be rewritten in matrix notation for a sample comprising *N* individuals as:

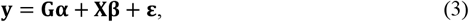

where **G** is the *N*×*M* matrix of standardized genotypes with effects **α**, **X** is the *N*×*k* matrix of control variables with effects **β**, and **ε** is the error term. Following Yang et al. (2010) in their original development of genome-based genomic-relatedness restricted maximum likelihood (GREML)^27^, we assume **β** to be fixed, 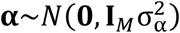 and 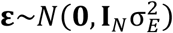. Here, 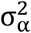 is the variance in the effects of the genetic variants and 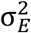 is the environmental variance. In the resulting linear mixed model (LMM), the total genetic contribution 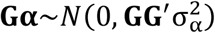. Moreover, the phenotypic variance-covariance matrix across individuals can be decomposed as:

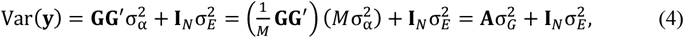

where **A =*M*^−1^ GG′** is the genomic-relatedness matrix (GRM) capturing genetic similarity between individuals based on all SNPs. In Equation 4, 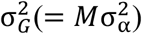 can be interpreted as the contribution of additive genetic effects to the phenotypic variance. Therefore, the SNP-based heritability of the trait of interest *y* can be defined as:

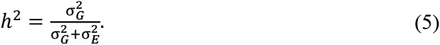

By combining Equations 3 and 4, we can write the LMM as:

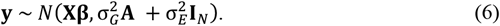

For two quantitative traits, observed in the same set of *N* individuals, this model can be generalized to the following bivariate model^17^:

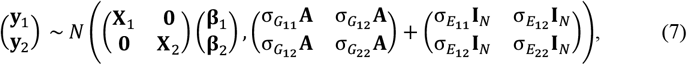

with **X**_1_ (resp. **X**_2_) the *N*-by-*k*_1_ (*N*-by-*k*_2_) matrix of control variables for Trait 1 (2) and associated fixed effects **β**_1_ (**β**_2_), *σ_G_st__* the genetic covariance between traits *s* and *t*, and *σ_E_st__* the environmental covariance between traits *s* and *t*, for *s* = 1, 2 and *t* = 1, 2. The Kronecker product (denoted by ‘⊗’) can be used to extend the bivariate model in Equation 7 to a multivariate model for *T* different traits (i.e., **y**_*t*_ for *t* = 1, …, *T*), as follows^49^:

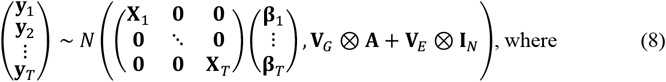

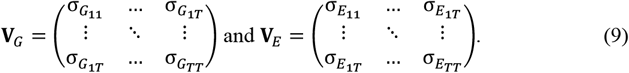

In the multivariate model, SNP-based heritability of trait *t*, denoted by 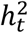, and the genetic correlation between traits *t* and *s*, denoted by *ρ_G_st__*, are defined as:

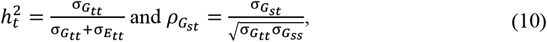

for *t* = 1, …, *T* and *s* = 1, …, *T*.

### Estimation procedure

To estimate the genetic and environmental covariance matrices **V_*G*_** and **V_*E*_** in Equations 8 and 9, we use restricted maximum likelihood (REML) estimation. As optimization method for our REML function, we employ a quasi-Newton approach, using a Broyden–Fletcher–Goldfarb–Shanno (BFGS) algorithm^50^. **Supplementary Information S1** provides efficient expressions for the log-likelihood and gradient to optimize the model with a time complexity that scales linearly with the number of observations and quadratically with the number of traits. The optimization procedure guarantees that the estimated matrices **V_*G*_** and **V_*E*_** are positive semi-definite, by imposing an underlying factor model for both components. After optimization, standard errors can be calculated with a computational complexity that scales linearly with the number of observations and quadratically with the number of parameters in the model (which in turn scales quadratically with the number of traits).

MGREML is available as command-line tool via https://github.com/devlaming/mgreml. Runtime analyses reported in **Supplementary Information S3** show that MGREML is computationally faster than pairwise bivariate GREML. Moreover, comparison with results obtained using LD-score regression^20^ shows that MGREML provides relatively tight confidence intervals for the heritability estimates and genetic correlation estimates.

### Sample and data

UK Biobank is a prospective cohort study in the UK that collects physical, health and cognitive measures, and biological samples (including genotype data) in about 500,000 individuals^51^. In 2016, UK Biobank started to collect brain imaging data with the aim to scan 100,000 subjects by 2022^21,52^. We selected 43,691 individuals with available genotype data from the UK Biobank brain imaging study who self-identified as ‘white British’ and with similar genetic ancestry based on a principal component analysis. After stringent quality control (**Supplementary Information S4**), we estimated pairwise genetic relationships using 1,384,830 autosomal common (Minor Allele Frequency ≥ 0.01) SNPs and retained 37,392 individuals whose pairwise relationship was estimated to be less than 0.025 (approximately corresponding to second- or third-degree cousins or more distant shared ancestry). From these unrelated individuals, we retained the 20,190 individuals with complete information on all 86 traits in our analyses. A description of all the variables used in the empirical analyses is available in **Supplementary Information S2**. Mapping of each cortical region to a network of intrinsic functional connectivity (**Fig. 3**) is based on the assignment of each brain parcel in the Harvard-Oxford atlas^53^ to the intrinsic functional connectivity network^38^ with the highest overlap.

