## Supplementary Information for "Multivariate analysis reveals shared genetic architecture of brain morphology and human behavior"

### S1 Supplementary methods

Genome-based restricted maximum likelihood (GREML) estimation, as developed and introduced by Yang et al. (2010), quantifies the degree to which genetic similarity between individuals maps to phenotypic similarity. Bivariate GREML has been developed to additionally estimate genetic correlations between combinations of two traits (Lee et al., 2012a). We generalise bivariate GREML to a multivariate version, MGREML, which can be used to analyse a large number of quantitative traits simultaneously. The derivations presented in this section reflect the implementation of our method in Python 3.x (available via `https://github.com/devlaming/mgreml`).

A maximum likelihood approach is taken to jointly estimate the genetic and environmental covariance between multiple traits. In this section, efficient expressions are presented that are fundamental to make MGREML estimation computationally feasible for large data sets. Most importantly, a canonical transformation and a commutation matrix are used to transform the full covariance matrix across traits and observations into a block-diagonal matrix.

As an optimisation method, we employ the quasi-Newton approach of the Broyden–Fletcher–Goldfarb–Shanno (BFGS) algorithm. For this algorithm, computationally efficient expressions are needed for the log-likelihood and the gradient. Moreover, to compute standard errors of our estimates, we need efficient expressions to calculate the information (AI) matrix, or an appropriate approximation thereof.

We take the following steps to obtain an implementation of MGREML that can perform BFGS steps in $O\left(NT^2\right)$ time and calculate the standard errors of heritabilities and genetic correlations in $O\left(NT^4\right)$ time, where $N$ denotes the sample size and $T$ the number of phenotypes included in the analysis:

1. Set up the multivariate model used in MGREML.
2. Transform phenotypes using eigenvectors from the genomic-relatedness matrix (GRM) $\mathbf{A}$ and re-order the observations, in such a way that the grand phenotypic covariance matrix across all traits and observations is block-diagonal.
3. Define the log-likelihood function as well as its gradient and the average information (AI) matrix.
4. Develop tractable notation for different covariates across traits.
5. Find efficient expressions to calculate the log-likelihood.
6. Parametrise the covariance matrices $\mathbf{V}_G$ and $\mathbf{V}_E$ to guarantee their positive semi-definiteness.

7. Given the parametrisation, find efficient expressions to calculate the gradient.
8. Given the parametrisation, find efficient expressions to calculate the AI matrix.
9. Maximise the log-likelihood using a BFGS algorithm.
10. Use a delta method to compute standard errors of estimated heritabilities, genetic correlations, and environmental correlations.

In the next subsections, we discuss these steps one by one in more detail. Throughout the derivations, we will make use of the fundamental property of linear combinations of multivariate normally distributed vectors, that is, if vector $\boldsymbol{\delta} \sim \mathcal{N}(\boldsymbol{\mu}, \boldsymbol{\Sigma})$, then the linear combination $\mathbf{C}\boldsymbol{\delta} + \mathbf{m} \sim \mathcal{N}(\mathbf{C}\boldsymbol{\mu} + \mathbf{m}, \mathbf{C}\boldsymbol{\Sigma}\mathbf{C}^\top)$.

#### Step 1. Setting up the multivariate model

The original model for bivariate GREML, as developed by Lee et al. (2012a), for two normally distributed quantitative traits, $Y_1$ and $Y_2$, observed in the same set of $N$ unrelated individuals, can be written as follows:

$$\begin{pmatrix} \mathbf{y}_1 \\ \mathbf{y}_2 \end{pmatrix} \sim \mathcal{N}\left(\begin{pmatrix} \mathbf{X}_1 & \mathbf{0} \\ \mathbf{0} & \mathbf{X}_2 \end{pmatrix}\begin{pmatrix} \boldsymbol{\beta}_1 \\ \boldsymbol{\beta}_2 \end{pmatrix}, \begin{pmatrix} \sigma_{G_{11}}\mathbf{A} & \sigma_{G_{12}}\mathbf{A} \\ \sigma_{G_{12}}\mathbf{A} & \sigma_{G_{22}}\mathbf{A} \end{pmatrix} + \begin{pmatrix} \sigma_{E_{11}}\mathbf{I}_N & \sigma_{E_{12}}\mathbf{I}_N \\ \sigma_{E_{12}}\mathbf{I}_N & \sigma_{E_{22}}\mathbf{I}_N \end{pmatrix}\right), \tag{1}$$

where parameter $\sigma_{G_{11}}$ (resp. $\sigma_{G_{22}}$) denotes the genetic variance of trait $Y_1$ ($Y_2$) and $\sigma_{G_{12}}$ denotes the genetic covariance of traits $Y_1$ and $Y_2$. The parameters with subscript $E$ are defined in an analogous manner for the environmental variances and covariance. Matrix $\mathbf{A}$ denotes the $N \times N$ genomic-relatedness matrix (GRM) and Matrix $\mathbf{I}_N$ denotes the $N \times N$ identity matrix.. Vector $\mathbf{y}_1$ (resp. $\mathbf{y}_2$) denotes the $N \times 1$ vector of outcomes for trait $Y_1$ ($Y_2$), and matrix $\mathbf{X}_1$ (resp. $\mathbf{X}_2$) denotes the fixed-effect covariates with effects $\boldsymbol{\beta}_1$ ($\boldsymbol{\beta}_2$) for $Y_1$ ($Y_2$).

Eq. 1 can be written more compactly using the Kronecker product (denoted by $\otimes$) and by applying the vectorisation operator (denoted by $\operatorname{vec}()$, where $\operatorname{vec}([\mathbf{v}_1 \quad \dots \quad \mathbf{v}_b]) = \begin{bmatrix}\mathbf{v}_1^\top & \dots & \mathbf{v}_b^\top\end{bmatrix}^\top$) to the $N \times 2$

matrix of phenotypes, $\mathbf{Y}^*$. That is:

$$\text{vec}\left(\mathbf{Y}^*\right) \sim \mathcal{N}\left(\mathbf{X}^*\boldsymbol{\beta}, \mathbf{V}_G \otimes \mathbf{A} + \mathbf{V}_E \otimes \mathbf{I}_N\right), \text{ where}$$

$$\mathbf{Y}^* = \begin{pmatrix} \mathbf{y}_1^* & \mathbf{y}_2^* \end{pmatrix}, \quad \mathbf{X}^* = \begin{pmatrix} \mathbf{X}_1^* & \mathbf{0} \\ \mathbf{0} & \mathbf{X}_2^* \end{pmatrix}, \quad \boldsymbol{\beta} = \begin{pmatrix} \boldsymbol{\beta}_1 \\ \boldsymbol{\beta}_2 \end{pmatrix},$$

$$\mathbf{V}_G = \begin{pmatrix} \sigma_{G_{11}} & \sigma_{G_{12}} \\ \sigma_{G_{12}} & \sigma_{G_{22}} \end{pmatrix}, \text{ and } \mathbf{V}_E = \begin{pmatrix} \sigma_{E_{11}} & \sigma_{E_{12}} \\ \sigma_{E_{12}} & \sigma_{E_{22}} \end{pmatrix}.$$

Here, $\mathbf{V}_G$ is the $2 \times 2$ genetic covariance matrix and $\mathbf{V}_E$ the $2 \times 2$ environmental covariance matrix. This model can easily be generalised to a model for $T$ normally distributed quantitative traits, all observed in the same set of $N$ individuals, in an $N \times T$ matrix $\mathbf{Y}^*$. That is:

$$\text{vec}\left(\mathbf{Y}^*\right) \sim \mathcal{N}\left(\mathbf{X}^*\boldsymbol{\beta}, \mathbf{V}^*\right), \text{ where } \mathbf{V}^* = \mathbf{V}_G \otimes \mathbf{A} + \mathbf{V}_E \otimes \mathbf{I}_N, \tag{2}$$

and where $\mathbf{X}^*$ is a block-diagonal matrix, comprising blocks $\mathbf{X}_t^*$ for traits $t = 1, \ldots, T$, where $\mathbf{X}_t^*$ is the $N \times K_t$ matrix of fixed-effects covariates for trait $t$, where $K_t$ denotes the number of fixed-effect covariates that apply to trait $t$. In this model, $\mathbf{V}_G$ denotes the $T \times T$ genetic covariance matrix across the $T$ traits and $\mathbf{V}_E$ the environmental covariance matrix.

#### Step 2. Transforming and re-ordering phenotypes

Using a canonical transformation (Thompson, 1977), such that we transform the data to be independent across individuals (rather than across traits), we can introduce sparsity in our model. That is, taking the eigenvalue decomposition (EVD) of $\mathbf{A} = \mathbf{P}\mathbf{D}\mathbf{P}^\top$, where $\mathbf{P}$ is an orthogonal matrix (i.e., such that $\mathbf{P}^\top\mathbf{P} = \mathbf{P}\mathbf{P}^\top = \mathbf{I}_N$, with $\mathbf{I}_N$ again being the identity matrix) and $\mathbf{D}$ a diagonal matrix of eigenvalues (EVs), $d_1, \ldots, d_N$, we can premultiply $\mathbf{Y}^*$ in Eq. 2 by $\mathbf{P}^\top$ to obtain a sparse formulation of the model. Specifically, we have that:

$$\text{vec}\left(\mathbf{P}^\top\mathbf{Y}^*\right) = \left(\mathbf{I}_T \otimes \mathbf{P}^\top\right)\text{vec}\left(\mathbf{Y}^*\right) \sim \mathcal{N}\left(\left(\mathbf{I}_T \otimes \mathbf{P}^\top\right)\mathbf{X}^*\boldsymbol{\beta}, \left(\mathbf{I}_T \otimes \mathbf{P}^\top\right)\mathbf{V}^*\left(\mathbf{I}_T \otimes \mathbf{P}\right)\right), \text{ where} \tag{3}$$

$$\left(\mathbf{I}_T \otimes \mathbf{P}^\top\right)\mathbf{V}^*\left(\mathbf{I}_T \otimes \mathbf{P}\right) = \mathbf{V}_G \otimes \left(\mathbf{P}^\top\mathbf{A}\mathbf{P}\right) + \mathbf{V}_E \otimes \left(\mathbf{P}^\top\mathbf{P}\right) \tag{4}$$

$$= \mathbf{V}_G \otimes \left(\mathbf{P}^\top\mathbf{P}\mathbf{D}\mathbf{P}^\top\mathbf{P}\right) + \mathbf{V}_E \otimes \left(\mathbf{P}^\top\mathbf{P}\right) \tag{5}$$

$$= \mathbf{V}_G \otimes \mathbf{D} + \mathbf{V}_E \otimes \mathbf{I}_N. \tag{6}$$

The distribution of $\text{vec}\left(\mathbf{P}^\top\mathbf{Y}^*\right)$ can now be written more compactly as:

$$\text{vec}\left(\mathbf{P}^\top\mathbf{Y}^*\right) \sim \mathcal{N}\left(\left(\mathbf{I}_T \otimes \mathbf{P}^\top\right)\mathbf{X}^*\boldsymbol{\beta}, \mathbf{V}_G \otimes \mathbf{D} + \mathbf{V}_E \otimes \mathbf{I}_N\right). \tag{7}$$

The covariance matrix of this model is highly sparse, but this sparsity is spread over many rows and columns. However, by pre-multiplying the left-hand side of our model by the appropriate commutation matrix, $\mathbf{K}^{(N,T)}$ (i.e., re-ordering observations such that we order by individuals first, and then by traits, rather than the other way around), we can obtain a model where the resulting covariance matrix $\mathbf{V}$ is block-diagonal. That is, based on Eq. 7, we can now write $\mathbf{y} = \mathbf{K}^{(N,T)}\text{vec}\left(\mathbf{P}^\top\mathbf{Y}^*\right) = \text{vec}\left(\mathbf{Y}^{*\top}\mathbf{P}\right) = \text{vec}\left(\mathbf{Y}\right)$ as follows:

$$\mathbf{y} \sim \mathcal{N}\left(\widetilde{\mathbf{X}}\boldsymbol{\beta}, \mathbf{V}\right), \text{ where } \mathbf{V} = \mathbf{D} \otimes \mathbf{V}_G + \mathbf{I}_N \otimes \mathbf{V}_E \text{ and } \widetilde{\mathbf{X}} = \begin{pmatrix} \mathbf{X}_1 \\ \mathbf{X}_2 \\ \vdots \\ \mathbf{X}_N \end{pmatrix}, \tag{8}$$

where

$$\mathbf{X}_j = \begin{pmatrix} \mathbf{x}_{j1}^\top & & \mathbf{0} \\ & \ddots & \\ \mathbf{0} & & \mathbf{x}_{jT}^\top \end{pmatrix}, \tag{9}$$

is the $T \times K$ matrix of fixed-effect covariates across traits for observation $j = 1, \ldots, N$ after the linear transformation by the eigenvectors from the GRM. Here, $K$ is the total number of covariates across traits (i.e., $K = \sum_{t=1}^{T} K_t$). More specifically, vector $\mathbf{x}_{jt}^\top$ equals the $j$-th row from the $N \times K_t$ matrices $\mathbf{P}^\top\mathbf{X}_t^*$, and the vectors of zeros are conformable to the dimensions of $\mathbf{x}_{jt}$. In turn, $\mathbf{X}_t^*$ is the $N \times K_t$ matrix of fixed-effect covariates for trait $t$.

Notice that several asterisks have been dropped. That is, we here define our $T \times N$ matrix of outcomes $\mathbf{Y}$ as ${\mathbf{Y}^*}^\top \mathbf{P}$, and our grand matrix of fixed-effect covariates, $\widetilde{\mathbf{X}}$, and its constituents, $\mathbf{X}_j$ for $j = 1, \ldots, N$, and our covariance matrix, $\mathbf{V}$, as outlined above. These definitions boil down to pre-multiplication of conformable matrices by the transposed matrix of eigenvectors of the GRM, and re-ordering of rows of vectorised matrices in accordance with the commutation matrix. These definitions of $\mathbf{Y}$ and $\mathbf{X}_j$ for $j = 1, \ldots, N$ enable us to reduce the computational complexity of our model relatively easily. Notice that $\mathbf{V}$ can be written as follows:

$$\mathbf{V} = \begin{pmatrix} \mathbf{V}_1 & & \mathbf{0} \\ & \ddots & \\ \mathbf{0} & & \mathbf{V}_N \end{pmatrix}, \text{ where} \tag{10}$$

$$\mathbf{V}_j = d_j \mathbf{V}_G + \mathbf{V}_E \text{ for } j = 1, \ldots, N, \tag{11}$$

where $\mathbf{V}_j$ is a $T \times T$ symmetric positive (semi)definite matrix. Thus, $\mathbf{V}$ is now a block-diagonal matrix in which each block on the diagonal is a $T \times T$ matrix. As such, $\mathbf{V}$ is nothing more than a linear combination of $\mathbf{V}_G$ and $\mathbf{V}_E$ with weights based on the consecutive eigenvalues of the GRM. This matrix has sufficient structure to permit significant computational gains.

The combination of (i) $d_j \geq 0 \, \forall \, j$, (ii) $\mathbf{V}_G$ being at least positive semidefinite, and (iii) $\mathbf{V}_E$ being positive definite, forms a set of sufficient conditions for $\mathbf{V}_j$ to be positive definite $\forall \, j$, and, in turn, for $\mathbf{V}$ to be positive definite. As the GRM is at least positive semidefinte by definition, the first condition holds for sure. Secondly, by choosing an appropriate parametrisation of our model, we ensure $\mathbf{V}_G$ is always at least positive semidefinite. Finally, by again choosing an appropriate parametrisation, we also ensure $\mathbf{V}_E$ is always at least positive semidefinite. In further derivations, we make the stronger assumption that $\mathbf{V}_E$ is always positive definite (thus, by virtue of the set of sufficient conditions, ensuring $\mathbf{V}$ is always positive definite). Although this assumption may fail to hold in certain empirical applications (e.g., perfectly multicollinear traits as input for an MGREML analysis), it is safe to make this assumption for the derivations. The situation is in fact quite analogous to derivations for, e.g., the ordinary least squares (OLS) estimator, where one has the assumption that regressors (i.e., the explanatory variables) are not perfectly multicollinear although there is nothing preventing such collinearity occurring in empirical data. As in OLS regression with perfect multicollinearity of regressors, MGREML will fail to find a solution because of rank deficiency in matrix $\mathbf{V}_E$.

#### Step 3. The log-likelihood function, its gradient, and the information matrix

**Log-likelihood function.** The generic log-likelihood of a mixed linear model for $n \times 1$ vector $\mathbf{y}$ (in our case $n = NT$), as a function of parameters in $\boldsymbol{\theta}$, with fixed-effect covariates $\widetilde{\mathbf{X}}$, and variance matrix $\mathbf{V}$, is given by

$$\log l(\boldsymbol{\theta}) = -\frac{1}{2}\left(n\log(2\pi) + \log|\mathbf{V}| + \log\left|\widetilde{\mathbf{X}}^\top \mathbf{V}^{-1}\widetilde{\mathbf{X}}\right| - \log\left|\widetilde{\mathbf{X}}^\top \widetilde{\mathbf{X}}\right| + \mathbf{y}^\top \mathbf{M}\mathbf{y}\right), \text{ where} \tag{12}$$

$$\mathbf{M} = \mathbf{V}^{-1} - \mathbf{V}^{-1}\widetilde{\mathbf{X}}\left(\widetilde{\mathbf{X}}^\top \mathbf{V}^{-1}\widetilde{\mathbf{X}}\right)^{-1}\widetilde{\mathbf{X}}^\top \mathbf{V}^{-1}. \tag{13}$$

Importantly, $\mathbf{V}$ is a function of $\boldsymbol{\theta}$. Thus, we aim to find the $\boldsymbol{\theta}$ that sets the matrix $\mathbf{V}$ such that it maximises the log-likelihood of the observed data under the assumed distribution. Although this model incorporates fixed-effects $\widetilde{\mathbf{X}}$, to correct for possible confounding effects, it is not concerned with estimating those fixed effects. The primary aim of the model is to estimate the parameters $\boldsymbol{\theta}$ that make the variance matrix fitting the data as well as possible. Nevertheless, one can still use, e.g., the generalised least squares (GLS) estimator to obtain estimates of the fixed effects, because the model yields estimates of $\widehat{\boldsymbol{\theta}}$ and $\widehat{\mathbf{V}}$. That is, one can obtain the fixed-effects estimates by:

$$\mathbf{b} = \left(\widetilde{\mathbf{X}}^\top \widehat{\mathbf{V}}^{-1}\widetilde{\mathbf{X}}\right)^{-1}\widetilde{\mathbf{X}}^\top \widehat{\mathbf{V}}^{-1}\mathbf{y}. \tag{14}$$

This option is also implemented in the `GCTA` software package (Yang et al., 2011). The generic log-likelihood in Eq. 12 is also described by Yang et al. (2011) and is, in turn, based on a broad literature, including the works of Harville (1977), Casella and Searle (1985), and Searle et al. (1992). This log-likelihood is such that the presence of fixed-effect covariates does not bias the estimation of variance components, something that typically occurs when applying classical maximum likelihood estimation to such problems. For the simple case without covariates, it is clear that in our case, without canonical transformation and commutation, the matrix $\mathbf{V}$ is a full $(NT) \times (NT)$ matrix. Thus, under this naïve approach, even computing the log-likelihood requires the calculation of the eigenvalues of $\mathbf{V}$ (to calculate $\log|\mathbf{V}|$ in a numerically stable manner). These eigenvalues can be computed in $O\left(N^3T^3\right)$ time when using standard algorithms. As such, the complexity can become prohibitively large when $N$ and $T$ increase.

**Restricted maximum likelihood.** In order to estimate $\mathbf{V}_G$ and $\mathbf{V}_E$, we need to find the parameters that maximise the log-likelihood function in Eq. 12. This approach is known as restricted maximum likelihood

(REML). We are able to perform REML estimation efficiently by using highly efficient expressions for the log-likelihood and the gradient that we derive in subsequent sections. These expressions allow for rapid application of line-search methods and a BFGS algorithm. In addition, once our estimates have converged, we will use an efficient expression for the so-called average information (AI) matrix. This expression will also be derived in subsequent sections. Given we can easily calculate the AI matrix, obtaining the covariance matrix of $\widehat{\boldsymbol{\theta}}$ is straightforward. In turn, we can use covariance matrix $\widehat{\boldsymbol{\theta}}$ in conjunction with a delta method to compute the standard errors of our estimates of genetic correlations and heritabilities. Derivations for this delta method are also provided later on in this section.

**Gradient of the log-likelihood.** For a given parameter $\theta_1$ in the set of parameters $\boldsymbol{\theta}$ (using index 1 without loss of generality), the gradient of the log-likelihood, in accordance with Yang et al. (2011), is given by:

$$g_1 = \frac{1}{2}\mathbf{y}^\top \mathbf{M}\frac{\partial \mathbf{V}}{\partial \theta_1}\mathbf{M}\mathbf{y} - \frac{1}{2}\text{tr}\left(\mathbf{M}\frac{\partial \mathbf{V}}{\partial \theta_1}\right). \tag{15}$$

As we will show later on, we are able to compute $NT \times 1$ vector $\mathbf{r} = \mathbf{M}\mathbf{y}$ in $O\left(NT\right)$ time in case covariates are absent from our model and $O\left(NT^2\right)$ time in case there is a fairly limited number of covariates. Using the expression in Eq. 13 for $\mathbf{M}$, we can rewrite the gradient as follows:

$$g_1 = \frac{1}{2}\mathbf{r}^\top \frac{\partial \mathbf{V}}{\partial \theta_1}\mathbf{r} - \frac{1}{2}\text{tr}\left(\mathbf{V}^{-1}\frac{\partial \mathbf{V}}{\partial \theta_1}\right) + \frac{1}{2}\text{tr}\left(\mathbf{V}^{-1}\widetilde{\mathbf{X}}\left(\widetilde{\mathbf{X}}^\top \mathbf{V}^{-1}\widetilde{\mathbf{X}}\right)^{-1}\widetilde{\mathbf{X}}^\top \mathbf{V}^{-1}\frac{\partial \mathbf{V}}{\partial \theta_1}\right). \tag{16}$$

In our case, $\mathbf{V}$ is a block-diagonal matrix and therefore its inverse and its derivatives (both first- and second-order partial derivatives) also have a block-diagonal structure. These block-diagonal matrices have equally sized blocks. It holds that for two block-diagonal matrix with equally-sized blocks, $\mathbf{B}$ and $\mathbf{C}$, that the block diagonal matrix resulting from their product has blocks that are given by the products of the corresponding blocks in the two matrices. That is, block $h$ in $\mathbf{BC}$ is the product of block $h$ in $\mathbf{B}$ times block $h$ in $\mathbf{C}$. Also, note that $\text{tr}\left(AB\right) = \text{tr}\left(BA\right)$ and that the trace of a block-diagonal matrix can be written as the sum of traces of those blocks. These insights can be used to further rewrite the gradient as follows:

$$g_1 = \frac{1}{2}\left[\left(\sum_{j=1}^{N} \mathbf{r}_j^\top \frac{\partial \mathbf{V}_j}{\partial \theta_1}\mathbf{r}_j\right) - \sum_{j=1}^{N} \operatorname{tr}\left(\mathbf{V}_j^{-1}\frac{\partial \mathbf{V}_j}{\partial \theta_1}\right) + \operatorname{tr}\left(\mathbf{Z}^{-1}\left(\sum_{j=1}^{N} \mathbf{X}_j^\top \mathbf{V}_j^{-1}\frac{\partial \mathbf{V}_j}{\partial \theta_1}\mathbf{V}_j^{-1}\mathbf{X}_j\right)\right)\right], \text{ where} \tag{17}$$

$$\mathbf{Z} = \widetilde{\mathbf{X}}^\top \mathbf{V}^{-1}\widetilde{\mathbf{X}} = \sum_{j=1}^{N} \mathbf{X}_j^\top \mathbf{V}_j^{-1}\mathbf{X}_j. \tag{18}$$

Although Equations 17 and 18 may seem more involved than Eq. 16, the gradient has now been decomposed into a contribution per observation. This decomposition is at the core of making time complexity for the gradient linear in the number of observations. For reasons of brevity, we refer to $\mathbf{Z}$ in Eq. 18 as the GLS precision matrix.

**Information matrix of the log-likelihood.** Without loss of generality, let $\overline{\mathcal{I}}_{12}$ denote the element of the average-information matrix corresponding to parameters $\theta_1$ and $\theta_2$. This element is defined as:

$$\overline{\mathcal{I}}_{12} = \frac{1}{2}\left(\mathcal{I}_{12} + \mathbb{E}\left[\mathcal{I}_{12}\right]\right), \tag{19}$$

where $\mathcal{I}_{12}$ (resp. $\mathbb{E}\left[\mathcal{I}_{12}\right]$) denotes the corresponding element from the observed (expected) information matrix. Based on the work by Gilmour et al. (1995), we know that

$$\mathcal{I}_{12} = \frac{1}{2}\operatorname{tr}\left(\mathbf{M}\frac{\partial^2 \mathbf{V}}{\partial \theta_1 \partial \theta_2}\right) - \frac{1}{2}\operatorname{tr}\left(\mathbf{M}\frac{\partial \mathbf{V}}{\partial \theta_1}\mathbf{M}\frac{\partial \mathbf{V}}{\theta_2}\right) + \mathbf{y}^\top \mathbf{M}\frac{\partial \mathbf{V}}{\partial \theta_1}\mathbf{M}\frac{\partial \mathbf{V}}{\partial \theta_2}\mathbf{M}\mathbf{y} - \frac{1}{2}\mathbf{y}^\top \mathbf{M}\frac{\partial^2 \mathbf{V}}{\partial \theta_1 \partial \theta_2}\mathbf{M}\mathbf{y} \text{ and} \tag{20}$$

$$\mathbb{E}\left[\mathcal{I}_{12}\right] = \frac{1}{2}\operatorname{tr}\left(\mathbf{M}\frac{\partial \mathbf{V}}{\partial \theta_1}\mathbf{M}\frac{\partial \mathbf{V}}{\theta_2}\right). \tag{21}$$

By combining terms, we obtain that an element of the average information matrix can be written as:

$$\overline{\mathcal{I}}_{12} = \frac{1}{4}\operatorname{tr}\left(\mathbf{M}\frac{\partial^2 \mathbf{V}}{\partial \theta_1 \partial \theta_2}\right) - \frac{1}{4}\mathbf{y}^\top \mathbf{M}\frac{\partial^2 \mathbf{V}}{\partial \theta_1 \partial \theta_2}\mathbf{M}\mathbf{y} + \frac{1}{2}\mathbf{y}^\top \mathbf{M}\frac{\partial \mathbf{V}}{\partial \theta_1}\mathbf{M}\frac{\partial \mathbf{V}}{\partial \theta_2}\mathbf{M}\mathbf{y}. \tag{22}$$

As the computational complexity of this expression is high, Gilmour et al. (1995) note that $\mathbf{y}^\top \mathbf{M}\frac{\partial^2 \mathbf{V}}{\partial \theta_1 \partial \theta_2}\mathbf{M}\mathbf{y}$ can be approximated by its expectation $\operatorname{tr}\left(\mathbf{M}\frac{\partial^2 \mathbf{V}}{\partial \theta_1 \partial \theta_2}\right)$ when these second-order derivatives are non-zero. This precedent is followed by others in the AI-REML literature (e.g., Yang et al. (2011)). Using this approximation, the expression for an element of the AI matrix reduces to:

$$\overline{\mathcal{I}}_{12} = \frac{1}{2}\mathbf{y}^\top \mathbf{M}\frac{\partial \mathbf{V}}{\partial \theta_1}\mathbf{M}\frac{\partial \mathbf{V}}{\partial \theta_2}\mathbf{M}\mathbf{y}. \tag{23}$$

Thus, given the efficient expression of $\mathbf{My}$ and the block-diagonal structure of the problem, an element of the AI matrix can be described as:

$$\overline{\mathcal{I}}_{12} = \frac{1}{2}\left[\mathbf{r}^\top \frac{\partial \mathbf{V}}{\partial \theta_1}\mathbf{V}^{-1}\frac{\partial \mathbf{V}}{\partial \theta_2}\mathbf{r} - \mathbf{r}^\top \frac{\partial \mathbf{V}}{\partial \theta_1}\mathbf{V}^{-1}\widetilde{\mathbf{X}}\left(\widetilde{\mathbf{X}}^\top \mathbf{V}^{-1}\widetilde{\mathbf{X}}\right)^{-1}\widetilde{\mathbf{X}}^\top \mathbf{V}^{-1}\frac{\partial \mathbf{V}}{\partial \theta_2}\mathbf{r}\right] \tag{24}$$

$$= \frac{1}{2}\left[\left(\sum_{j=1}^{N}\mathbf{r}_j^\top \frac{\partial \mathbf{V}_j}{\partial \theta_1}\mathbf{V}_j^{-1}\frac{\partial \mathbf{V}_j}{\partial \theta_2}\mathbf{r}_j\right) - \left(\sum_{j=1}^{N}\mathbf{r}_j^\top \frac{\partial \mathbf{V}_j}{\partial \theta_1}\mathbf{V}_j^{-1}\mathbf{X}_j\right)\mathbf{Z}^{-1}\left(\sum_{j=1}^{N}\mathbf{X}_j^\top \mathbf{V}_j^{-1}\frac{\partial \mathbf{V}_j}{\partial \theta_2}\mathbf{r}\right)\right]. \tag{25}$$

#### Step 4. Tractable notation for different covariates across traits

In most derivations that will follow, we assume there is one set of $k$ fixed-effect covariates that applies to all traits. In that case, we allow for $Tk$ fixed effects in total. This assumption simplifies derivations considerably. However, here we show that the resulting expressions can easily be generalised to cases in which different sets of covariates apply to different traits, as well as to the case of having no covariates at all. The biggest advantage of considering one set of covariates that applies to all traits, is that we can write down the matrix of fixed-effect covariates for a particular observation $j$ as a Kronecker product.

Let $\mathcal{K}_t$ denote the set of covariates that applies to trait $t$ for $t = 1, \ldots, T$. We assume that $\mathcal{K} = \bigcup_{t=1}^{T} \mathcal{K}_t$ is the complete set of covariates, which, without loss of generality, for now applies to all traits. That is, $\mathcal{K} = \mathcal{K}_t \, \forall \, t$, with $k = ||\mathcal{K}||$ denoting the total number of unique covariates. Now, letting

$$\mathbf{X} = \mathbf{P}^\top \mathbf{X}^*, \tag{26}$$

with $\mathbf{X}^*$ and $\mathbf{X}$ denoting the $N \times k$ matrices of fixed-effect covariates (the latter after the canonical transformation) with observations in the rows and covariates in the columns, we obtain that the $T \times Tk$ matrices $\mathbf{X}_j$ for observations $j = 1, \ldots, N$, as defined in Eq. 9, can be written as:

$$\mathbf{X}_j = \mathbf{I}_T \otimes \mathbf{x}_j^\top. \tag{27}$$

Here, the $1 \times k$ vector $\mathbf{x}_j^\top$ is the $j$-th row from $\mathbf{X}$ for $j = 1, \ldots, N$. With this notation for the same set of covariates applying to all traits, we can easily generalise to the case of different covariates for different traits. Now, assume there is a binary matrix $\mathbf{S}_t$ for each trait, such that $\mathbf{S}_t$ is a $K_t \times k$ matrix ($K_t \leq k$), with $\mathbf{S}_t\mathbf{S}_t^\top = \mathbf{I}_{K_t}$. When covariate $j$ (from the full set of covariates, $\mathcal{K}$) is the $i$-th covariate for trait $t$, then element $i, j$ in matrix $\mathbf{S}_t$ is equal to one. Otherwise, that element equals zero. Note that in case all covariates apply to a given trait, $\mathbf{S}_t$ is simply equal to $\mathbf{I}_k$. Effectively, $\mathbf{S}_t\mathbf{A}$ yields a submatrix of matrix $\mathbf{A}$, comprising only $K_t$ unique rows from the $k$ rows in $\mathbf{A}$. If matrix $\mathbf{A}$ is square, $\mathbf{S}_t\mathbf{A}\mathbf{S}_t^\top$ yields the $K_t \times K_t$ submatrix of $\mathbf{A}$ in which the appropriate rows and columns from $\mathbf{A}$ are selected.

Now, we can construct a grand block-diagonal matrix

$$\mathbf{S} = \begin{pmatrix} \mathbf{S}_1 & & \mathbf{0} \\ & \ddots & \\ \mathbf{0} & & \mathbf{S}_T \end{pmatrix}, \tag{28}$$

such that $\mathbf{S}\mathbf{S}^\top = \mathbf{I}_K$, where $K = \sum_{t=1}^{T} K_t$. Using these definitions, the GLS precision matrix can be written as:

$$\mathbf{Z} = \mathbf{S}\left[\sum_{j=1}^{N} (\mathbf{I}_T \otimes \mathbf{x}_j)\, \mathbf{V}_j^{-1} \left(\mathbf{I}_T \otimes \mathbf{x}_j^\top\right)\right] \mathbf{S}^\top, \tag{29}$$

and the $T \times K$ matrix of fixed-effect covariates for observation $j$ as

$$\mathbf{X}_j = \left(\mathbf{I}_T \otimes \mathbf{x}_j^\top\right) \mathbf{S}^\top. \tag{30}$$

Now, we have derived expressions for two scenarios. In the first scenario, there is a set of $k$ fixed-effect covariates that applies to all traits. In the second scenario, there are different covariates for different traits. The derived expressions allow to obtain efficient expressions for the log-likelihood, gradient, and the information matrix.

#### Step 5. Calculating the log-likelihood efficiently

To calculate the log-likelihood rapidly, we need efficient expressions for $\mathbf{V}^{-1}$, $\log|\mathbf{V}|$, $\log|\mathbf{Z}|$, and $\mathbf{y}^\top\mathbf{M}\mathbf{y} = \mathbf{y}^\top\mathbf{r}$ with $\mathbf{r}$ referring to the rescaled GLS residuals or just the residuals.

**Decomposition of the variance matrix.** The full variance matrix, $\mathbf{V}$, in our model in Eq. 8, has a block-diagonal structure as illustrated in Eq. 10 and Eq. 11. Hence, the inverse of $\mathbf{V}$ is a block-diagonal matrix too, with $\mathbf{V}_j^{-1}$ for $j = 1, \ldots, N$ as blocks. Also, the determinant of $\mathbf{V}$ is the product of the determinants of $\mathbf{V}_j$ for $j = 1, \ldots, N$. Let the EVD of $\mathbf{V}_E$ be given by $\mathbf{Q}\mathbf{\Phi}\mathbf{Q}^\top$, with $\mathbf{Q}\mathbf{Q}^\top = \mathbf{Q}^\top\mathbf{Q} = \mathbf{I}_T$. We then can rewrite $\mathbf{V}_j$ as follows:

$$\mathbf{V}_j = \mathbf{Q}\boldsymbol{\Phi}^{\frac{1}{2}} \left( d_j \widetilde{\mathbf{V}}_G + \mathbf{I}_T \right) \boldsymbol{\Phi}^{\frac{1}{2}} \mathbf{Q}^\top, \text{ where} \tag{31}$$

$$\widetilde{\mathbf{V}}_G = \boldsymbol{\Phi}^{-\frac{1}{2}} \mathbf{Q}^\top \mathbf{V}_G \mathbf{Q} \boldsymbol{\Phi}^{-\frac{1}{2}}. \tag{32}$$

As our model assumes a positive definite $\mathbf{V}_E$, $\boldsymbol{\Phi}$ is a diagonal matrix with positive diagonal entries and $\boldsymbol{\Phi}^{-\frac{1}{2}}$ is defined in terms of real numbers. Moreover, because $\mathbf{V}_G$ is at least positive semi-definite, so is $\widetilde{\mathbf{V}}_G$ (this can be shown easily using quadratic forms with respect to $\widetilde{\mathbf{V}}_G$, which can be rewritten as quadratic forms with respect to $\mathbf{V}_G$). Let $\mathbf{L}\boldsymbol{\Lambda}\mathbf{L}^\top$ denote the EVD of $\widetilde{\mathbf{V}}_G$, with $\mathbf{L}\mathbf{L}^\top = \mathbf{L}^\top\mathbf{L} = \mathbf{I}_T$. Now we can rewrite $\mathbf{V}_j$ as:

$$\mathbf{V}_j = \mathbf{Q}\boldsymbol{\Phi}^{\frac{1}{2}} \mathbf{L} \left( d_j \boldsymbol{\Lambda} + \mathbf{I}_T \right) \mathbf{L}^\top \boldsymbol{\Phi}^{\frac{1}{2}} \mathbf{Q}^\top. \tag{33}$$

Hence, we have that

$$\mathbf{V}_j^{-1} = \mathbf{F}\mathbf{D}_j^* \mathbf{F}^\top, \text{ where} \tag{34}$$

$$\mathbf{D}_j^* = \left( d_j \boldsymbol{\Lambda} + \mathbf{I}_T \right)^{-1} \text{ and} \tag{35}$$

$$\mathbf{F} = \mathbf{Q}\boldsymbol{\Phi}^{-\frac{1}{2}} \mathbf{L}. \tag{36}$$

Note that although $\mathbf{F}$ is square, it is neither necessarily a symmetric nor necessarily an orthogonal matrix. By means of this expression for $\mathbf{V}_j$, we are able to invert the $(NT) \times (NT)$ covariance matrix $\mathbf{V}$ at the price of performing $N$ matrix multiplications, each in $O\left(T^3\right)$ time. That is, multiplying $T \times T$ matrix $\mathbf{F}\mathbf{D}_j^*$ and $T \times T$ matrix $\mathbf{F}^\top$. Thus, we have reduced time for inverting $\mathbf{V}$ from $O\left(N^3 T^3\right)$ to $O\left(NT^3\right)$.

In our case, a further reduction is possible, because we can exploit the fact that $\mathbf{V}_j^{-1}$ only varies from observation to observation in terms of the diagonal matrix $\mathbf{D}_j^*$. This allows us to derive fast expressions for the log-likelihood and gradient, because it is not needed to explicitly calculate $\mathbf{V}_j^{-1}$. As we will show, this implies we can calculate the log-likelihood and gradient in $O\left(NT^2\right)$ time.

**Efficient determinant of the variance matrix.** For two conformable square matrices, $\mathbf{B}$ and $\mathbf{C}$, the following identity holds:

$$|\mathbf{AB}| = |\mathbf{A}|\,|\mathbf{B}|\,. \tag{37}$$

Moreover, the determinant of a block-diagonal matrix can be written as the product of the determinants of the blocks. Therefore, the determinant of $\mathbf{V}$ can be rewritten as:

$$\begin{aligned}
|\mathbf{V}| &= \prod_{j=1}^{N} |\mathbf{V}_j| && (38)\\
&= \prod_{j=1}^{N} \left( \left| \mathbf{Q}\boldsymbol{\Phi}^{\frac{1}{2}}\mathbf{L}\left(d_j\boldsymbol{\Lambda} + \mathbf{I}_T\right)\mathbf{L}^\top\boldsymbol{\Phi}^{\frac{1}{2}}\mathbf{Q}^\top \right| \right) && (39)\\
&= \prod_{j=1}^{N} \left( |\mathbf{Q}| \left|\boldsymbol{\Phi}^{\frac{1}{2}}\right| |\mathbf{L}| \left|d_j\boldsymbol{\Lambda} + \mathbf{I}_T\right| \left|\mathbf{L}^\top\right| \left|\boldsymbol{\Phi}^{\frac{1}{2}}\right| \left|\mathbf{Q}^\top\right| \right) && (40)\\
&= \prod_{j=1}^{N} \left( \left|d_j\boldsymbol{\Lambda} + \mathbf{I}_T\right| \left|\mathbf{L}^\top\right| |\mathbf{L}| \left|\boldsymbol{\Phi}^{\frac{1}{2}}\right| \left|\boldsymbol{\Phi}^{\frac{1}{2}}\right| \left|\mathbf{Q}^\top\right| |\mathbf{Q}| \right) && (41)\\
&= \prod_{j=1}^{N} \left( \left|d_j\boldsymbol{\Lambda} + \mathbf{I}_T\right| \left|\mathbf{L}^\top\mathbf{L}\right| |\boldsymbol{\Phi}| \left|\mathbf{Q}^\top\mathbf{Q}\right| \right) && (42)\\
&= \prod_{j=1}^{N} \left( \prod_{t=1}^{T} \left(d_j\lambda_t + 1\right) |\mathbf{I}_T| \prod_{t=1}^{T} \phi_t\, |\mathbf{I}_T| \right) && (43)\\
&= \prod_{j=1}^{N} \left( \prod_{t=1}^{T} \left(d_j\lambda_t + 1\right) \prod_{t=1}^{T} \phi_t \right), && (44)
\end{aligned}$$

where $\lambda_t$ is the $t$-th diagonal entry of $\boldsymbol{\Lambda}$ and where $\phi_t$ is defined analogously with respect to $\boldsymbol{\Phi}$. Hence, the log-determinant of $\mathbf{V}$ is given by

$$\log|\mathbf{V}| = N\sum_{t=1}^{T}\log\left(\phi_t\right) + \sum_{j=1}^{N}\sum_{t=1}^{T}\log\left(d_j\lambda_t + 1\right), \tag{45}$$

where $\lambda_t$ are the EVs of $\widetilde{\mathbf{V}}_G$ in $\boldsymbol{\Lambda}$, $d_j$ are the EVs of the GRM, and $\phi_t$ the EVs of $\mathbf{V}_E$ in $\boldsymbol{\Phi}$. Calculating this log-determinant now takes $O\left(NT\right)$ time, given we have precomputed the EVD of the GRM, $\widetilde{\mathbf{V}}_G$, and $\mathbf{V}_E$.

**GLS precision matrix, its inverse, and determinant.** Here we start again with the assumption of identical covariates across traits (i.e., $\mathbf{S} = \mathbf{I}_{Tk}$). Using our expression for $\mathbf{V}_j^{-1}$ in Eq. 34 and properties of the Kronecker product, the GLS precision matrix can be rewritten as:

$$\mathbf{Z} = \sum_{j=1}^{N} \mathbf{X}_j^\top \mathbf{V}_j^{-1} \mathbf{X}_j \tag{46}$$

$$= \sum_{j=1}^{N} \left(\mathbf{I}_T \otimes \mathbf{x}_j\right) \left(\mathbf{V}_j^{-1} \otimes \mathbf{I}_1\right) \left(\mathbf{I}_T \otimes \mathbf{x}_j^\top\right) \tag{47}$$

$$= \sum_{j=1}^{N} \left(\mathbf{V}_j^{-1} \otimes \mathbf{x}_j \mathbf{x}_j^\top\right) \tag{48}$$

$$= \sum_{j=1}^{N} \left(\left(\mathbf{F}\mathbf{D}_j^* \mathbf{F}^\top\right) \otimes \mathbf{x}_j \mathbf{x}_j^\top\right) \tag{49}$$

$$= \sum_{j=1}^{N} \left(\mathbf{F} \otimes \mathbf{I}_k\right) \left(\mathbf{D}_j^* \otimes \mathbf{x}_j \mathbf{x}_j^\top\right) \left(\mathbf{F}^\top \otimes \mathbf{I}_k\right) \tag{50}$$

$$= \left(\mathbf{F} \otimes \mathbf{I}_k\right) \left[\sum_{j=1}^{N} \left(\mathbf{D}_j^* \otimes \mathbf{x}_j \mathbf{x}_j^\top\right)\right] \left(\mathbf{F}^\top \otimes \mathbf{I}_k\right) \tag{51}$$

$$= \left(\mathbf{F} \otimes \mathbf{I}_k\right) \begin{bmatrix} \mathbf{B}_1 & & \mathbf{0} \\ & \ddots & \\ \mathbf{0} & & \mathbf{B}_T \end{bmatrix} \left(\mathbf{F}^\top \otimes \mathbf{I}_k\right), \text{ where } \mathbf{B}_t = \sum_{j=1}^{N} \frac{1}{d_j \lambda_t + 1} \mathbf{x}_j \mathbf{x}_j^\top = \mathbf{X}^\top \mathbf{D}_t^\dagger \mathbf{X} \text{ and } \mathbf{D}_t^\dagger = \left(\mathbf{D}\lambda_t + \mathbf{I}_N\right)^{-1}. \tag{52}$$

Recall here that $\mathbf{X}$ is the $N \times k$ matrix of covariates, after the canonical transformation. Given that $k = O(1)$, each matrix $\mathbf{B}_t$ can be computed in $O(N)$ time. We can, therefore, calculate $\mathbf{B}_t$ across all traits in $O(NT)$ time. The outer pre- and post-multiplications of the resulting block-diagonal matrix are trivial. Thus, overall, $\mathbf{Z}$ can be calculated in $O(NT)$ time. Notice here how we have avoided the need to make $\mathbf{V}_j^{-1}$ explicit, thus avoiding a computation in $O\left(NT^3\right)$ time. We will pursue similar strategies for the remaining terms in the log-likelihood, gradient, and AI matrix.

The inverse of the precision matrix can be obtained efficiently, as

$$\mathbf{Z}^{-1} = \left(\left(\mathbf{F}^{-1}\right)^\top \otimes \mathbf{I}_k\right) \mathbf{J} \left(\mathbf{F}^{-1} \otimes \mathbf{I}_k\right), \text{ with} \tag{53}$$

$$\mathbf{J} = \begin{pmatrix} \mathbf{B}_1^{-1} & & \mathbf{0} \\ & \ddots & \\ \mathbf{0} & & \mathbf{B}_T^{-1} \end{pmatrix}. \tag{54}$$

Here, $\mathbf{F}$ is the $T \times T$ matrix which is fixed across observations and parameters and is. Therefore, it is computationally cheap to invert it (i.e., in $O\left(T^3\right)$ time). The blocks of the inner matrix can jointly be inverted in $O(T)$ time, assuming $k = O(1)$.

For the determinant of the precision matrix, we have that

$$
\begin{aligned}
|\mathbf{Z}| &= |\mathbf{F} \otimes \mathbf{I}_k| \prod_{t=1}^{T} |\mathbf{B}_t| \left|\mathbf{F}^\top \otimes \mathbf{I}_k\right| && (55)\\
&= \prod_{t=1}^{T} |\mathbf{B}_t| \, |\mathbf{F} \otimes \mathbf{I}_k| \left|\mathbf{F}^\top \otimes \mathbf{I}_k\right| && (56)\\
&= \prod_{t=1}^{T} |\mathbf{B}_t| \left|(\mathbf{F} \otimes \mathbf{I}_k)\left(\mathbf{F}^\top \otimes \mathbf{I}_k\right)\right| && (57)\\
&= \prod_{t=1}^{T} |\mathbf{B}_t| \left|\mathbf{F}\mathbf{F}^\top \otimes \mathbf{I}_k\right| && (58)\\
&= \prod_{t=1}^{T} |\mathbf{B}_t| \left|\mathbf{F}\mathbf{F}^\top\right|^k && (59)\\
&= \prod_{t=1}^{T} |\mathbf{B}_t| \left|\mathbf{Q}\mathbf{\Phi}^{-\frac{1}{2}}\mathbf{L}\mathbf{L}^\top\mathbf{\Phi}^{-\frac{1}{2}}\mathbf{Q}^\top\right|^k && (60)\\
&= \prod_{t=1}^{T} |\mathbf{B}_t| \left|\mathbf{Q}\mathbf{\Phi}^{-1}\mathbf{Q}^\top\right|^k && (61)\\
&= \prod_{t=1}^{T} |\mathbf{B}_t| \left(|\mathbf{Q}| \left|\mathbf{\Phi}^{-1}\right| \left|\mathbf{Q}^\top\right|\right)^k && (62)\\
&= \prod_{t=1}^{T} |\mathbf{B}_t| \left(\left|\mathbf{\Phi}^{-1}\right| |\mathbf{Q}| \left|\mathbf{Q}^\top\right|\right)^k && (63)\\
&= \prod_{t=1}^{T} |\mathbf{B}_t| \left(\left|\mathbf{\Phi}^{-1}\right| \left|\mathbf{Q}\mathbf{Q}^\top\right|\right)^k && (64)\\
&= \prod_{t=1}^{T} |\mathbf{B}_t| \, |\mathbf{\Phi}|^{-k}. && (65)
\end{aligned}
$$

Here, $\mathbf{\Phi}$ are the eigenvalues of $\mathbf{V}_E$. Therefore, the log-determinant of $\mathbf{Z}$ is:

$$
\log|\mathbf{Z}| = \sum_{t=1}^{T} \left(\log|\mathbf{B}_t| - k\log(\phi_t)\right), \tag{66}
$$

where $\mathbf{B}_t = \mathbf{X}^\top \mathbf{D}_t^\dagger \mathbf{X}$ and where $\phi_t$ denotes the $t$-th eigenvalue of $\mathbf{V}_E$. For numerical stability, we compute the eigenvalue decomposition of $\mathbf{B}_t$ for $t = 1, \ldots, T$ and compute $\log|\mathbf{B}_t|$ by taking the sum of the logarithm of the resulting eigenvalues. We use the same eigenvalue decompositions to set $\mathbf{B}_t^{-1}$.

In case we have different covariates across traits, our precision matrix is given by:

$$\mathbf{Z} = \left(\mathbf{S}\left(\mathbf{F} \otimes \mathbf{I}_k\right)\right) \begin{bmatrix} \mathbf{B}_1 & & \mathbf{0} \\ & \ddots & \\ \mathbf{0} & & \mathbf{B}_T \end{bmatrix} \left(\left(\mathbf{F}^\top \otimes \mathbf{I}_k\right) \mathbf{S}^\top\right). \tag{67}$$

Here, $\mathbf{S}$ is the binary $K \times Tk$ matrix as defined earlier. This equation implies $\mathbf{Z}$ can still be obtained in $O(NT)$ when the covariates differ by trait, assuming $k = O(1)$. In such a case, we compute the eigenvalue decomposition of $\mathbf{Z}$ as it is (which can still be done in $O\left(T^3\right)$ time), and use this decomposition to compute both $\log|\mathbf{Z}|$ and $\mathbf{Z}^{-1}$.

**GLS residuals.** Expressions for the GLS residuals and $\mathbf{y}^\top\mathbf{M}\mathbf{y}$ can now also be rewritten such that we can compute them in $O\left(NT^2\right)$ time. To see this, note that for the case of indentical covariates across traits, we can compute the $T \times 1$ vectors of rescaled GLS residuals, $\mathbf{r}_j$, the $Tk \times 1$ vector of GLS estimates, $\mathbf{b}$, and the rescaled $T \times 1$ phenotype vectors, $\widetilde{\mathbf{y}}_j$, for $j = 1, \ldots, N$ as follows:

$$\mathbf{M}\mathbf{y} = \begin{pmatrix} \mathbf{r}_1 \\ \vdots \\ \mathbf{r}_N \end{pmatrix}, \text{ where} \tag{68}$$

$$\mathbf{r}_j = \widetilde{\mathbf{y}}_j - \mathbf{F}\left(\mathbf{D}_j^*\mathbf{F}^\top\left(\left(\mathbf{I}_T \otimes \mathbf{x}_j^\top\right)\mathbf{b}\right)\right), \tag{69}$$

$$\mathbf{b} = \mathbf{Z}^{-1}\left(\sum_{j=1}^{N}\left(\mathbf{I}_T \otimes \mathbf{x}_j\right)\widetilde{\mathbf{y}}_j\right), \text{ and} \tag{70}$$

$$\widetilde{\mathbf{y}}_j = \left(\mathbf{F}\left(\mathbf{D}_j^*\left(\mathbf{F}^\top\mathbf{y}_j\right)\right)\right). \tag{71}$$

Now, let $\widetilde{\mathbf{Y}} = \left(\widetilde{\mathbf{y}}_1 \quad \ldots \quad \widetilde{\mathbf{y}}_N\right)$ denote the $T \times N$ matrix of $T \times 1$ vectors $\widetilde{\mathbf{y}}_j$ for $j = 1, \ldots, N$. Then, $\mathbf{b}$ can be written as:

$$\mathbf{b} = \mathbf{Z}^{-1}\left(\sum_{j=1}^{N}\left(\mathbf{I}_T \otimes \mathbf{x}_j\right)\left(\widetilde{\mathbf{y}}_j \otimes \mathbf{I}_1\right)\right) \tag{72}$$

$$= \mathbf{Z}^{-1}\left(\sum_{j=1}^{N}\widetilde{\mathbf{y}}_j \otimes \mathbf{x}_j\right) \tag{73}$$

$$= \mathbf{Z}^{-1}\text{vec}\left(\mathbf{X}^\top\widetilde{\mathbf{Y}}^\top\right). \tag{74}$$

Given these expressions, $\widetilde{\mathbf{Y}}$ can be calculated in $O\left(NT^2\right)$ time. Moreover, given that $k = O(1)$, $\mathbf{X}^\top \widetilde{\mathbf{Y}}^\top$ can be calculated in $O(NT)$ time, provided we have $\widetilde{\mathbf{Y}}$. Using $\widetilde{\mathbf{Y}}$, $\mathbf{b}$, and $\left(\mathbf{I}_T \otimes \mathbf{x}_j^\top\right)$, it is straightforward to construct the rescaled GLS residuals. The full set of residuals can therefore be obtained in $O\left(NT^2\right)$ time.

Let $\mathbf{R} = (\mathbf{r}_1 \quad \ldots \quad \mathbf{r}_N)$ denote the $T \times N$ matrix of rescaled GLS residuals. Then, we obtain that:

$$\mathbf{y}^\top \mathbf{M} \mathbf{y} = \boldsymbol{\iota}^\top \left(\mathbf{R} \circ \mathbf{Y}\right) \boldsymbol{\iota}. \tag{75}$$

Here, $\boldsymbol{\iota}^\top$ is a $1 \times T$ vector of ones and $\boldsymbol{\iota}$ a $N \times 1$ vector of ones, and '$\circ$' denotes the Hadamard or element-wise product. Recall that $\mathbf{Y}$ is the $T \times N$ matrix of phenotypes after post-multiplication by $\mathbf{P}$, the eigenvectors of the GRM. In case we have different covariates across traits, we only need to modify our expressions for $\mathbf{b}$ and $\mathbf{r}_j$. In this particular case, we have that:

$$\mathbf{r}_j = \widetilde{\mathbf{y}}_j - \mathbf{F}\left(\mathbf{D}_j^* \mathbf{F}^\top \left(\left(\mathbf{I}_T \otimes \mathbf{x}_j^\top\right) \mathbf{S}^\top \mathbf{b}\right)\right), \text{ where} \tag{76}$$

$$\mathbf{b} = \mathbf{Z}^{-1} \mathbf{S} \text{vec}\left(\mathbf{X}^\top \widetilde{\mathbf{Y}}^\top\right). \tag{77}$$

**Constant.** In case we are interested in the constant term in the log-likelihood function, we need an efficient expression for $\log\left|\widetilde{\mathbf{X}}^\top \widetilde{\mathbf{X}}\right|$. Under identical covariates across traits, we have that

$$\widetilde{\mathbf{X}}^\top \widetilde{\mathbf{X}} = \sum_{j=1}^{N} \left(\mathbf{I}_T \otimes \mathbf{x}_j\right) \left(\mathbf{I}_T \otimes \mathbf{x}_j^\top\right) \tag{78}$$

$$= \sum_{j=1}^{N} \left(\mathbf{I}_T \otimes \mathbf{x}_j \mathbf{x}_j^\top\right) \tag{79}$$

$$= \begin{pmatrix} \mathbf{X}^\top \mathbf{X} & & \mathbf{0} \\ & \ddots & \\ \mathbf{0} & & \mathbf{X}^\top \mathbf{X} \end{pmatrix}. \tag{80}$$

Here, $\mathbf{X}$ is the $N \times k$ matrix of covariates that is identical across all traits, after the canonical transformation. It therefore holds that:

$$\log\left|\widetilde{\mathbf{X}}^\top \widetilde{\mathbf{X}}\right| = T \log\left|\mathbf{X}^\top \mathbf{X}\right|, \tag{81}$$

where $\log\left|\mathbf{X}^{\top}\mathbf{X}\right|$ can be obtained in a numerically stable manner from the EVD of $\mathbf{X}^{\top}\mathbf{X}$. $\mathbf{X}^{\top}\mathbf{X}$ and its EVD can be obtained in $O(N)$ time, provided $k = O(1)$. Note that when there are different covariates across traits,

$$\log\left|\widetilde{\mathbf{X}}^{\top}\widetilde{\mathbf{X}}\right| = \sum_{t=1}^{T}\log\left|\mathbf{S}_t\mathbf{X}^{\top}\mathbf{X}\mathbf{S}_t^{\top}\right|. \tag{82}$$

This step involves taking submatrices of $\mathbf{X}^{\top}\mathbf{X}$ and computation of the determinants of those submatrices. When we assume $k = O(1)$, all these determinants can be obtained in $O(T)$ time in total. Here, the bottleneck is the calculation of $\mathbf{X}^{\top}\mathbf{X}$, which takes $O(N)$ time when $k = O(1)$. Thus, from a computational point of view, the calculation of $\log\left|\mathbf{X}^{\top}\mathbf{X}\right|$ is trivial for reasonable values of $N$ and $T$, and for $k = O(1)$.

**Overall complexity of the log-likelihood.** We can compute the MGREML log-likelihood in $O\left(NT^2\right)$ time, provided $k$ (i.e., the number of covariates in the union of the sets of covariates across traits) is $O(1)$. We need $O\left(NT^2\right)$ time for computing the log-likelihood for the three main cases, *viz.*, (i) having no covariates at all, (ii) having identical covariates across traits, and (iii) having only partial overlap between the sets of covariates across traits. However, we expect the first case to need the lowest runtime because calculating terms such as $\log|\mathbf{Z}|$ is not needed when there no covariates. The calculation will be slowest in case of non-identical covariates across traits, because of the additional steps that need to be taken.

#### Step 6. Parametrisation of the covariance matrices

Before we can derive efficient expressions for the gradient and AI matrix, we need to settle the parametrization of our model so that we can define the partial derivatives of $\mathbf{V}$ with respect to our parameters. To do so, we follow a simple factor model for both $\mathbf{V}_G$ and $\mathbf{V}_E$. That is:

$$\mathbf{V}_G = \mathbf{C}_G\mathbf{C}_G^{\top}, \tag{83}$$

$$\mathbf{V}_E = \mathbf{C}_E\mathbf{C}_E^{\top}, \tag{84}$$

where $\mathbf{C}_G$ has size $T \times F_G$ and $\mathbf{C}_E$ has size $T \times F_E$, with $F_G \leq T$ and (for identification purposes) $F_E = T$. Thus, we have at most as many genetic factors as we have traits, and we have as many environmental factors as we have traits. Further conditions of $\mathbf{C}_G$ and $\mathbf{C}_E$ matrices will be discussed later on. However, for present purposes these definitions of $\mathbf{V}_G$ and $\mathbf{V}_E$ suffice. Importantly, these definitions ensure $\mathbf{V}_G$ and $\mathbf{V}_E$ are always valid covariance matrices (i.e., they are both at least positive semidefinite).

**Partial derivatives.** Let $\gamma$ denote the genetic parameter in row $t$ and column $f$ of $\mathbf{C}_G$. Thus, it is the weight of the path from genetic factor $f$ to trait $t$ or, put differently, it is the effect of some latent genetic factor $f$ (with unit variance) on trait $t$. By the product rule, the partial derivative of $\mathbf{V}_G$ with respect to this parameter is:

$$\frac{\partial \mathbf{V}_G}{\partial \gamma} = \frac{\partial \mathbf{C}_G}{\partial \gamma}\mathbf{C}_G^\top + \mathbf{C}_G \frac{\partial \mathbf{C}_G^\top}{\partial \gamma} \tag{85}$$

$$= \frac{\partial \mathbf{C}_G}{\partial \gamma}\mathbf{C}_G^\top + \left(\frac{\partial \mathbf{C}_G}{\partial \gamma}\mathbf{C}_G^\top\right)^\top. \tag{86}$$

Notice that the partial derivative of $\mathbf{C}_G$ with respect to $\gamma$ is zero everywhere, except for the element in row $t$ and column $f$. By recognising that we can write $\mathbf{C}_G$ as $(\mathbf{c}_{G1} \quad \ldots \quad \mathbf{c}_{GF_G})$, where $\mathbf{c}_{Gf}$ is $T \times 1$ vector of coefficients from factor $f$ to all traits, it follows that

$$\frac{\partial \mathbf{C}_G}{\partial \gamma}\mathbf{C}_G^\top = \frac{\partial \mathbf{C}_G}{\partial \gamma}\begin{pmatrix} \mathbf{c}_{G1}^\top \\ \vdots \\ \mathbf{c}_{GF_G}^\top \end{pmatrix} \tag{87}$$

$$= \begin{pmatrix} \mathbf{0}_{(t-1)\times T} \\ \mathbf{c}_{Gf}^\top \\ \mathbf{0}_{(T-t)\times T} \end{pmatrix}. \tag{88}$$

Therefore,

$$\frac{\partial \mathbf{V}_G}{\partial \gamma} = \begin{pmatrix} \mathbf{0}_{T\times(t-1)} & \mathbf{c}_{Gf} & \mathbf{0}_{T\times(T-t)} \end{pmatrix} + \begin{pmatrix} \mathbf{0}_{(t-1)\times T} \\ \mathbf{c}_{Gf}^\top \\ \mathbf{0}_{(T-t)\times T} \end{pmatrix}. \tag{89}$$

Analogously, with $\varepsilon$ denoting the coefficient from environmental factor $f$ to trait $t$, the partial derivative of $\mathbf{V}_E$ with respect to $\varepsilon$ is:

$$\frac{\partial \mathbf{V}_E}{\partial \varepsilon} = \begin{pmatrix} \mathbf{0}_{T\times(t-1)} & \mathbf{c}_{Ef} & \mathbf{0}_{T\times(T-t)} \end{pmatrix} + \begin{pmatrix} \mathbf{0}_{(t-1)\times T} \\ \mathbf{c}_{Ef}^\top \\ \mathbf{0}_{(T-t)\times T} \end{pmatrix}. \tag{90}$$

Based on these expressions, it follows that:

$$\frac{\partial \mathbf{V}_j}{\partial \gamma} = d_j \left( \begin{pmatrix} \mathbf{0}_{T\times(t-1)} & \mathbf{c}_{Gf} & \mathbf{0}_{T\times(T-t)} \end{pmatrix} + \begin{pmatrix} \mathbf{0}_{(t-1)\times T} \\ \mathbf{c}_{Gf}^\top \\ \mathbf{0}_{(T-t)\times T} \end{pmatrix} \right) \text{ and } \tag{91}$$

$$\frac{\partial \mathbf{V}_j}{\partial \varepsilon} = \left( \begin{pmatrix} \mathbf{0}_{T\times(t-1)} & \mathbf{c}_{Ef} & \mathbf{0}_{T\times(T-t)} \end{pmatrix} + \begin{pmatrix} \mathbf{0}_{(t-1)\times T} \\ \mathbf{c}_{Ef}^\top \\ \mathbf{0}_{(T-t)\times T} \end{pmatrix} \right). \tag{92}$$

Hence, the partial derivatives of the covariance matrices for observations $j = 1, \ldots, N$ are independent of observation-specific terms for partial derivatives with respect to environmental parameters. Moreover, for genetic parameters, except for a scaling coefficient $d_j$ (i.e., the EVs from the GRM), the partial derivatives of the covariance matrices for observations $j = 1, \ldots, N$ are also independent of observation-specific terms. This insight is useful in reducing the computational complexity of the gradient and AI matrix.

#### Step 7. Calculating the gradient efficiently

From the expression of the gradient of the log-likelihood in Eq. 17, it follows that we need efficient expressions for the squared sum of residuals and two traces.

**Squared sum of residuals.** Here, we seek an efficient expression for $\sum_{j=1}^{N} \mathbf{r}_j^\top \frac{\partial \mathbf{V}_j}{\partial \theta_1} \mathbf{r}_j$. In Step 5, we already derived efficient expressions for computing $\mathbf{r}_j$ for $j = 1, \ldots, N$ in $O\left(NT^2\right)$ time. For $\theta_1$ being a genetic parameter $\gamma$, denoting the coefficient from genetic factor $f$ to trait $t$, we therefore obtain the following expression:

$$\sum_{j=1}^{N} \mathbf{r}_j^\top \frac{\partial \mathbf{V}_j}{\partial \theta_1} \mathbf{r}_j = \sum_{j=1}^{N} d_j \mathbf{r}_j^\top \left[ \begin{pmatrix} \mathbf{0}_{T\times(t-1)} & \mathbf{c}_{Gf} & \mathbf{0}_{T\times(T-t)} \end{pmatrix} + \begin{pmatrix} \mathbf{0}_{(t-1)\times T} \\ \mathbf{c}_{Gf}^\top \\ \mathbf{0}_{(T-t)\times T} \end{pmatrix} \right] \mathbf{r}_j \tag{93}$$

$$= \sum_{j=1}^{N} d_j \left[ \begin{pmatrix} \mathbf{0}_{1\times(t-1)} & \mathbf{r}_j^\top \mathbf{c}_{Gf} & \mathbf{0}_{1\times(T-t)} \end{pmatrix} \mathbf{r}_j + \mathbf{r}_j^\top \begin{pmatrix} \mathbf{0}_{(t-1)\times 1} \\ \mathbf{c}_{Gf}^\top \mathbf{r}_j \\ \mathbf{0}_{(T-t)\times 1} \end{pmatrix} \right] \tag{94}$$

$$= \sum_{j=1}^{N} d_j \left[ \mathbf{r}_j^\top \mathbf{c}_{Gf} R_{tj} + R_{jt} \mathbf{c}_{Gf}^\top \mathbf{r}_j \right] \tag{95}$$

$$= 2 \sum_{j=1}^{N} R_{tj} d_j \left( \mathbf{r}_j^\top \mathbf{c}_{Gf} \right). \tag{96}$$

Here, $R_{tj}$ denotes element $t, j$ from the $T \times N$ matrix of rescaled GLS residuals, $\mathbf{R}$. Thus, $R_{tj}$ is the rescaled GLS residual for individual $j$ when explaining trait $t$. We can rewrite this expression much more compactly and efficiently as:

$$\sum_{j=1}^{N} \mathbf{r}_j^\top \frac{\partial \mathbf{V}_j}{\partial \theta_1} \mathbf{r}_j = \left\{ 2\mathbf{R}\mathbf{D}\mathbf{R}^\top \mathbf{C}_G \right\}_{tf}, \tag{97}$$

where $\{\mathbf{A}\}_{ij}$ denotes the element in row $i$ and column $j$ of a given matrix $\mathbf{A}$, and where $\mathbf{D}$ denotes the diagonal matrix with eigenvalues from the GRM. As $\mathbf{D}$ is a diagonal matrix, calculating $\mathbf{R}\mathbf{D}\mathbf{R}^\top \mathbf{C}_G$ is trivial (e.g., construct $\widetilde{\mathbf{R}} = \mathbf{R}\mathbf{D}^{\frac{1}{2}}$ and take $\widetilde{\mathbf{R}}^\top \widetilde{\mathbf{R}} \mathbf{C}_G$). Importantly, notice that the matrix product $2\mathbf{R}\mathbf{D}\mathbf{R}^\top \mathbf{C}_G$ provides the values of $\sum_{j=1}^{N} \mathbf{r}_j^\top \frac{\partial \mathbf{V}_j}{\partial \theta_1} \mathbf{r}_j$ for all genetic parameters. Therefore, given some genetic parameter $\gamma$, we just need to take the appropriate row and column from this matrix to obtain the contribution of $\sum_{j=1}^{N} \mathbf{r}_j^\top \frac{\partial \mathbf{V}_j}{\partial \theta_1} \mathbf{r}_j$ to the element of the gradient vector that corresponds to that parameter. Analogously, for $\theta_1$ being a environmental parameter $\varepsilon$ denoting the coefficient from environmental factor $f$ to trait $t$, we get:

$$\sum_{j=1}^{N} \mathbf{r}_j^\top \frac{\partial \mathbf{V}_j}{\partial \varepsilon} \mathbf{r}_j = \left\{ 2\mathbf{R}\mathbf{R}^\top \mathbf{C}_E \right\}_{tf}. \tag{98}$$

Notice that by computing $2\mathbf{R}\mathbf{D}\mathbf{R}^\top \mathbf{C}_G$ and $2\mathbf{R}\mathbf{R}^\top \mathbf{C}_E$, we have two matrices of which the appropriate entries are the values of $\sum_{j=1}^{N} \mathbf{r}_j^\top \frac{\partial \mathbf{V}_j}{\partial \theta_1} \mathbf{r}_j$ for all parameters in our model. Thus, calculating this part of the gradient has a computational complexity equal to that of computing these two matrices. This can be done in $O\left(NT^2\right)$

time, because the number of factors in our model is at most proportional to the number of traits.

**First trace.** Next, we need an efficient expression for $\sum_{j=1}^{N} \mathrm{tr}\left(\mathbf{V}_j^{-1}\frac{\partial \mathbf{V}_j}{\partial \theta_1}\right)$. First, let us define

$$\mathbf{T}_G = \sum_{j=1}^{N} \mathbf{V}_j^{-1} d_j \text{ and } \mathbf{T}_E = \sum_{j=1}^{N} \mathbf{V}_j^{-1}. \tag{99}$$

By substituting $\mathbf{V}_j^{-1}$ with the expression for it in Eq. 34, we get:

$$\mathbf{T}_G = \mathbf{F}\left(\sum_{j=1}^{N} d_j \mathbf{D}_j^*\right)\mathbf{F}^\top \text{ and } \mathbf{T}_E = \mathbf{F}\left(\sum_{j=1}^{N} \mathbf{D}_j^*\right)\mathbf{F}^\top. \tag{100}$$

These expressions imply that both $\mathbf{T}_G$ and $\mathbf{T}_E$ can be obtained by taking a sum of diagonal matrices, and by respectively pre- and post-multiplying the resultant matrices by $\mathbf{F}$ and $\mathbf{F}^\top$.

Now, let us consider a genetic parameter, $\gamma$, denoting the effect of genetic factor $f$ on trait $t$. In this case, we have that:

$$\begin{aligned}
\sum_{j=1}^{N} \mathrm{tr}\left(\mathbf{V}_j^{-1}\frac{\partial \mathbf{V}_j}{\partial \gamma}\right) &= \mathrm{tr}\left(\sum_{j=1}^{N}\left(\mathbf{V}_j^{-1} d_j \frac{\partial \mathbf{V}_G}{\partial \gamma}\right)\right) = \mathrm{tr}\left(\left(\sum_{j=1}^{N}\mathbf{V}_j^{-1} d_j\right)\frac{\partial \mathbf{V}_G}{\partial \gamma}\right) = \mathrm{tr}\left(\mathbf{T}_G \frac{\partial \mathbf{V}_G}{\partial \gamma}\right) && (101)\\
&= \mathrm{tr}\left(\mathbf{T}_G\left(\begin{pmatrix} \mathbf{0}_{T\times(t-1)} & \mathbf{c}_{Gf} & \mathbf{0}_{T\times(T-t)} \end{pmatrix} + \begin{pmatrix} \mathbf{0}_{(t-1)\times T} \\ \mathbf{c}_{Gf}^\top \\ \mathbf{0}_{(T-t)\times T} \end{pmatrix}\right)\right) && (102)\\
&= \mathrm{tr}\left(\begin{pmatrix} \mathbf{0}_{T\times(t-1)} & \mathbf{T}_G\mathbf{c}_{Gf} & \mathbf{0}_{T\times(T-t)} \end{pmatrix}\right) + \mathrm{tr}\left(\mathbf{T}_G \begin{pmatrix} \mathbf{0}_{(t-1)\times T} \\ \mathbf{c}_{Gf}^\top \\ \mathbf{0}_{(T-t)\times T} \end{pmatrix}\right) && (103)\\
&= 2\mathrm{tr}\left(\begin{pmatrix} \mathbf{0}_{T\times(t-1)} & \mathbf{T}_G\mathbf{c}_{Gf} & \mathbf{0}_{T\times(T-t)} \end{pmatrix}\right) && (104)\\
&= \{2\mathbf{T}_G\mathbf{C}_G\}_{tf}\,. && (105)
\end{aligned}$$

Analogously, for environmental parameter $\varepsilon$, indicating the effect of environmental factor $f$ on trait $t$, we have:

$$\sum_{j=1}^{N} \mathrm{tr}\left(\mathbf{V}_j^{-1}\frac{\partial \mathbf{V}_j}{\partial \varepsilon}\right) = \{2\mathbf{T}_E\mathbf{C}_E\}_{tf}\,. \tag{106}$$

Matrices $\mathbf{T}_E$ and $\mathbf{T}_G$ can be obtained in $O(NT)$ time. Further calculation of $2\mathbf{T}_G\mathbf{C}_G$ and $2\mathbf{T}\mathbf{C}_E$ is trivial. By calculating these two matrices and taking the appropriate elements, we have the contribution of each parameter in our model to the gradient with respect to the $\sum_{j=1}^{N} \operatorname{tr}\left(\mathbf{V}_j^{-1}\frac{\partial \mathbf{V}_j}{\partial \theta_1}\right)$ term. Thus, calculating this term for all parameters in the model can be done in $O(NT)$ time.

**Second trace.** To derive an efficient expression for the second trace, we start with the case of different covariates for each traits and then consider the special case of equal covariates across traits. To see how the computational complexity of the second trace can be reduced, even when we have different covariates per trait, first note that our aim is to find an efficient expression for the following term from Eq. 17:

$$\operatorname{tr}\left(\mathbf{Z}^{-1}\left(\sum_{j=1}^{N}\mathbf{X}_j^\top\mathbf{V}_j^{-1}\frac{\partial \mathbf{V}_j}{\partial \theta_1}\mathbf{V}_j^{-1}\mathbf{X}_j\right)\right). \tag{107}$$

Here, we cannot use properties of the Kronecker product to make certain terms cancel out with respect to the inverse of $\mathbf{Z}$. Therefore, our derivations focus here on finding an efficient expression for $\mathbf{X}_j^\top\mathbf{V}_j^{-1}\frac{\partial \mathbf{V}_j}{\partial \theta_1}\mathbf{V}_j^{-1}\mathbf{X}_j$. However, it is important to recognise that it is still possible to write our matrix of covariates as a Kronecker product, post-multiplied by $\mathbf{S}^\top$. That is, we have that:

$$\mathbf{X}_j = \left(\mathbf{I}_T \otimes \mathbf{x}_j^\top\right)\mathbf{S}^\top. \tag{108}$$

Let's first focus on the term for $\gamma$. Because of the constancy of the partial derivatives for $\mathbf{V}_j$ with respect to $\gamma$ over the observations (except for scalar $d_j$), we have that:

$$\begin{aligned}
\sum_{j=1}^{N}\mathbf{X}_j^\top\mathbf{V}_j^{-1}\frac{\partial \mathbf{V}_j}{\partial \gamma}\mathbf{V}_j^{-1}\mathbf{X}_j &= \sum_{j=1}^{N} d_j\mathbf{X}_j^\top\mathbf{V}_j^{-1}\frac{\partial \mathbf{V}_G}{\partial \gamma}\mathbf{V}_j^{-1}\mathbf{X}_j && (109)\\
&= \sum_{j=1}^{N} d_j\mathbf{S}\left(\mathbf{I}_T\otimes\mathbf{x}_j\right)\mathbf{F}\mathbf{D}_j^*\mathbf{F}^\top\frac{\partial \mathbf{V}_G}{\partial \gamma}\mathbf{F}\mathbf{D}_j^*\mathbf{F}^\top\left(\mathbf{I}_T\otimes\mathbf{x}_j^\top\right)\mathbf{S}^\top && (110)\\
&= \mathbf{S}\left[\sum_{j=1}^{N} d_j\left(\mathbf{I}_T\otimes\mathbf{x}_j\right)\mathbf{F}\mathbf{D}_j^*\mathbf{F}^\top\frac{\partial \mathbf{V}_G}{\partial \gamma}\mathbf{F}\mathbf{D}_j^*\mathbf{F}^\top\left(\mathbf{I}_T\otimes\mathbf{x}_j^\top\right)\right]\mathbf{S}^\top. && (111)
\end{aligned}$$

Thus, using $\mathbf{S}$ to select the appropriate submatrix of the $Tk\times Tk$ matrix between square brackets is something that can be done after aggregating observations. Therefore, we can focus on finding an efficient expression for the term in between the square brackets. For that term, we have that:

$$\sum_{j=1}^{N} d_j \left(\mathbf{I}_T \otimes \mathbf{x}_j\right) \mathbf{F}\mathbf{D}_j^* \mathbf{F}^\top \frac{\partial \mathbf{V}_G}{\partial \gamma} \mathbf{F}\mathbf{D}_j^* \mathbf{F}^\top \left(\mathbf{I}_T \otimes \mathbf{x}_j^\top\right) \tag{112}$$

$$= \sum_{j=1}^{N} d_j \left(\mathbf{I}_T \otimes \mathbf{x}_j\right) \left(\mathbf{F} \otimes \mathbf{I}_1\right) \mathbf{D}_j^* \mathbf{F}^\top \frac{\partial \mathbf{V}_G}{\partial \gamma} \mathbf{F}\mathbf{D}_j^* \left(\mathbf{F}^\top \otimes \mathbf{I}_1\right) \left(\mathbf{I}_T \otimes \mathbf{x}_j^\top\right) \tag{113}$$

$$= \sum_{j=1}^{N} d_j \left(\mathbf{F} \otimes \mathbf{x}_j\right) \mathbf{D}_j^* \mathbf{F}^\top \frac{\partial \mathbf{V}_G}{\partial \gamma} \mathbf{F}\mathbf{D}_j^* \left(\mathbf{F}^\top \otimes \mathbf{x}_j^\top\right) \tag{114}$$

$$= \sum_{j=1}^{N} d_j \left(\mathbf{F} \otimes \mathbf{I}_k\right) \left(\mathbf{I}_T \otimes \mathbf{x}_j\right) \mathbf{D}_j^* \mathbf{F}^\top \frac{\partial \mathbf{V}_G}{\partial \gamma} \mathbf{F}\mathbf{D}_j^* \left(\mathbf{I}_T \otimes \mathbf{x}_j^\top\right) \left(\mathbf{F}^\top \otimes \mathbf{I}_k\right) \tag{115}$$

$$= \left(\mathbf{F} \otimes \mathbf{I}_k\right) \left[\sum_{j=1}^{N} d_j \left(\mathbf{I}_T \otimes \mathbf{x}_j\right) \mathbf{D}_j^* \mathbf{F}^\top \frac{\partial \mathbf{V}_G}{\partial \gamma} \mathbf{F}\mathbf{D}_j^* \left(\mathbf{I}_T \otimes \mathbf{x}_j^\top\right)\right] \left(\mathbf{F}^\top \otimes \mathbf{I}_k\right) \tag{116}$$

$$= \left(\mathbf{F} \otimes \mathbf{I}_k\right) \left[\sum_{j=1}^{N} \left(\mathbf{I}_T \otimes \mathbf{x}_j\right) \left(\left(d_j \mathbf{D}_j^* \mathbf{F}^\top \frac{\partial \mathbf{V}_G}{\partial \gamma} \mathbf{F}\mathbf{D}_j^*\right) \otimes \mathbf{I}_1\right) \left(\mathbf{I}_T \otimes \mathbf{x}_j^\top\right)\right] \left(\mathbf{F}^\top \otimes \mathbf{I}_k\right) \tag{117}$$

$$= \left(\mathbf{F} \otimes \mathbf{I}_k\right) \left[\sum_{j=1}^{N} \left(\left(d_j \mathbf{D}_j^* \mathbf{F}^\top \frac{\partial \mathbf{V}_G}{\partial \gamma} \mathbf{F}\mathbf{D}_j^*\right) \otimes \left(\mathbf{x}_j \mathbf{x}_j^\top\right)\right)\right] \left(\mathbf{F}^\top \otimes \mathbf{I}_k\right). \tag{118}$$

The Kronecker product in the summand can be written as:

$$\left(d_j \mathbf{D}_j^* \mathbf{F}^\top \frac{\partial \mathbf{V}_G}{\partial \gamma} \mathbf{F}\mathbf{D}_j^*\right) \otimes \left(\mathbf{x}_j \mathbf{x}_j^\top\right) \tag{119}$$

$$= \left(\left(\mathbf{D}_j^* \mathbf{F}^\top\right) \otimes \mathbf{I}_k\right) \left(\left(d_j \frac{\partial \mathbf{V}_G}{\partial \gamma}\right) \otimes \left(\mathbf{x}_j \mathbf{x}_j^\top\right)\right) \left(\left(\mathbf{F}\mathbf{D}_j^*\right) \otimes \mathbf{I}_k\right). \tag{120}$$

The middle Kronecker product in the preceding expression can be written as:

$$d_j \left[\begin{pmatrix} \mathbf{0}_{T\times(t-1)} & \mathbf{c}_{Gf} & \mathbf{0}_{T\times(T-t)} \end{pmatrix} \otimes \left(\mathbf{x}_j \mathbf{x}_j^\top\right) + \begin{pmatrix} \mathbf{0}_{(t-1)\times T} \\ \mathbf{c}_{Gf}^\top \\ \mathbf{0}_{(T-t)\times T} \end{pmatrix} \otimes \left(\mathbf{x}_j \mathbf{x}_j^\top\right)\right] \tag{121}$$

$$= d_j \begin{pmatrix} \mathbf{0}_{k\times(t-1)k} & c_{Gf1} \mathbf{x}_j \mathbf{x}_j^\top & \mathbf{0}_{k\times(T-t)k} \\ \vdots & \vdots & \vdots \\ \mathbf{0}_{k\times(t-1)k} & c_{GfT} \mathbf{x}_j \mathbf{x}_j^\top & \mathbf{0}_{k\times(T-t)k} \end{pmatrix} + d_j \begin{pmatrix} \mathbf{0}_{k\times(t-1)k} & c_{Gf1} \mathbf{x}_j \mathbf{x}_j^\top & \mathbf{0}_{k\times(T-t)k} \\ \vdots & \vdots & \vdots \\ \mathbf{0}_{k\times(t-1)k} & c_{GfT} \mathbf{x}_j \mathbf{x}_j^\top & \mathbf{0}_{k\times(T-t)k} \end{pmatrix}^\top, \tag{122}$$

with $c_{Gft}$ denoting the genetic effect of genetic factor $f$ on trait $t$. Thus, the Kronecker product in the summand can be written as:

$$\left(d_j \mathbf{D}_j^* \mathbf{F}^\top \frac{\partial \mathbf{V}_G}{\partial \gamma} \mathbf{F} \mathbf{D}_j^*\right) \otimes \left(\mathbf{x}_j \mathbf{x}_j^\top\right) \tag{123}$$

$$= d_j \mathbf{H}_j + d_j \mathbf{H}_j^\top, \tag{124}$$

where

$$\mathbf{H}_j = \left(\left(\mathbf{D}_j^* \mathbf{F}^\top\right) \otimes \mathbf{I}_k\right) \begin{pmatrix} \mathbf{0}_{k\times(t-1)k} & c_{Gf1}\mathbf{x}_j\mathbf{x}_j^\top & \mathbf{0}_{k\times(T-t)k} \\ \vdots & \vdots & \vdots \\ \mathbf{0}_{k\times(t-1)k} & c_{GfT}\mathbf{x}_j\mathbf{x}_j^\top & \mathbf{0}_{k\times(T-t)k} \end{pmatrix} \left(\left(\mathbf{F}\mathbf{D}_j^*\right) \otimes \mathbf{I}_k\right) \tag{125}$$

$$= \begin{pmatrix} \mathbf{0}_{k\times(t-1)k} & \mathbf{x}_j\mathbf{x}_j^\top \left(d_j\lambda_1+1\right)^{-1} \sum_{s=1}^T c_{Gfs}F_{s1} & \mathbf{0}_{k\times(T-t)k} \\ \vdots & \vdots & \vdots \\ \mathbf{0}_{k\times(t-1)k} & \mathbf{x}_j\mathbf{x}_j^\top \left(d_j\lambda_T+1\right)^{-1} \sum_{s=1}^T c_{Gfs}F_{sT} & \mathbf{0}_{k\times(T-t)k} \end{pmatrix} \left(\left(\mathbf{F}\mathbf{D}_j^*\right) \otimes \mathbf{I}_k\right) \tag{126}$$

$$= \begin{pmatrix} \mathbf{0}_{k\times(t-1)k} & \mathbf{x}_j\mathbf{x}_j^\top \left(d_j\lambda_1+1\right)^{-1} \left\{\mathbf{c}_{Gf}^\top\mathbf{F}\right\}_1 & \mathbf{0}_{k\times(T-t)k} \\ \vdots & \vdots & \vdots \\ \mathbf{0}_{k\times(t-1)k} & \mathbf{x}_j\mathbf{x}_j^\top \left(d_j\lambda_T+1\right)^{-1} \left\{\mathbf{c}_{Gf}^\top\mathbf{F}\right\}_T & \mathbf{0}_{k\times(T-t)k} \end{pmatrix} \left(\left(\mathbf{F}\mathbf{D}_j^*\right) \otimes \mathbf{I}_k\right) \tag{127}$$

$$= \begin{pmatrix} \mathbf{x}_j\mathbf{x}_j^\top \left(d_j\lambda_1+1\right)^{-1}\left(d_j\lambda_1+1\right)^{-1} \left\{\mathbf{c}_{Gf}^\top\mathbf{F}\right\}_1 F_{t1} & \dots & \mathbf{x}_j\mathbf{x}_j^\top \left(d_j\lambda_1+1\right)^{-1}\left(d_j\lambda_T+1\right)^{-1} \left\{\mathbf{c}_{Gf}^\top\mathbf{F}\right\}_1 F_{tT} \\ \vdots & & \vdots \\ \mathbf{x}_j\mathbf{x}_j^\top \left(d_j\lambda_T+1\right)^{-1}\left(d_j\lambda_1+1\right)^{-1} \left\{\mathbf{c}_{Gf}^\top\mathbf{F}\right\}_T F_{t1} & \dots & \mathbf{x}_j\mathbf{x}_j^\top \left(d_j\lambda_T+1\right)^{-1}\left(d_j\lambda_T+1\right)^{-1} \left\{\mathbf{c}_{Gf}^\top\mathbf{F}\right\}_T F_{tT} \end{pmatrix} \tag{128}$$

$$= \begin{pmatrix} \mathbf{x}_j\mathbf{x}_j^\top \left(d_j\lambda_1+1\right)^{-1}\left(d_j\lambda_1+1\right)^{-1} \left\{\mathbf{C}_{G}^\top\mathbf{F}\right\}_{f1} F_{t1} & \dots & \mathbf{x}_j\mathbf{x}_j^\top \left(d_j\lambda_1+1\right)^{-1}\left(d_j\lambda_T+1\right)^{-1} \left\{\mathbf{C}_{G}^\top\mathbf{F}\right\}_{f1} F_{tT} \\ \vdots & & \vdots \\ \mathbf{x}_j\mathbf{x}_j^\top \left(d_j\lambda_T+1\right)^{-1}\left(d_j\lambda_1+1\right)^{-1} \left\{\mathbf{C}_{G}^\top\mathbf{F}\right\}_{fT} F_{t1} & \dots & \mathbf{x}_j\mathbf{x}_j^\top \left(d_j\lambda_T+1\right)^{-1}\left(d_j\lambda_T+1\right)^{-1} \left\{\mathbf{C}_{G}^\top\mathbf{F}\right\}_{fT} F_{tT} \end{pmatrix}. \tag{129}$$

In this expression, $\{\mathbf{v}\}_i$ denotes element $i$ in a given vector $\mathbf{v}$ and $F_{ij}$ denotes element $i, j$ in $\mathbf{F}$. As a result,

$$\sum_{j=1}^{N} d_j \left(\mathbf{I}_T \otimes \mathbf{x}_j\right) \mathbf{F}\mathbf{D}_j^* \mathbf{F}^\top \frac{\partial \mathbf{V}_G}{\partial \gamma} \mathbf{F}\mathbf{D}_j^* \mathbf{F}^\top \left(\mathbf{I}_T \otimes \mathbf{x}_j^\top\right) \tag{130}$$

$$= \left(\mathbf{F} \otimes \mathbf{I}_k\right) \left[\sum_{j=1}^{N} d_j \mathbf{H}_j + d_j \mathbf{H}_j^\top\right] \left(\mathbf{F}^\top \otimes \mathbf{I}_k\right) \tag{131}$$

$$= \left[\left(\mathbf{F} \otimes \mathbf{I}_k\right) \left(\sum_{j=1}^{N} d_j \mathbf{H}_j\right) \left(\mathbf{F}^\top \otimes \mathbf{I}_k\right)\right] + \left[\left(\mathbf{F} \otimes \mathbf{I}_k\right) \left(\sum_{j=1}^{N} d_j \mathbf{H}_j\right)^\top \left(\mathbf{F}^\top \otimes \mathbf{I}_k\right)\right]. \tag{132}$$

Therefore,

$$\mathrm{tr}\left(\mathbf{Z}^{-1} \left(\sum_{j=1}^{N} \mathbf{X}_j^\top \mathbf{V}_j^{-1} \frac{\partial \mathbf{V}_j}{\partial \gamma} \mathbf{V}_j^{-1} \mathbf{X}_j\right)\right) \tag{133}$$

$$= \mathrm{tr}\left(\mathbf{Z}^{-1}\mathbf{S} \left(\left[\left(\mathbf{F} \otimes \mathbf{I}_k\right) \left(\sum_{j=1}^{N} d_j \mathbf{H}_j\right) \left(\mathbf{F}^\top \otimes \mathbf{I}_k\right)\right] + \left[\left(\mathbf{F} \otimes \mathbf{I}_k\right) \left(\sum_{j=1}^{N} d_j \mathbf{H}_j\right)^\top \left(\mathbf{F}^\top \otimes \mathbf{I}_k\right)\right]\right) \mathbf{S}^\top\right) \tag{134}$$

$$= \mathrm{tr}\left(\mathbf{Z}^{-1}\mathbf{S} \left[\left(\mathbf{F} \otimes \mathbf{I}_k\right) \left(\sum_{j=1}^{N} d_j \mathbf{H}_j\right) \left(\mathbf{F}^\top \otimes \mathbf{I}_k\right)\right] \mathbf{S}^\top\right) + \mathrm{tr}\left(\mathbf{Z}^{-1}\mathbf{S} \left[\left(\mathbf{F} \otimes \mathbf{I}_k\right) \left(\sum_{j=1}^{N} d_j \mathbf{H}_j\right)^\top \left(\mathbf{F}^\top \otimes \mathbf{I}_k\right)\right] \mathbf{S}^\top\right) \tag{135}$$

$$= \mathrm{tr}\left(\mathbf{Z}^{-1}\mathbf{S} \left[\left(\mathbf{F} \otimes \mathbf{I}_k\right) \left(\sum_{j=1}^{N} d_j \mathbf{H}_j\right) \left(\mathbf{F}^\top \otimes \mathbf{I}_k\right)\right] \mathbf{S}^\top\right) + \mathrm{tr}\left(\mathbf{S} \left[\left(\mathbf{F} \otimes \mathbf{I}_k\right) \left(\sum_{j=1}^{N} d_j \mathbf{H}_j\right) \left(\mathbf{F}^\top \otimes \mathbf{I}_k\right)\right] \mathbf{S}^\top \mathbf{Z}^{-1}\right) \tag{136}$$

$$= \mathrm{tr}\left(\mathbf{Z}^{-1}\mathbf{S} \left[\left(\mathbf{F} \otimes \mathbf{I}_k\right) \left(\sum_{j=1}^{N} d_j \mathbf{H}_j\right) \left(\mathbf{F}^\top \otimes \mathbf{I}_k\right)\right] \mathbf{S}^\top\right) + \mathrm{tr}\left(\mathbf{Z}^{-1}\mathbf{S} \left[\left(\mathbf{F} \otimes \mathbf{I}_k\right) \left(\sum_{j=1}^{N} d_j \mathbf{H}_j\right) \left(\mathbf{F}^\top \otimes \mathbf{I}_k\right)\right] \mathbf{S}^\top\right) \tag{137}$$

$$= 2\mathrm{tr}\left(\mathbf{Z}^{-1}\mathbf{S} \left[\left(\mathbf{F} \otimes \mathbf{I}_k\right) \left(\sum_{j=1}^{N} d_j \mathbf{H}_j\right) \left(\mathbf{F}^\top \otimes \mathbf{I}_k\right)\right] \mathbf{S}^\top\right). \tag{138}$$

Thus, our efforts to find an efficient expression for the second trace lead us to finding an efficient expression for $\sum_{j=1}^{N} d_j \mathbf{H}_j$. Note here that:

$$\sum_{j=1}^{N} d_j \mathbf{H}_j = \left(\text{diag}\left(\left\{\mathbf{C}_G^\top \mathbf{F}\right\}_{f\bullet}\right) \otimes \mathbf{I}_k\right) \mathbf{B} \left(\text{diag}\left(\{\mathbf{F}\}_{t\bullet}\right) \otimes \mathbf{I}_k\right), \tag{139}$$

with

$$\text{diag}\left(\left\{\mathbf{C}_G^\top \mathbf{F}\right\}_{f\bullet}\right) = \begin{pmatrix} \left\{\mathbf{C}_G^\top \mathbf{F}\right\}_{f1} & & 0 \\ & \ddots & \\ 0 & & \left\{\mathbf{C}_G^\top \mathbf{F}\right\}_{fT} \end{pmatrix}, \tag{140}$$

$$\text{diag}\left(\{\mathbf{F}\}_{t\bullet}\right) = \begin{pmatrix} F_{t1} & & 0 \\ & \ddots & \\ 0 & & F_{tT} \end{pmatrix}, \tag{141}$$

$$\mathbf{B}_G = \begin{pmatrix} \mathbf{B}_{G11} & \dots & \mathbf{B}_{G1T} \\ \vdots & & \vdots \\ \mathbf{B}_{G1T} & \dots & \mathbf{B}_{GTT} \end{pmatrix}, \tag{142}$$

$$\mathbf{B}_{Gst} = \mathbf{X}^\top \mathbf{D}\mathbf{D}_s^\dagger \mathbf{D}_t^\dagger \mathbf{X}. \tag{143}$$

Therefore,

$$\text{tr}\left(\mathbf{Z}^{-1}\left(\sum_{j=1}^{N} \mathbf{X}_j^\top \mathbf{V}_j^{-1} \frac{\partial \mathbf{V}_j}{\partial \gamma} \mathbf{V}_j^{-1} \mathbf{X}_j\right)\right) \tag{144}$$

$$= 2\text{tr}\left(\mathbf{Z}^{-1}\mathbf{S}\left[(\mathbf{F} \otimes \mathbf{I}_k)\left(\left(\text{diag}\left(\left\{\mathbf{C}_G^\top \mathbf{F}\right\}_{f\bullet}\right) \otimes \mathbf{I}_k\right) \mathbf{B}_G \left(\text{diag}\left(\{\mathbf{F}\}_{t\bullet}\right) \otimes \mathbf{I}_k\right)\right)\left(\mathbf{F}^\top \otimes \mathbf{I}_k\right)\right]\mathbf{S}^\top\right). \tag{145}$$

With our previously defined binary $K \times Tk$ selector matrix $\mathbf{S}$, we can now rewrite our trace as follows:

$$\text{tr}\left(\mathbf{Z}^{-1}\left(\sum_{j=1}^{N} \mathbf{X}_j^\top \mathbf{V}_j^{-1} \frac{\partial \mathbf{V}_j}{\partial \gamma} \mathbf{V}_j^{-1} \mathbf{X}_j\right)\right) \tag{146}$$

$$= 2\text{tr}\left(\left(\text{diag}\left(\left\{\mathbf{C}_G^\top \mathbf{F}\right\}_{f\bullet}\right) \otimes \mathbf{I}_k\right) \mathbf{B}_G \left(\text{diag}\left(\{\mathbf{F}\}_{t\bullet}\right) \otimes \mathbf{I}_k\right)\left[\left(\mathbf{F}^\top \otimes \mathbf{I}_k\right)\mathbf{S}^\top \mathbf{Z}^{-1} \mathbf{S}\left(\mathbf{F} \otimes \mathbf{I}_k\right)\right]\right). \tag{147}$$

In the last expression, notice that $\operatorname{diag}\left(\left\{\mathbf{C}_G^\top \mathbf{F}\right\}_{f\bullet}\right) \otimes \mathbf{I}_k$ and $\operatorname{diag}\left(\{\mathbf{F}\}_{t\bullet}\right) \otimes \mathbf{I}_k$ are both diagonal matrices. Therefore, by properties of the Hadamard product, we have that:

$$\operatorname{tr}\left(\mathbf{Z}^{-1}\left(\sum_{j=1}^{N} \mathbf{X}_j^\top \mathbf{V}_j^{-1} \frac{\partial \mathbf{V}_j}{\partial \gamma} \mathbf{V}_j^{-1} \mathbf{X}_j\right)\right) = 2\left(\left\{\mathbf{C}_G^\top \mathbf{F}\right\}_{f\bullet}^{\top} \otimes \boldsymbol{\iota}_{1\times k}\right)\left[\mathbf{B}_G \circ \mathbf{J}\right]\left(\{\mathbf{F}\}_{t\bullet} \otimes \boldsymbol{\iota}_{k\times 1}\right) \tag{148}$$

$$= \left\{2\left(\mathbf{C}_G^\top \mathbf{F} \otimes \boldsymbol{\iota}_{1\times k}\right)\left[\mathbf{B}_G \circ \mathbf{J}\right]\left(\mathbf{F}^\top \otimes \boldsymbol{\iota}_{k\times 1}\right)\right\}_{ft}, \text{ where} \tag{149}$$

$$\mathbf{J} = \left(\mathbf{F}^\top \otimes \mathbf{I}_k\right) \mathbf{S}^\top \mathbf{Z}^{-1} \mathbf{S} \left(\mathbf{F} \otimes \mathbf{I}_k\right), \tag{150}$$

$$\mathbf{B}_G = \begin{pmatrix} \mathbf{B}_{G11} & \dots & \mathbf{B}_{G1T} \\ \vdots & & \vdots \\ \mathbf{B}_{G1T} & \dots & \mathbf{B}_{GTT} \end{pmatrix}, \text{ and} \tag{151}$$

$$\mathbf{B}_{Gst} = \mathbf{X}^\top \mathbf{D} \mathbf{D}_s^\dagger \mathbf{D}_t^\dagger \mathbf{X}. \tag{152}$$

Therefore, to determine the second trace, we need to compute our grand $Tk \times Tk$ matrix $\mathbf{B}_G$. This calculation can be done in $O\left(NT^2\right)$ time. Next, we need to compute the element-wise product of this grand matrix with $\mathbf{J}$, which is readily available from previous steps and computationally trivial. Finally, we need to pre-multiply the resultant matrix by the $F_G \times Tk$ matrix $\mathbf{C}_G^\top \mathbf{F} \otimes \boldsymbol{\iota}_{1\times k}$ and post-multiply by the $Tk \times T$ matrix $\mathbf{F}^\top \otimes \boldsymbol{\iota}_{k\times 1}$, where $F_G$ denotes the number of genetic factors in our model. These two multiplications can be carried out in $O\left(T^3\right)$ time. Thus, overall, the second trace can be computed in $O\left(NT^2\right)$ time, provided $k = O(1)$ and thereby $K = O(T)$.

Analogously, for $\varepsilon$, the effect of environmental factor $f$ on trait $t$, we have that:

$$\operatorname{tr}\left(\mathbf{Z}^{-1}\left(\sum_{j=1}^{N} \mathbf{X}_j^\top \mathbf{V}_j^{-1} \frac{\partial \mathbf{V}_j}{\partial \varepsilon} \mathbf{V}_j^{-1} \mathbf{X}_j\right)\right) = \left\{2\left(\mathbf{C}_E^\top \mathbf{F} \otimes \boldsymbol{\iota}_{1\times k}\right)\left[\mathbf{B}_E \circ \mathbf{J}\right]\left(\mathbf{F}^\top \otimes \boldsymbol{\iota}_{k\times 1}\right)\right\}_{ft}, \text{ where} \tag{153}$$

$$\mathbf{B}_E = \begin{pmatrix} \mathbf{B}_{E11} & \dots & \mathbf{B}_{E1T} \\ \vdots & & \vdots \\ \mathbf{B}_{E1T} & \dots & \mathbf{B}_{ETT} \end{pmatrix} \text{ and} \tag{154}$$

$$\mathbf{B}_{Est} = \mathbf{X}^\top \mathbf{D}_s^\dagger \mathbf{D}_t^\dagger \mathbf{X}. \tag{155}$$

Notice that in case of equal covariates across traits, $\mathbf{J}$ as defined in Eq 150 reduces to $\mathbf{J}$ as defined in Eq 54.

When $\mathbf{S} = \mathbf{I}_{Tk}$, we can substitute $\mathbf{Z}^{-1}$ in Eq. 149 by the efficient expressions in Eq. 53. Doing so yields:

$$\operatorname{tr}\left(\mathbf{Z}^{-1}\left(\sum_{j=1}^{N}\mathbf{X}_j^\top\mathbf{V}_j^{-1}\frac{\partial\mathbf{V}_j}{\partial\gamma}\mathbf{V}_j^{-1}\mathbf{X}_j\right)\right) \tag{156}$$

$$= \left\{2\left(\mathbf{C}_G^\top\mathbf{F}\otimes\boldsymbol{\iota}_{1\times k}\right)\left[\begin{pmatrix}\mathbf{B}_{G11} & \dots & \mathbf{B}_{G1T}\\ \vdots & & \vdots\\ \mathbf{B}_{G1T} & \dots & \mathbf{B}_{GTT}\end{pmatrix}\circ\right.\right. \tag{157}$$

$$\left.\left.\left(\left(\mathbf{F}^\top\otimes\mathbf{I}_k\right)\left(\left(\mathbf{F}^{-1}\right)^\top\otimes\mathbf{I}_k\right)\begin{bmatrix}\mathbf{B}_1^{-1} & & \mathbf{0}\\ & \ddots & \\ \mathbf{0} & & \mathbf{B}_T^{-1}\end{bmatrix}\left(\mathbf{F}^{-1}\otimes\mathbf{I}_k\right)\left(\mathbf{F}\otimes\mathbf{I}_k\right)\right)\right]\left(\mathbf{F}^\top\otimes\boldsymbol{\iota}_{k\times 1}\right)\right\}_{ft} \tag{158}$$

$$= \left\{2\left(\mathbf{C}_G^\top\mathbf{F}\otimes\boldsymbol{\iota}_{1\times k}\right)\left[\begin{pmatrix}\mathbf{B}_{G11} & \dots & \mathbf{B}_{G1T}\\ \vdots & & \vdots\\ \mathbf{B}_{G1T} & \dots & \mathbf{B}_{GTT}\end{pmatrix}\circ\begin{bmatrix}\mathbf{B}_1^{-1} & & \mathbf{0}\\ & \ddots & \\ \mathbf{0} & & \mathbf{B}_T^{-1}\end{bmatrix}\right]\left(\mathbf{F}^\top\otimes\boldsymbol{\iota}_{k\times 1}\right)\right\}_{ft} \tag{159}$$

$$= \left\{2\left(\mathbf{C}_G^\top\mathbf{F}\otimes\boldsymbol{\iota}_{1\times k}\right)\begin{bmatrix}\mathbf{B}_1^{-1}\circ\mathbf{B}_{G11} & & \mathbf{0}\\ & \ddots & \\ \mathbf{0} & & \mathbf{B}_T^{-1}\circ\mathbf{B}_{GTT}\end{bmatrix}\left(\mathbf{F}^\top\otimes\boldsymbol{\iota}_{k\times 1}\right)\right\}_{ft} \tag{160}$$

$$= \left\{2\mathbf{C}_G^\top\mathbf{F}\begin{bmatrix}\boldsymbol{\iota}_{1\times k}\left(\mathbf{B}_1^{-1}\circ\mathbf{B}_{G11}\right)\boldsymbol{\iota}_{k\times 1} & & \mathbf{0}\\ & \ddots & \\ \mathbf{0} & & \boldsymbol{\iota}_{1\times k}\left(\mathbf{B}_T^{-1}\circ\mathbf{B}_{GTT}\right)\boldsymbol{\iota}_{k\times 1}\end{bmatrix}\mathbf{F}^\top\right\}_{ft} \tag{161}$$

$$= \left\{2\mathbf{F}\begin{bmatrix}\boldsymbol{\iota}_{1\times k}\left(\mathbf{B}_1^{-1}\circ\mathbf{B}_{G11}\right)\boldsymbol{\iota}_{k\times 1} & & \mathbf{0}\\ & \ddots & \\ \mathbf{0} & & \boldsymbol{\iota}_{1\times k}\left(\mathbf{B}_T^{-1}\circ\mathbf{B}_{GTT}\right)\boldsymbol{\iota}_{k\times 1}\end{bmatrix}\mathbf{F}^\top\mathbf{C}_G\right\}_{tf}. \tag{162}$$

By virtue of the symmetry of both $\mathbf{B}_t^{-1}$ and $\mathbf{B}_{Gtt}$, we get that:

$$\boldsymbol{\iota}_{1\times k}\left(\mathbf{B}_t^{-1}\circ\mathbf{B}_{Gtt}\right)\boldsymbol{\iota}_{k\times 1} = \operatorname{tr}\left(\mathbf{B}_t^{-1}\mathbf{B}_{Gtt}\right). \tag{163}$$

Thus, in case of identical covariates across traits, we have that:

$$\operatorname{tr}\left(\mathbf{Z}^{-1}\left(\sum_{j=1}^{N}\mathbf{X}_j^\top\mathbf{V}_j^{-1}\frac{\partial\mathbf{V}_j}{\partial\gamma}\mathbf{V}_j^{-1}\mathbf{X}_j\right)\right)$$

$$=\left\{2\mathbf{F}\begin{bmatrix}\operatorname{tr}\left(\mathbf{B}_1^{-1}\mathbf{B}_{G11}\right) & & \mathbf{0}\\ & \ddots & \\ \mathbf{0} & & \operatorname{tr}\left(\mathbf{B}_T^{-1}\mathbf{B}_{GTT}\right)\end{bmatrix}\mathbf{F}^\top\mathbf{C}_G\right\}_{tf}.$$

Analogously, for $\varepsilon$, the effect of environmental factor $f$ on trait $t$, we can derive that:

$$\operatorname{tr}\left(\mathbf{Z}^{-1}\left(\sum_{j=1}^{N}\mathbf{X}_j^\top\mathbf{V}_j^{-1}\frac{\partial\mathbf{V}_j}{\partial\varepsilon}\mathbf{V}_j^{-1}\mathbf{X}_j\right)\right) \tag{164}$$

$$=\left\{2\mathbf{F}\begin{bmatrix}\operatorname{tr}\left(\mathbf{B}_1^{-1}\mathbf{B}_{E11}\right) & & \mathbf{0}\\ & \ddots & \\ \mathbf{0} & & \operatorname{tr}\left(\mathbf{B}_T^{-1}\mathbf{B}_{ETT}\right)\end{bmatrix}\mathbf{F}^\top\mathbf{C}_E\right\}_{tf}. \tag{165}$$

Overall, computing the second trace can be done in $O\left(NT^2\right)$ time when covariates differ across traits and in $O\left(NT\right)$ time in case of identical covariates across traits. These statements about the computational order rely on $k = O\left(1\right)$.

**Overall complexity of the calculation of the gradient.** We can compute the gradient in $O\left(NT^2\right)$ time, provided $k = O\left(1\right)$. Notice again, however, that having no covariates at all is numerically easier in terms of the degree to which the problem scales with $O\left(NT^2\right)$. The reason is that many terms can be ignored altogether (e.g., $\log|\mathbf{Z}|$) in a model without covariates.

#### Step 8. Calculating the AI matrix efficiently

From the expression of the AI matrix of the log-likelihood in Eq. 25, it follows that we need efficient expressions for weighted squared sum of residuals for each combination of parameters.

**First squared sum of residuals in the AI matrix.** Here, we need to find an efficient expression for $\sum_{j=1}^{N}\mathbf{r}_j^\top\frac{\partial\mathbf{V}_j}{\partial\theta_1}\mathbf{V}_j^{-1}\frac{\partial\mathbf{V}_j}{\partial\theta_2}\mathbf{r}_j$. Central in this derivation is finding an efficient expression for $\frac{\partial\mathbf{V}_j}{\partial\theta_2}\mathbf{r}_j$. For a genetic parameter $\gamma$ and environmental parameter $\varepsilon$, we have:

$$\frac{\partial \mathbf{V}_j}{\partial \gamma}\mathbf{r}_j = d_j \frac{\partial \mathbf{V}_G}{\partial \gamma}\mathbf{r}_j \tag{166}$$

$$= d_j \left[ \begin{pmatrix} \mathbf{0}_{T\times(t-1)} & \mathbf{c}_{Gf} & \mathbf{0}_{T\times(T-t)} \end{pmatrix} + \begin{pmatrix} \mathbf{0}_{(t-1)\times T} \\ \mathbf{c}_{Gf}^\top \\ \mathbf{0}_{(T-t)\times T} \end{pmatrix} \right] \mathbf{r}_j \tag{167}$$

$$= d_j \left[ \mathbf{c}_{Gf} R_{tj} + \begin{pmatrix} \mathbf{0}_{(t-1)\times 1} \\ \mathbf{c}_{Gf}^\top \mathbf{r}_j \\ \mathbf{0}_{(T-t)\times 1} \end{pmatrix} \right] \text{ and} \tag{168}$$

$$\frac{\partial \mathbf{V}_j}{\partial \varepsilon}\mathbf{r}_j = \left[ \mathbf{c}_{Ef} R_{tj} + \begin{pmatrix} \mathbf{0}_{(t-1)\times 1} \\ \mathbf{c}_{Ef}^\top \mathbf{r}_j \\ \mathbf{0}_{(T-t)\times 1} \end{pmatrix} \right]. \tag{169}$$

With $\theta_1$ being a genetic parameter, $\lambda$, denoting the effect of genetic factor $g$ on trait $u$, and $\theta_2$ being a genetic parameter, $\gamma$, denoting the effect of genetic factor $f$ on trait $t$, we obtain that:

$$\sum_{j=1}^{N} \mathbf{r}_j^\top \frac{\partial \mathbf{V}_j}{\partial \theta_1} \mathbf{V}_j^{-1} \frac{\partial \mathbf{V}_j}{\partial \theta_2} \mathbf{r}_j = \sum_{j=1}^{N} \mathbf{r}_j^\top \frac{\partial \mathbf{V}_j}{\partial \lambda} \mathbf{V}_j^{-1} \frac{\partial \mathbf{V}_j}{\partial \gamma} \mathbf{r}_j \tag{170}$$

$$= \sum_{j=1}^{N} d_j^2 \left[ \mathbf{c}_{Gg}^\top R_{uj} + \begin{pmatrix} \mathbf{0}_{1\times(u-1)} & \mathbf{c}_{Gg}^\top \mathbf{r}_j & \mathbf{0}_{1\times(T-u)} \end{pmatrix} \right] \mathbf{V}_j^{-1} \left[ \mathbf{c}_{Gf} R_{tj} + \begin{pmatrix} \mathbf{0}_{(t-1)\times 1} \\ \mathbf{c}_{Gf}^\top \mathbf{r}_j \\ \mathbf{0}_{(T-t)\times 1} \end{pmatrix} \right] \tag{171}$$

$$= \sum_{j=1}^{N} d_j^2 \left[ \left( \mathbf{c}_{Gg}^\top \mathbf{V}_j^{-1} \mathbf{c}_{Gf} R_{uj} R_{tj} \right) + \left( \begin{pmatrix} \mathbf{0}_{1\times(u-1)} & \mathbf{c}_{Gg}^\top \mathbf{r}_j & \mathbf{0}_{1\times(T-u)} \end{pmatrix} \mathbf{V}_j^{-1} \mathbf{c}_{Gf} R_{tj} \right) \right. \tag{172}$$

$$\left. + \mathbf{c}_{Gg}^\top \mathbf{V}_j^{-1} \begin{pmatrix} \mathbf{0}_{(t-1)\times 1} \\ \mathbf{c}_{Gf}^\top \mathbf{r}_j \\ \mathbf{0}_{(T-t)\times 1} \end{pmatrix} R_{uj} + \begin{pmatrix} \mathbf{0}_{1\times(u-1)} & \mathbf{c}_{Gg}^\top \mathbf{r}_j & \mathbf{0}_{1\times(T-u)} \end{pmatrix} \mathbf{V}_j^{-1} \begin{pmatrix} \mathbf{0}_{(t-1)\times 1} \\ \mathbf{c}_{Gf}^\top \mathbf{r}_j \\ \mathbf{0}_{(T-t)\times 1} \end{pmatrix} \right] \tag{173}$$

$$= \sum_{j=1}^{N} d_j^2 \left[ \left\{ \mathbf{C}_G^\top \mathbf{V}_j^{-1} \mathbf{C}_G \right\}_{gf} R_{uj} R_{tj} + \left\{ \mathbf{C}_G^\top \mathbf{R} \right\}_{gj} \left\{ \mathbf{V}_j^{-1} \mathbf{C}_G \right\}_{uf} R_{tj} \right. \tag{174}$$

$$\left. + \left\{ \mathbf{C}_G^\top \mathbf{R} \right\}_{fj} \left\{ \mathbf{V}_j^{-1} \mathbf{C}_G \right\}_{tg} R_{uj} + \left\{ \mathbf{V}_j^{-1} \right\}_{ut} \left\{ \mathbf{C}_G^\top \mathbf{R} \right\}_{gj} \left\{ \mathbf{C}_G^\top \mathbf{R} \right\}_{fj} \right]. \tag{175}$$

Similarly, with $\theta_1$ being an environmental parameter, $\nu$, denoting the effect of environmental factor $g$ on trait $u$, and $\theta_2$ being a genetic parameter, $\gamma$, denoting the effect of genetic factor $f$ on trait $t$, we have that:

$$\sum_{j=1}^{N} \mathbf{r}_j^\top \frac{\partial \mathbf{V}_j}{\partial \theta_1} \mathbf{V}_j^{-1} \frac{\partial \mathbf{V}_j}{\partial \theta_2} \mathbf{r}_j = \sum_{j=1}^{N} \mathbf{r}_j^\top \frac{\partial \mathbf{V}_j}{\partial \nu} \mathbf{V}_j^{-1} \frac{\partial \mathbf{V}_j}{\partial \gamma} \mathbf{r}_j \tag{176}$$

$$= \sum_{j=1}^{N} d_j \left[ \left\{ \mathbf{C}_E^\top \mathbf{V}_j^{-1} \mathbf{C}_G \right\}_{gf} R_{uj} R_{tj} + \left\{ \mathbf{C}_E^\top \mathbf{R} \right\}_{gj} \left\{ \mathbf{V}_j^{-1} \mathbf{C}_G \right\}_{uf} R_{tj} \right. \tag{177}$$

$$\left. + \left\{ \mathbf{C}_G^\top \mathbf{R} \right\}_{fj} \left\{ \mathbf{V}_j^{-1} \mathbf{C}_E \right\}_{tg} R_{uj} + \left\{ \mathbf{V}_j^{-1} \right\}_{ut} \left\{ \mathbf{C}_E^\top \mathbf{R} \right\}_{gj} \left\{ \mathbf{C}_G^\top \mathbf{R} \right\}_{fj} \right]. \tag{178}$$

Finally, for $\theta_1$ being an environmental parameter, $\nu$, denoting the effect of environmental factor $g$ on trait $u$, and $\theta_2$ being an environmental parameter, $\varepsilon$, denoting the effect of environmental factor $f$ on trait $t$, we have that:

$$\sum_{j=1}^{N}\mathbf{r}_j^\top \frac{\partial \mathbf{V}_j}{\partial \theta_1}\mathbf{V}_j^{-1}\frac{\partial \mathbf{V}_j}{\partial \theta_2}\mathbf{r}_j = \sum_{j=1}^{N}\mathbf{r}_j^\top \frac{\partial \mathbf{V}_j}{\partial \nu}\mathbf{V}_j^{-1}\frac{\partial \mathbf{V}_j}{\partial \varepsilon}\mathbf{r}_j \tag{179}$$

$$= \sum_{j=1}^{N}\left[\left\{\mathbf{C}_E^\top \mathbf{V}_j^{-1}\mathbf{C}_E\right\}_{gf} R_{uj}R_{tj} + \left\{\mathbf{C}_E^\top \mathbf{R}\right\}_{gj}\left\{\mathbf{V}_j^{-1}\mathbf{C}_E\right\}_{uf} R_{tj}\right. \tag{180}$$

$$\left. + \left\{\mathbf{C}_E^\top \mathbf{R}\right\}_{fj}\left\{\mathbf{V}_j^{-1}\mathbf{C}_E\right\}_{tg} R_{uj} + \left\{\mathbf{V}_j^{-1}\right\}_{ut}\left\{\mathbf{C}_E^\top \mathbf{R}\right\}_{gj}\left\{\mathbf{C}_E^\top \mathbf{R}\right\}_{fj}\right]. \tag{181}$$

While calculating the log-likelihood, we already obtained $\mathbf{R}$. Moreover, matrices $\mathbf{C}_E^\top\mathbf{R}$ and $\mathbf{C}_G^\top\mathbf{R}$ in the three preceding expressions both need to be computed only once. This calculation can be done in $O\left(NT^2\right)$ time. Thus, the computationally most intensive steps here are obtaining matrices $\mathbf{V}_j^{-1}\mathbf{C}_G$, $\mathbf{V}_j^{-1}\mathbf{C}_E$, $\mathbf{C}_G^\top\mathbf{V}_j^{-1}\mathbf{C}_G$, $\mathbf{C}_E^\top\mathbf{V}_j^{-1}\mathbf{C}_G$, and $\mathbf{C}_E^\top\mathbf{V}_j^{-1}\mathbf{C}_E$. Each of these matrix multiplications can be carried out in at most $O\left(T^3\right)$ time. Thus, across observations, carrying out all these matrix multiplications for $j = 1, \ldots, N$ can be done in $O\left(NT^3\right)$ time.

Bearing all this in mind, we can initialise a $P \times P$ matrix $\overline{\boldsymbol{\mathcal{I}}}^*$ with $P$ denoting the number of free parameters in our model. Here, it holds that $P = O\left(T^2\right)$. For each given observation, $j$, and a given combination of parameters, $\{\theta_1, \theta_2\}$, we can add $\mathbf{r}_j^\top \frac{\partial \mathbf{V}_j}{\partial \theta_1}\mathbf{V}_j^{-1}\frac{\partial \mathbf{V}_j}{\partial \theta_2}\mathbf{r}_j$ to the appropriate element in $\overline{\boldsymbol{\mathcal{I}}}^*$. In this fashion, we can calculate $\sum_{j=1}^{N}\mathbf{r}_j^\top \frac{\partial \mathbf{V}_j}{\partial \theta_1}\mathbf{V}_j^{-1}\frac{\partial \mathbf{V}_j}{\partial \theta_2}\mathbf{r}_j$ for all combinations of parameters in $O\left(NT^4\right)$ time.

**Second squared sum of residuals in the AI matrix.** Here, we need to find an efficient expression for $\left(\sum_{j=1}^{N}\mathbf{r}_j^\top \frac{\partial \mathbf{V}_j}{\partial \theta_1}\mathbf{V}_j^{-1}\mathbf{X}_j\right)\mathbf{Z}^{-1}\left(\sum_{j=1}^{N}\mathbf{X}_j^\top\mathbf{V}_j^{-1}\frac{\partial \mathbf{V}_j}{\partial \theta_2}\mathbf{r}\right)$. In case of identical covariates across traits, this term can be rewritten as:

$$\left(\sum_{j=1}^{N}\mathbf{r}_j^\top \frac{\partial \mathbf{V}_j}{\partial \theta_1}\mathbf{V}_j^{-1}\mathbf{X}_j\right)\mathbf{Z}^{-1}\left(\sum_{j=1}^{N}\mathbf{X}_j^\top\mathbf{V}_j^{-1}\frac{\partial \mathbf{V}_j}{\partial \theta_2}\mathbf{r}\right) \tag{182}$$

$$= \left(\sum_{j=1}^{N}\mathbf{r}_j^\top \frac{\partial \mathbf{V}_j}{\partial \theta_1}\mathbf{F}\mathbf{D}_j^*\mathbf{F}^\top\mathbf{X}_j\right)\mathbf{Z}^{-1}\left(\sum_{j=1}^{N}\mathbf{X}_j^\top\mathbf{F}\mathbf{D}_j^*\mathbf{F}^\top\frac{\partial \mathbf{V}_j}{\partial \theta_2}\mathbf{r}_j\right) \tag{183}$$

$$= \left(\sum_{j=1}^{N}\mathbf{r}_j^\top \frac{\partial \mathbf{V}_j}{\partial \theta_1}\mathbf{F}\mathbf{D}_j^*\mathbf{F}^\top\mathbf{X}_j\left(\left(\mathbf{F}^{-1}\right)^\top \otimes \mathbf{I}_k\right)\right)\mathbf{J}\left(\sum_{j=1}^{N}\left(\mathbf{F}^{-1}\otimes\mathbf{I}_k\right)\mathbf{X}_j^\top\mathbf{F}\mathbf{D}_j^*\mathbf{F}^\top\frac{\partial \mathbf{V}_j}{\partial \theta_2}\mathbf{r}_j\right). \tag{184}$$

Substituting $\mathbf{F}$ by $\mathbf{F} \otimes \mathbf{I}_1$ and $\mathbf{X}_j$ by its definition as Kronecker product, we get that:

$$\left(\sum_{j=1}^{N} \mathbf{r}_j^\top \frac{\partial \mathbf{V}_j}{\partial \theta_1} \mathbf{V}_j^{-1} \mathbf{X}_j\right) \mathbf{Z}^{-1} \left(\sum_{j=1}^{N} \mathbf{X}_j^\top \mathbf{V}_j^{-1} \frac{\partial \mathbf{V}_j}{\partial \theta_2} \mathbf{r}\right) \tag{185}$$

$$= \left(\sum_{j=1}^{N} \mathbf{r}_j^\top \frac{\partial \mathbf{V}_j}{\partial \theta_1} \mathbf{F} \mathbf{D}_j^* \left(\mathbf{F}^\top \otimes \mathbf{I}_1\right) \left(\mathbf{I}_T \otimes \mathbf{x}_j^\top\right) \left(\left(\mathbf{F}^{-1}\right)^\top \otimes \mathbf{I}_k\right)\right) \mathbf{J} \left(\sum_{j=1}^{N} \left(\mathbf{F}^{-1} \otimes \mathbf{I}_k\right) \left(\mathbf{I}_T \otimes \mathbf{x}_j\right) \left(\mathbf{F} \otimes \mathbf{I}_1\right) \mathbf{D}_j^* \mathbf{F}^\top \frac{\partial \mathbf{V}_j}{\partial \theta_2} \mathbf{r}_j\right) \tag{186}$$

$$= \left(\sum_{j=1}^{N} \mathbf{r}_j^\top \frac{\partial \mathbf{V}_j}{\partial \theta_1} \mathbf{F} \mathbf{D}_j^* \left(\mathbf{I}_T \otimes \mathbf{x}_j^\top\right)\right) \mathbf{J} \left(\sum_{j=1}^{N} \left(\mathbf{I}_T \otimes \mathbf{x}_j\right) \mathbf{D}_j^* \mathbf{F}^\top \frac{\partial \mathbf{V}_j}{\partial \theta_2} \mathbf{r}_j\right). \tag{187}$$

Here, $\mathbf{J}$ is as defined in Eq. 54. For $\theta_2$ being a genetic parameter $\gamma$ denoting the effect of genetic factor $f$ on trait $t$, we have:

$$\sum_{j=1}^{N} \left(\mathbf{I}_T \otimes \mathbf{x}_j\right) \mathbf{D}_j^* \mathbf{F}^\top \frac{\partial \mathbf{V}_j}{\partial \gamma} \mathbf{r}_j \tag{188}$$

$$= \begin{pmatrix} \sum_{j=1}^{N} \mathbf{x}_j d_j \left(d_j \lambda_1 + 1\right)^{-1} \left(\left\{\mathbf{F}^\top \mathbf{C}_G\right\}_{1f} R_{tj} + \{\mathbf{F}\}_{t1} \left\{\mathbf{C}_G^\top \mathbf{R}\right\}_{fj}\right) \\ \vdots \\ \sum_{j=1}^{N} \mathbf{x}_j d_j \left(d_j \lambda_T + 1\right)^{-1} \left(\left\{\mathbf{F}^\top \mathbf{C}_G\right\}_{Tf} R_{tj} + \{\mathbf{F}\}_{tT} \left\{\mathbf{C}_G^\top \mathbf{R}\right\}_{fj}\right) \end{pmatrix}. \tag{189}$$

Analogously, for $\theta_2$ being an environmental parameter $\varepsilon$ denoting the effect of environmental factor $f$ on trait $t$, we have:

$$\sum_{j=1}^{N} \left(\mathbf{I}_T \otimes \mathbf{x}_j\right) \mathbf{D}_j^* \mathbf{F}^\top \frac{\partial \mathbf{V}_j}{\partial \varepsilon} \mathbf{r}_j \tag{190}$$

$$= \begin{pmatrix} \sum_{j=1}^{N} \mathbf{x}_j \left(d_j \lambda_1 + 1\right)^{-1} \left(\left\{\mathbf{F}^\top \mathbf{C}_E\right\}_{1f} R_{tj} + \{\mathbf{F}\}_{t1} \left\{\mathbf{C}_E^\top \mathbf{R}\right\}_{fj}\right) \\ \vdots \\ \sum_{j=1}^{N} \mathbf{x}_j \left(d_j \lambda_T + 1\right)^{-1} \left(\left\{\mathbf{F}^\top \mathbf{C}_E\right\}_{Tf} R_{tj} + \{\mathbf{F}\}_{tT} \left\{\mathbf{C}_E^\top \mathbf{R}\right\}_{fj}\right) \end{pmatrix}. \tag{191}$$

When we define the $Tk \times 1$ vectors $\mathbf{w}_\gamma$ and $\mathbf{w}_\varepsilon$ as:

$$\mathbf{w}_\gamma = \begin{pmatrix} \sum_{j=1}^{N} \mathbf{x}_j d_j \left(d_j \lambda_1 + 1\right)^{-1} \left( \left\{ \mathbf{F}^\top \mathbf{C}_G \right\}_{1f} R_{tj} + \{\mathbf{F}\}_{t1} \left\{ \mathbf{C}_G^\top \mathbf{R} \right\}_{fj} \right) \\ \vdots \\ \sum_{j=1}^{N} \mathbf{x}_j d_j \left(d_j \lambda_T + 1\right)^{-1} \left( \left\{ \mathbf{F}^\top \mathbf{C}_G \right\}_{Tf} R_{tj} + \{\mathbf{F}\}_{tT} \left\{ \mathbf{C}_G^\top \mathbf{R} \right\}_{fj} \right) \end{pmatrix} \tag{192}$$

$$\mathbf{w}_\varepsilon = \begin{pmatrix} \sum_{j=1}^{N} \mathbf{x}_j \left(d_j \lambda_1 + 1\right)^{-1} \left( \left\{ \mathbf{F}^\top \mathbf{C}_E \right\}_{1f} R_{tj} + \{\mathbf{F}\}_{t1} \left\{ \mathbf{C}_E^\top \mathbf{R} \right\}_{fj} \right) \\ \vdots \\ \sum_{j=1}^{N} \mathbf{x}_j \left(d_j \lambda_T + 1\right)^{-1} \left( \left\{ \mathbf{F}^\top \mathbf{C}_E \right\}_{Tf} R_{tj} + \{\mathbf{F}\}_{tT} \left\{ \mathbf{C}_E^\top \mathbf{R} \right\}_{fj} \right) \end{pmatrix}, \tag{193}$$

we can stack these vectors into a $Tk \times P$ matrix $\mathbf{W}$. Here, $P$ denotes the number of parameters and we need to make sure that the parameters are indexed in the same manner over the columns of $\mathbf{W}$ as over the columns of $\overline{\mathcal{I}}^*$. The second term of the AI matrix across all parameter combinations is then given by $\mathbf{W}^\top \mathbf{J} \mathbf{W}$. Given that we already finalised the calculation of $\overline{\mathcal{I}}^*$ when computing the first squared sum of residuals in the AI matrix, our final AI matrix can be expressed as:

$$\overline{\mathcal{I}} = \frac{1}{2} \left( \overline{\mathcal{I}}^* - \mathbf{W}^\top \mathbf{J} \mathbf{W} \right). \tag{194}$$

However, these vectors $\mathbf{w}$ can be written more efficiently in terms of Hadamard products:

$$\mathbf{w}_\gamma = \operatorname{vec} \left( \mathbf{X}^\top \left[ \begin{pmatrix} \frac{d_1}{d_1 \lambda_1 + 1} & \cdots & \frac{d_1}{d_1 \lambda_T + 1} \\ \vdots & \ddots & \vdots \\ \frac{d_N}{d_N \lambda_1 + 1} & \cdots & \frac{d_N}{d_N \lambda_T + 1} \end{pmatrix} \circ \left( \{\mathbf{R}\}_{t\bullet} \left\{ \mathbf{C}_G^\top \mathbf{F} \right\}_{f\bullet}^\top + \left\{ \mathbf{C}_G^\top \mathbf{R} \right\}_{f\bullet} \{\mathbf{F}\}_{t\bullet}^\top \right) \right] \right), \tag{195}$$

$$\mathbf{w}_\varepsilon = \operatorname{vec} \left( \mathbf{X}^\top \left[ \begin{pmatrix} \frac{1}{d_1 \lambda_1 + 1} & \cdots & \frac{1}{d_1 \lambda_T + 1} \\ \vdots & \ddots & \vdots \\ \frac{1}{d_N \lambda_1 + 1} & \cdots & \frac{1}{d_N \lambda_T + 1} \end{pmatrix} \circ \left( \{\mathbf{R}\}_{t\bullet} \left\{ \mathbf{C}_E^\top \mathbf{F} \right\}_{f\bullet}^\top + \left\{ \mathbf{C}_E^\top \mathbf{R} \right\}_{f\bullet} \{\mathbf{F}\}_{t\bullet}^\top \right) \right] \right). \tag{196}$$

Here, $\{\mathbf{R}\}_{t\bullet}$, $\left\{ \mathbf{C}_G^\top \mathbf{F} \right\}_{f\bullet}$, etc., are all column vectors. Thus, the second term in each Hadamard product is the sum of two outer products of pairs of vector. Each pair comprises an $N \times 1$ vector and a $T \times 1$ vector, and therefore the outer products are all of size $N \times T$.

$\mathbf{C}_E^\top \mathbf{F}$, $\mathbf{C}_G^\top \mathbf{F}$, $\mathbf{R}$, $\mathbf{F}$, $\mathbf{C}_E^\top \mathbf{R}$, and $\mathbf{C}_G^\top \mathbf{R}$ and their elements are readily available after calculating the log-likelihood and its gradient. Computing the Hadamard products can be done in $O(NT)$ time. Pre-multiplying these Hadamard products by $\mathbf{X}^\top$ also requires $O(NT)$ time, provided $k = O(1)$. Vectorisation and pre-multiplication by the block-diagonal matrix is trivial. As this chain of calculations needs to be carried out for all $P$ parameters, where $P = O\left(T^2\right)$, calculating the full matrix $\mathbf{W}$ requires $O\left(NT^3\right)$ time. Finally, we need to compute $\mathbf{W}^\top\mathbf{J}\mathbf{W}$ in $O\left(T^5\right)$ time. Thus, assuming $T < N$, this means the time complexity of the AI matrix lies in computing $\overline{\mathcal{I}}^*$. This computation requires $O\left(NT^4\right)$ time.

Finally, in case the covariates are not identical across traits we have that:

$$\left(\sum_{j=1}^{N}\mathbf{r}_j^\top\frac{\partial\mathbf{V}_j}{\partial\theta_1}\mathbf{V}_j^{-1}\mathbf{X}_j\right)\mathbf{Z}^{-1}\left(\sum_{j=1}^{N}\mathbf{X}_j^\top\mathbf{V}_j^{-1}\frac{\partial\mathbf{V}_j}{\partial\theta_2}\mathbf{r}\right) \tag{197}$$

$$= \left(\sum_{j=1}^{N}\mathbf{r}_j^\top\frac{\partial\mathbf{V}_j}{\partial\theta_1}\mathbf{F}\mathbf{D}_j^*\mathbf{F}^\top\left(\mathbf{I}_T\otimes\mathbf{x}_j^\top\right)\mathbf{S}^\top\right)\mathbf{Z}^{-1}\left(\sum_{j=1}^{N}\mathbf{S}\left(\mathbf{I}_T\otimes\mathbf{x}_j^\top\right)\mathbf{F}\mathbf{D}_j^*\mathbf{F}^\top\frac{\partial\mathbf{V}_j}{\partial\theta_2}\mathbf{r}_j\right) \tag{198}$$

$$= \left(\sum_{j=1}^{N}\mathbf{r}_j^\top\frac{\partial\mathbf{V}_j}{\partial\theta_1}\mathbf{F}\mathbf{D}_j^*\mathbf{F}^\top\left(\mathbf{I}_T\otimes\mathbf{x}_j^\top\right)\right)\mathbf{S}^\top\mathbf{Z}^{-1}\mathbf{S}\left(\sum_{j=1}^{N}\left(\mathbf{I}_T\otimes\mathbf{x}_j\right)\mathbf{F}\mathbf{D}_j^*\mathbf{F}^\top\frac{\partial\mathbf{V}_j}{\partial\theta_2}\mathbf{r}_j\right) \tag{199}$$

$$= \left(\sum_{j=1}^{N}\mathbf{r}_j^\top\frac{\partial\mathbf{V}_j}{\partial\theta_1}\mathbf{F}\mathbf{D}_j^*\left(\mathbf{I}_T\otimes\mathbf{x}_j^\top\right)\right)\left(\mathbf{F}^\top\otimes\mathbf{I}_k\right)\mathbf{S}^\top\mathbf{Z}^{-1}\mathbf{S}\left(\mathbf{F}\otimes\mathbf{I}_k\right)\left(\sum_{j=1}^{N}\left(\mathbf{I}_T\otimes\mathbf{x}_j\right)\mathbf{D}_j^*\mathbf{F}^\top\frac{\partial\mathbf{V}_j}{\partial\theta_2}\mathbf{r}_j\right) \tag{200}$$

$$= \left(\sum_{j=1}^{N}\mathbf{r}_j^\top\frac{\partial\mathbf{V}_j}{\partial\theta_1}\mathbf{F}\mathbf{D}_j^*\left(\mathbf{I}_T\otimes\mathbf{x}_j^\top\right)\right)\mathbf{J}\left(\sum_{j=1}^{N}\left(\mathbf{I}_T\otimes\mathbf{x}_j\right)\mathbf{D}_j^*\mathbf{F}^\top\frac{\partial\mathbf{V}_j}{\partial\theta_2}\mathbf{r}_j\right), \tag{201}$$

where $\mathbf{J}$ is as defined in Eq. 150. When using the definition of $\mathbf{J}$ as in Eq. 150 (rather than Eq. 54), we get that expressions for the AI matrix with different covariates across traits are the same as expressions for the AI matrix in case of identical covariates.

**Overall complexity of the calculation of the AI matrix.** The time complexity of calculating the AI matrix is linear in the number of observations, and quadratic in the number of parameters. With $P = O\left(T^2\right)$, the AI matrix can be computed in $O\left(NT^4\right)$ time. The computational complexity thus increases rapidly with the number of traits, but our approach seems to yield the lowest time complexity that can be expected to be attainable. The reason is that the number of parameters in the MGREML model increases quadratically with the number of traits considered. In an information matrix, the number of unique elements increases quadratically with the number of parameters. Therefore, the factor $T^4$ cannot reasonably be avoided. Similarly, each individual contributes linearly to the AI matrix as well as to its time complexity. Therefore, the factor $N$ cannot reasonably be avoided either.

#### Step 9. Maximizing the likelihood using a BFGS algorithm

To maximize the likelihood function, we use the quasi-Newton approach of the Broyden-Fletcher-Goldfarb-Shanno (BFGS) algorithm (Nocedal and Wright, 2006). The reason for doing this is that BFGS only needs gradient and function evaluations, and these evaluations can be computed relatively fast. BFGS is a quasi-Newton method where each update takes the form

$$\boldsymbol{\theta}_{k+1} = \boldsymbol{\theta}_k + \alpha_k \mathbf{p}_k. \tag{202}$$

Here, $\alpha$ is the step size of the line search, $k$ is an iteration counter, and $\mathbf{p}_k$ is the search direction

$$\mathbf{p}_k = -\mathbf{B}_k^{-1} \nabla \log l\left(\boldsymbol{\theta}_k\right) \tag{203}$$

with $\nabla \log l\left(\boldsymbol{\theta}\right)$ the gradient of $\log l\left(\boldsymbol{\theta}\right)$. The approximation of the inverse of the Hessian $\mathbf{B}_{k+1}^{-1}$ is defined by:

$$\mathbf{s}_k = \boldsymbol{\theta}_{k+1} - \boldsymbol{\theta}_k = \alpha_k \mathbf{p}_k, \tag{204}$$

$$\mathbf{d}_k = \nabla \log l\left(\boldsymbol{\theta}_{k+1}\right) - \nabla \log l\left(\boldsymbol{\theta}_k\right), \tag{205}$$

$$\rho_k = (\mathbf{s}_k^\top \mathbf{d}_k)^{-1}, \tag{206}$$

$$\mathbf{B}_{k+1}^{-1} = (\mathbf{I} - \rho_k \mathbf{s}_k \mathbf{d}_k^\top) \mathbf{B}_k^{-1} (\mathbf{I} - \rho_k \mathbf{d}_k \mathbf{s}_k^\top) + \rho_k \mathbf{s}_k \mathbf{s}_k^\top. \tag{207}$$

Using these expressions, the BFGS algorithm can be described as:

1. Given start $\boldsymbol{\theta}_0$, convergence tolerance $\varepsilon > 0$, and $\mathbf{B}_0^{-1} = \mathbf{I}$.
2. $k \leftarrow 0$.
3. While $\|\nabla \log l\left(\boldsymbol{\theta}_k\right)\| > \varepsilon$.
4. Compute search direction $\mathbf{p}_k = -\mathbf{B}_k^{-1} \nabla \log l\left(\boldsymbol{\theta}_k\right)$.
5. Set $\boldsymbol{\theta}_{k+1} = \boldsymbol{\theta}_k + \alpha_k \mathbf{p}_k$, with $\alpha_k$ being obtained with a Golden section line search.
6. Compute $B_{k+1}^{-1}$ as in Eq. 207.
7. $k \leftarrow k + 1$.
8. End while

#### Step 10. Computing standard errors using a delta method

The estimation procedure returns the parametrization in terms of $\boldsymbol{\theta}$, that is, in terms of the factor model that underpins $\mathbf{V}_G$ and $\mathbf{V}_E$. In practice, it is often more interesting to analyse the genetic and environmental covariance matrices, the genetic and environmental correlation matrices, and the heritability estimates. In this section, the appropriate standard errors for these transformations are derived.

For some function $g(\boldsymbol{\theta})$, the delta method states that when this function is evaluated at the maximum likelihood estimate, $\widehat{\boldsymbol{\theta}}_{\mathrm{ML}}$, the value this function returns is (approximately) distributed as:

$$g(\widehat{\boldsymbol{\theta}}_{\mathrm{ML}}) \sim \mathcal{N}\left(g(\widehat{\boldsymbol{\theta}}_{\mathrm{ML}}), \nabla g(\widehat{\boldsymbol{\theta}}_{\mathrm{ML}})^\top \mathcal{I}^{-1}(\widehat{\boldsymbol{\theta}}_{\mathrm{ML}}) \nabla g(\widehat{\boldsymbol{\theta}}_{\mathrm{ML}})\right). \tag{208}$$

Here, $\nabla g(\boldsymbol{\theta}) = \partial g(\boldsymbol{\theta})/\partial \boldsymbol{\theta}$ is the gradient of $g()$ with respect to $\boldsymbol{\theta}$. In what follows, the functions $g(\boldsymbol{\theta})$ and their gradients are defined to derive the estimates and standard errors of the SNP heritability, the genetic and environmental correlation matrix, and the genetic and environmental variance-covariance matrix.

**Heritability.** For the SNP heritability of trait $t$, $h_t^2$, we have that:

$$h_t^2 = V_{G_t} \left(V_{G_t} + V_{E_t}\right)^{-1} = \frac{V_{G_t}}{V_t}, \tag{209}$$

where $V_{G_t}$ denotes the genetic variance of trait $t$, $V_{E_t}$ the corresponding environmental variance, and $V_t = V_{G_t} + V_{E_t}$ the total variance of trait $t$. Using, the product rule and the chain rule, we get:

$$\begin{aligned}
\partial h_t^2 &= \partial V_{G_t} \left(V_{G_t} + V_{E_t}\right)^{-1} - V_{G_t} \left(V_{G_t} + V_{E_t}\right)^{-2} \left(\partial V_{G_t} + \partial V_{E_t}\right) && (210)\\
&= \frac{\partial V_{G_t}}{V_t} - \frac{V_{G_t}}{V_t} \frac{\partial V_{G_t} + \partial V_{E_t}}{V_t} && (211)\\
&= \frac{\partial V_{G_t}}{V_t} - h_t^2 \frac{\partial V_{G_t} + \partial V_{E_t}}{V_t} && (212)\\
&= \frac{\left(1 - h_t^2\right) \partial V_{G_t} - h_t^2 \partial V_{E_t}}{V_t}. && (213)
\end{aligned}$$

Thus, quite intuitively, when $h_t^2 = 0$, changes in $V_{E_t}$ will not affect $h_t^2$ (as long as $V_{G_t} = 0$, the heritability will remain zero). Conversely, when $h_t^2 = 1$, changes in $V_{G_t}$ will not affect $h_t^2$ (as long as $V_{E_t} = 0$, the heritability will remain one). Finally, the larger $V_t$ is, the less $h_t^2$ will be affected by changes in either $V_{E_t}$ or $V_{G_t}$.

Let $\mathbf{b}_t$ be a $T \times 1$ binary vector with all elements equal to zero except for element $t$. We then have that:

$$V_{G_t} = \mathbf{b}_t^\top \mathbf{C}_G \mathbf{C}_G^\top \mathbf{b}_t. \tag{214}$$

This expression in turn implies, by the product rule, that:

$$\begin{aligned}
\partial V_{G_t} &= \mathbf{b}_t^\top \left( \partial \mathbf{C}_G \mathbf{C}_G^\top + \mathbf{C}_G \partial \mathbf{C}_G^\top \right) \mathbf{b}_t \\
&= \mathbf{b}_t^\top \partial \mathbf{C}_G \mathbf{C}_G^\top \mathbf{b}_t + \mathbf{b}_t^\top \mathbf{C}_G \left( \partial \mathbf{C}_G \right)^\top \mathbf{b}_t \\
&= \mathbf{b}_t^\top \partial \mathbf{C}_G \mathbf{C}_G^\top \mathbf{b}_t + \left( \mathbf{b}_t^\top \mathbf{C}_G \left( \partial \mathbf{C}_G \right)^\top \mathbf{b}_t \right)^\top \\
&= \mathbf{b}_t^\top \partial \mathbf{C}_G \mathbf{C}_G^\top \mathbf{b}_t + \mathbf{b}_t^\top \partial \mathbf{C}_G \mathbf{C}_G^\top \mathbf{b}_t \\
&= 2\mathbf{b}_t^\top \partial \mathbf{C}_G \mathbf{C}_G^\top \mathbf{b}_t.
\end{aligned}$$

Analogously, we have that:

$$\partial V_{E_t} = 2\mathbf{b}_t^\top \partial \mathbf{C}_E \mathbf{C}_E^\top \mathbf{b}_t. \tag{215}$$

Let $\gamma$ denote the effect of genetic factor $f$ on trait $t$. In this case, we have that:

$$\frac{\partial \mathbf{C}_G}{\partial \gamma} = \begin{cases} 1, & \text{for element} \quad t, f \\ 0, & \text{elsewhere} \end{cases} \tag{216}$$

Thus,

$$\mathbf{b}_t^\top \frac{\partial \mathbf{C}_G}{\partial \gamma} = \begin{pmatrix} \mathbf{0}_{1\times(f-1)} & 1 & \mathbf{0}_{1\times(F_G-f)} \end{pmatrix} \tag{217}$$

This expression reduces to:

$$\mathbf{b}_t^\top \frac{\partial \mathbf{C}_G}{\partial \gamma} \mathbf{C}_G^\top \mathbf{b}_t = \gamma. \tag{218}$$

By means of substituting, we obtain that:

$$\frac{\partial h_t^2}{\partial \gamma} = 2\gamma \frac{1 - h_t^2}{V_t}. \tag{219}$$

Analogously, for the effect $\varepsilon$ of some environmental factor $f$ on trait $t$, we have that:

$$\frac{\partial h_t^2}{\partial \varepsilon} = -2\varepsilon \frac{h_t^2}{V_t}. \tag{220}$$

These parameters can be computed easily for each trait.

**Genetic correlation and environmental correlation.** The genetic correlation between two traits, $t$ and $u$, equals:

$$\rho_{G_{tu}} = \frac{\text{Cov}_{G_{tu}}}{\sqrt{V_{G_t} V_{G_u}}}. \tag{221}$$

Here, $V_{G_t}$ and $V_{G_u}$ denote the genetic variance of traits $t$ and $u$ respectively, and $\text{Cov}_{G_{tu}}$ denotes the genetic covariance of $t$ and $u$. By definition, the correlation is one and the standard error is zero in case $t = u$. Therefore, we focus exclusively on the case where $t \neq u$.

By using the product and chain rule, we get that:

$$\begin{aligned} \partial \rho_{G_{tu}} &= \frac{\partial \text{Cov}_{G_{tu}}}{\sqrt{V_{G_t} V_{G_u}}} - \frac{1}{2} \frac{\text{Cov}_{G_{tu}}}{\sqrt{V_{G_t} V_{G_u}}} \frac{\partial V_{G_t} V_{G_u} + V_{G_t} \partial V_{G_u}}{V_{G_t} V_{G_u}} && (222) \\ &= \frac{\partial \text{Cov}_{G_{tu}}}{\sqrt{V_{G_t} V_{G_u}}} - \frac{1}{2} \rho_{G_{tu}} \frac{\partial V_{G_t} V_{G_u} + V_{G_t} \partial V_{G_u}}{V_{G_t} V_{G_u}}. && (223) \end{aligned}$$

By defining vector $\mathbf{b}_u$ analogously to $\mathbf{b}_t$, substituting expressions found when deriving the gradient of the SNP heritability of $t$, and recognising that:

$$\text{Cov}_{G_{tu}} = \mathbf{b}_t^\top \mathbf{C}_G \mathbf{C}_G^\top \mathbf{b}_u \text{ and} \tag{224}$$

$$\partial \text{Cov}_{G_{tu}} = \mathbf{b}_t^\top \partial \mathbf{C}_G \mathbf{C}_G^\top \mathbf{b}_u + \mathbf{b}_u^\top \partial \mathbf{C}_G \mathbf{C}_G^\top \mathbf{b}_t, \tag{225}$$

we can derive that:

$$\partial \rho_{G_{tu}} = \frac{\mathbf{b}_t^\top \partial \mathbf{C}_G \mathbf{C}_G^\top \mathbf{b}_u + \mathbf{b}_u^\top \partial \mathbf{C}_G \mathbf{C}_G^\top \mathbf{b}_t}{\sqrt{V_{G_t} V_{G_u}}} - \rho_{G_{tu}} \frac{\mathbf{b}_t^\top \partial \mathbf{C}_G \mathbf{C}_G^\top \mathbf{b}_t V_{G_u} + \mathbf{b}_u^\top \partial \mathbf{C}_G \mathbf{C}_G^\top \mathbf{b}_u V_{G_t}}{V_{G_t} V_{G_u}}. \tag{226}$$

Notice that if $t \neq u$, for a parameter $\gamma$ that constitutes the genetic effect of factor $f$ on trait $t$ and a parameter $\lambda$ that constitutes the effect of genetic factor $g$ on trait $u$, the following holds regardless of whether $f = g$ or

$f \neq g$:

$$\frac{\mathbf{b}_t^\top \partial \mathbf{C}_G \mathbf{C}_G^\top \mathbf{b}_t}{\partial \lambda} = 0, \tag{227}$$

$$\frac{\mathbf{b}_u^\top \partial \mathbf{C}_G \mathbf{C}_G^\top \mathbf{b}_u}{\partial \gamma} = 0, \tag{228}$$

$$\frac{\mathbf{b}_t^\top \partial \mathbf{C}_G \mathbf{C}_G^\top \mathbf{b}_u}{\partial \lambda} = 0, \text{ and} \tag{229}$$

$$\frac{\mathbf{b}_u^\top \partial \mathbf{C}_G \mathbf{C}_G^\top \mathbf{b}_t}{\partial \gamma} = 0. \tag{230}$$

Moreover, from the preceding derivations of the gradient of the heritability, we know that:

$$\frac{\mathbf{b}_t^\top \partial \mathbf{C}_G \mathbf{C}_G^\top \mathbf{b}_t}{\partial \gamma} = \gamma, \text{ and} \tag{231}$$

$$\frac{\mathbf{b}_u^\top \partial \mathbf{C}_G \mathbf{C}_G^\top \mathbf{b}_u}{\partial \lambda} = \lambda. \tag{232}$$

Finally, when $f = g$ (i.e., $\lambda$ and $\gamma$ correspond to the same genetic factor), we have that:

$$\frac{\mathbf{b}_t^\top \partial \mathbf{C}_G \mathbf{C}_G^\top \mathbf{b}_u}{\partial \gamma} = \lambda, \text{ and} \tag{233}$$

$$\frac{\mathbf{b}_u^\top \partial \mathbf{C}_G \mathbf{C}_G^\top \mathbf{b}_t}{\partial \lambda} = \gamma. \tag{234}$$

Notice that if $f \neq g$, these two partial derivatives are also zero. By substituting expressions, we get for $t \neq u$ and $f = g$:

$$\frac{\partial \rho_{G_{tu}}}{\partial \gamma} = \frac{\lambda}{\sqrt{V_{G_t} V_{G_u}}} - \rho_{G_{tu}} \frac{\gamma}{V_{G_t}}, \text{ and} \tag{235}$$

$$\frac{\partial \rho_{G_{tu}}}{\partial \lambda} = \frac{\gamma}{\sqrt{V_{G_t} V_{G_u}}} - \rho_{G_{tu}} \frac{\lambda}{V_{G_u}}. \tag{236}$$

Either $\lambda$ or $\gamma$ may be constrained to zero in the given factor model. Finally, it holds that:

$$\frac{\partial \rho_{G_{tt}}}{\partial \gamma} = 0. \tag{237}$$

By analogy, we obtain that:

$$\frac{\partial \rho_{E_{tu}}}{\partial \varepsilon} = \frac{\nu}{\sqrt{V_{E_t} V_{E_u}}} - \rho_{E_{tu}} \frac{\varepsilon}{V_{E_t}}, \tag{238}$$

$$\frac{\partial \rho_{E_{tu}}}{\partial \nu} = \frac{\varepsilon}{\sqrt{V_{E_t} V_{E_u}}} - \rho_{E_{tu}} \frac{\nu}{V_{E_u}}, \text{ and} \tag{239}$$

$$\frac{\partial \rho_{E_{tt}}}{\partial \varepsilon} = 0. \tag{240}$$

In the first equality, $\varepsilon$ is the effect of some environmental factor on trait $t$ and $\nu$ is the effect of that same environmental factor on trait $u$ (if any, because $\nu$ can be constrained to zero in the given factor model). In the second equality, $\nu$ is the effect of some environmental factor on trait $u$ and $\varepsilon$ the environmental effect of that same factor on trait $t$ (if any, because, as before, $\varepsilon$ can be constrained to zero in the given factor model).

**Variance components.** For the estimated factor coefficients in $\mathbf{C}_G$ and $\mathbf{C}_E$, the covariance matrix is readily available. However, sometimes one may be interested in the covariance between the elements in $\mathbf{V}_G$ and $\mathbf{V}_E$. That is, one may be interested in the covariance of the variance components rather than in the factor coefficients. For this case, we consider the genetic (respectively environmental) variance of trait $t$. We have already seen that when $\gamma$ ($\varepsilon$) denotes the effect of genetic (environmental) factor $f$ on trait $t$, it holds that:

$$\frac{\partial V_{G_t}}{\partial \gamma} = 2\gamma, \text{ and} \tag{241}$$

$$\frac{\partial V_{E_t}}{\partial \varepsilon} = 2\varepsilon. \tag{242}$$

Thus, gradient vectors for the genetic and environmental variance can be calculated easily. Let's now consider the genetic covariance between traits $t \neq u$, again letting $\gamma$ denote the effect of genetic factor $f$ on trait $t$ and letting $\lambda$ denote the effect of that same genetic factor $f$ on trait $u$. From before, we know that:

$$\frac{\partial \text{Cov}_{G_{tu}}}{\partial \gamma} = \lambda, \text{ and} \tag{243}$$

$$\frac{\partial \text{Cov}_{G_{tu}}}{\partial \lambda} = \gamma. \tag{244}$$

Similarly, for the environmental covariance between $t \neq u$, with $\varepsilon$ denoting the effect of environmental factor $f$ on trait $t$ and $\nu$ denoting the effect of that same environmental factor on trait $u$, we have that:

$$\frac{\partial \text{Cov}_{E_{tu}}}{\partial \varepsilon} = \nu, \text{ and} \tag{245}$$

$$\frac{\partial \text{Cov}_{E_{tu}}}{\partial \nu} = \varepsilon. \tag{246}$$

Consequently, gradient vectors of genetic and environmental covariances can also be calculated easily.

#### Implementation practicalities

**Controlling for population stratification.** A primary concern in genetic studies is bias resulting from population stratification. To deal with this, a common practice in the GWAS and GREML literature is the inclusion of the lead principal components (PCs) of the GRM as fixed-effect covariates in the model. Instead of using the lead PCs as fixed-effect covariates, which increases the computational burden, MGREML removes the effect of the lead PCs from the data. That is, to control for the $K$ lead PCs, the last $K$ rows of $\mathbf{P}^\top\mathbf{Y}$ in the canonical transformation are ignored. In our software implementation of MGREML, one can set $K$ to be flexible regarding the extent of population stratification in a particular data set.

**Initialising the coefficient matrix $\mathbf{C}_G$.** To launch the optimization algorithm, starting values for the matrix with coefficients $\mathbf{C}_G$ (Eq. 83) are needed. In our software implementation of MGREML, users can supply their own starting values to the optimization algorithm. However, in most cases suitable starting values are not readily available. In that case, the MGREML software sets the starting value for each trait $t$ equal to the square root of the observed phenotypic variance of the trait divided by the number of factors influencing the trait.

**Unbalanced data.** In case a data set is unbalanced (i.e., not all traits and/or relevant control variables are available for all individuals in the data), the mathematical complexity of REML estimation increases dramatically. When we consider the full $(NT) \times 1$ phenotype vector $\mathbf{y}$, associated variance matrix $\mathbf{V}$, and matrix of fixed-effect covariates $\widetilde{\mathbf{X}}$, missing data requires us to keep only the subset of the rows of $\widetilde{\mathbf{X}}$ and $\mathbf{y}$ as well as the rows and columns of $\mathbf{V}$ for which both the phenotypic as well as data on the control variables is available. Under balanced data, $\mathbf{V}$ is a block-diagonal matrix with all blocks being of equal size, *viz.*, $T \times T$ (see Eq. 11). Under unbalanced data, the variance-covariance matrix is still block-diagonal. However, the blocks are then no longer necessarily of equal size. Thus, at first sight, we can no longer use our efficient expressions.

Yet, here we show that by including a set of dummy variables to control for missing data we fit we can still apply our computationally efficient expressions. To see this, we first inspect the case of balanced data more closely. By overloading notation from previous parts, we can denote the singular-value decomposition (SVD) of the matrix of fixed-effect covariates by:

$$\widetilde{\mathbf{X}} = \begin{pmatrix} \mathbf{P}_1 & \mathbf{P}_0 \end{pmatrix} \begin{pmatrix} \boldsymbol{\Theta} \\ \mathbf{0} \end{pmatrix} \mathbf{Q}^\top. \tag{247}$$

Here, it holds that ${\mathbf{P}_0}^\top \widetilde{\mathbf{X}} = \mathbf{0}$. We now define $\mathbf{K} = {\mathbf{P}_0}^\top$, and we let $r = \text{rank}\left(\widetilde{\mathbf{X}}\right)$ denote the rank of $\widetilde{\mathbf{X}}$. then, $\text{rank}\left(\mathbf{P}_0\right) = NT - r$. As a result, the REML log-likelihood function in case of balanced data (ignoring the constant), can be described as:

$$l\left(\boldsymbol{\theta}\right) = -\frac{1}{2}\log\left(\left|\mathbf{K}\mathbf{V}\mathbf{K}^\top\right|\right) - \frac{1}{2}\mathbf{y}^\top\mathbf{K}^\top\left(\mathbf{K}\mathbf{V}\mathbf{K}^\top\right)^{-1}\mathbf{K}\mathbf{y}. \tag{248}$$

Searle et al. (1992) show that:

$$\mathbf{K}^\top\left(\mathbf{K}\mathbf{V}\mathbf{K}^\top\right)^{-1}\mathbf{K} = \mathbf{V}^{-1} - \mathbf{V}^{-1}\widetilde{\mathbf{X}}\left(\widetilde{\mathbf{X}}^\top\mathbf{V}^{-1}\widetilde{\mathbf{X}}\right)^{-1}\widetilde{\mathbf{X}}^\top\mathbf{V}^{-1}. \tag{249}$$

This identity allowed us to formulate the log-likelihood in Eq. 12, for which we developed efficient expressions in the preceding parts. Let $M$ now denote the total number of missing values across phenotypes. Then (overloading notation), the $(NT - M) \times (NT)$ binary matrix $\mathbf{S}$ consisting of zeros and ones with precisely a single one per row and at most a single one per column such that $\mathbf{S}\mathbf{S}^\top = \mathbf{I}_{NT-M}$, effectively selects the observations with non-missing data from $\mathbf{y}$. We can use matrix $\mathbf{S}$ to compute:

$$\mathbf{y}^* = \mathbf{S}\mathbf{y}, \tag{250}$$

$$\mathbf{V}^* = \mathbf{S}\mathbf{V}\mathbf{S}^\top, \text{ and} \tag{251}$$

$$\widetilde{\mathbf{X}}^* = \mathbf{S}\widetilde{\mathbf{X}} = \begin{pmatrix} \mathbf{P}_1^* & \mathbf{P}_0^* \end{pmatrix} \begin{pmatrix} \boldsymbol{\Theta}^* \\ \mathbf{0} \end{pmatrix} \mathbf{Q}^{*\top}. \tag{252}$$

The last expression constitutes the singular value decomposition of $\widetilde{\mathbf{X}}^*$, such that $\mathbf{P}_0^{*\top}\widetilde{\mathbf{X}}^* = \mathbf{0}$. We define $\mathbf{K}^* = \mathbf{P}_0^{*\top}$. With $r = \text{rank}\left(\widetilde{\mathbf{X}}^*\right)$ denoting the rank of $\widetilde{\mathbf{X}}^*$, $\text{rank}\left(\mathbf{P}_0^*\right) = NT - M - r$. Therefore, the REML log-likelihood function in case of unbalanced data (again ignoring the constant) can be described as:

$$l(\boldsymbol{\theta}) = -\frac{1}{2}\log\left(\left|\mathbf{K}^*\mathbf{V}^*\mathbf{K}^{*\top}\right|\right) - \frac{1}{2}\mathbf{y}^{*\top}\mathbf{K}^{*\top}\left(\mathbf{K}^*\mathbf{V}^*\mathbf{K}^{*\top}\right)^{-1}\mathbf{K}^*\mathbf{y}^* \tag{253}$$

$$= -\frac{1}{2}\log\left(\left|\mathbf{K}^*\mathbf{S}\mathbf{V}\mathbf{S}^\top\mathbf{K}^{*\top}\right|\right) - \frac{1}{2}\mathbf{y}^\top\mathbf{S}^\top\mathbf{K}^{*\top}\left(\mathbf{K}^*\mathbf{S}\mathbf{V}\mathbf{S}^\top\mathbf{K}^{*\top}\right)^{-1}\mathbf{K}^*\mathbf{S}\mathbf{y}. \tag{254}$$

At first sight, it seems we can no longer apply the identity by Searle et al. (1992). However, $\mathbf{K}^*\mathbf{S}$ in its entirety can be considered as a design matrix, like $\mathbf{K}$. With missing observations coded with an arbitrary value (e.g., zero) and a set of $M$ dummies that code for the missing observations, $\mathbf{K}^*\mathbf{S}$ is orthogonal to $\widetilde{\mathbf{X}}$. Based on this, $\mathbf{S}^\top\mathbf{K}^{*\top}\left(\mathbf{K}^*\mathbf{S}\mathbf{V}\mathbf{S}^\top\mathbf{K}^{*\top}\right)^{-1}\mathbf{K}^*\mathbf{S}$ can be expressed in terms of a projection matrix. This expression allows to express the log-likelihood under missing data as in Eq. 12, for which we have efficient expressions. In this case, the projection matrix is based on $\widetilde{\mathbf{X}}$ and a set of $M$ dummies that code for the missing data.

More formally, we have an $(NT) \times K$ matrix of covariates $\widetilde{\mathbf{X}}$, an $(NT - M) \times (NT)$ selection matrix $\mathbf{S}$ with elements that are either zero or one and with row-sum $\sum_{j=1}^{NT} S_{ij} = 1\,\forall\, i$ and column-sum $\sum_{i=1}^{(NT-M)} S_{ij} \in \{0,1\}\,\forall j$, and we have an $(NT-M)\times(NT-M-r)$ matrix $\mathbf{P}_0^*$ with orthonormal columns (i.e., $\mathbf{P}_0^{*\top}\mathbf{P}_0^* = \mathbf{I}_{NT-M-r}$) that lie in the null-space of the $(NT - M) \times K$ matrix $\widetilde{\mathbf{X}}^* = \mathbf{S}\widetilde{\mathbf{X}}$, such that $\mathbf{P}_0^{*\top}\widetilde{\mathbf{X}}^* = \mathbf{0}_{(NT-M-r)\times K}$. Finally, $\mathbf{M}$ is an $M \times (NT)$ matrix, defined analogously to $\mathbf{S}$, in such a manner that it selects missing observations rather than non-missing observations as $\mathbf{S}$ does. We can now show that $\mathbf{S}\mathbf{S}^\top = \mathbf{I}_{NT-M}$, $\mathbf{M}\mathbf{M}^\top = \mathbf{I}_M$, $\mathbf{S}\mathbf{M}^\top = \mathbf{0}_{(NT-M)\times M}$, $\mathbf{M}\mathbf{S}^\top = \mathbf{0}_{M\times(NT-M)}$, and $\mathbf{S}^\top\mathbf{S} + \mathbf{M}^\top\mathbf{M} = \mathbf{I}_{NT}$.

**Theorem 1.** *$\widetilde{\boldsymbol{P}_0} = \boldsymbol{S}^\top\boldsymbol{P}_0^*$ lies in the left null space of $\widetilde{\boldsymbol{X}}_M = \left[\widetilde{\boldsymbol{X}},\ \boldsymbol{M}^\top\right]$ and has rank$\left(\widetilde{\boldsymbol{P}_0}\right) =$ rank$(\boldsymbol{P}_0^*) \equiv NT - M - r$.*

*Proof.*

$$\widetilde{\mathbf{P}_0}^\top\widetilde{\mathbf{X}}_M = \mathbf{P}_0^{\top *}\mathbf{S}\left[\widetilde{\mathbf{X}},\ \mathbf{M}^\top\right] \tag{255}$$

$$= \left[\mathbf{P}_0^{\top *}\mathbf{S}\widetilde{\mathbf{X}},\ \mathbf{P}_0^{\top *}\mathbf{S}\mathbf{M}^\top\right] \tag{256}$$

$$= \left[\mathbf{P}_0^{\top *}\widetilde{\mathbf{X}}^*,\ \mathbf{P}_0^{\top *}\mathbf{0}_{(NT-M)\times M}\right] \tag{257}$$

$$= \mathbf{0}_{(NT-M-r)\times(K+M)}. \tag{258}$$

Hence, $\widetilde{\mathbf{P}_0}$ lies in the null-space of $\widetilde{\mathbf{X}}_M$.

$$\operatorname{rank}\left(\widetilde{\mathbf{P}_0}\right) = \operatorname{rank}\left(\widetilde{\mathbf{P}_0}^\top \widetilde{\mathbf{P}_0}\right) \tag{259}$$

$$= \operatorname{rank}\left(\mathbf{P}_0^{*\top} \mathbf{S}\mathbf{S}^\top \mathbf{P}_0^*\right) \tag{260}$$

$$= \operatorname{rank}\left(\mathbf{P}_0^{*\top} \mathbf{P}_0^*\right) \tag{261}$$

$$= \operatorname{rank}\left(\mathbf{P}_0^*\right) \equiv NT - M - r. \tag{262}$$

Hence, $\operatorname{rank}\left(\widetilde{\mathbf{P}_0}\right) = NT - M - r$. □

**Theorem 2.** *rank* $\left(\widetilde{\boldsymbol{X}}_M\right) = r + M$ *and, therefore,* $\widetilde{\boldsymbol{P}_0}$ *spans the null space of* $\widetilde{\boldsymbol{X}}_M$.

*Proof.* $\operatorname{rank}\left(\widetilde{\mathbf{X}}_M\right)$ is the number of independent columns in $\widetilde{\mathbf{X}}_M$. Hence, orthogonalizing one subset of columns of $\widetilde{\mathbf{X}}_M$ with respect to another, non-overlapping subset of columns of $\widetilde{\mathbf{X}}_M$ other does not change the rank. Letting $\mathbf{I} - \mathbf{M}^\top \left(\mathbf{M}\mathbf{M}^\top\right)^{-1} \mathbf{M} = \mathbf{I} - \mathbf{M}^\top \mathbf{M} = \mathbf{S}^\top \mathbf{S}$ denote the orthogonal projection matrix that removes the collinearity with columns of $\mathbf{M}^\top$, we obtain – based on the orthogonalization-argument – that,

$$\operatorname{rank}\left(\widetilde{\mathbf{X}}_M\right) = \operatorname{rank}\left(\left[\widetilde{\mathbf{X}},\ \mathbf{M}^\top\right]\right) \tag{263}$$

$$= \operatorname{rank}\left(\left[\mathbf{S}^\top \mathbf{S}\widetilde{\mathbf{X}},\ \mathbf{M}^\top\right]\right) \tag{264}$$

$$= \operatorname{rank}\left(\mathbf{S}^\top \mathbf{S}\widetilde{\mathbf{X}}\right) + \operatorname{rank}\left(\mathbf{M}^\top\right) \tag{265}$$

$$= \operatorname{rank}\left(\widetilde{\mathbf{X}}^\top \mathbf{S}^\top \mathbf{S}\mathbf{S}^\top \mathbf{S}\widetilde{\mathbf{X}}\right) + \operatorname{rank}\left(\mathbf{M}\mathbf{M}^\top\right) \tag{266}$$

$$= \operatorname{rank}\left(\widetilde{\mathbf{X}}^\top \mathbf{S}^\top \mathbf{S}\widetilde{\mathbf{X}}\right) + \operatorname{rank}\left(\mathbf{I}_M\right) \tag{267}$$

$$= \operatorname{rank}\left(\mathbf{S}\widetilde{\mathbf{X}}\right) + M \tag{268}$$

$$= \operatorname{rank}\left(\widetilde{\mathbf{X}}^*\right) + M = r + M. \tag{269}$$

Given that $\widetilde{\mathbf{X}}_M$ is an $(NT) \times (K + M)$ matrix with $NT \gg K + M \geq r + M$ and with rank equal to $r + M$, its null space is spanned by $NT - M - r$ independent columns. □

The preceding two theorems show that $\widetilde{\mathbf{P}_0}^\top = \mathbf{K}^* \mathbf{S}$ can be regarded as a design matrix, for which the following identity holds:

$$\mathbf{S}^\top \mathbf{K}^{*\top} \left(\mathbf{K}^* \mathbf{S}\mathbf{V}\mathbf{S}^\top \mathbf{K}^{*\top}\right)^{-1} \mathbf{K}^* \mathbf{S} = \mathbf{V}^{-1} - \mathbf{V}^{-1} \widetilde{\mathbf{X}}_M \left(\mathbf{X}_M^\top \mathbf{V}^{-1} \widetilde{\mathbf{X}}_M\right)^{-1} \mathbf{X}_M^\top \mathbf{V}^{-1}, \tag{270}$$

with $\widetilde{\mathbf{X}}_M = \left[\widetilde{\mathbf{X}},\ \mathbf{M}^\top\right]$.

These derivations and proofs show that efficient MGREML estimation can still be applied to unbalanced data sets by treating the data as balanced and including dummy variables for missing values as fixed-effect covariates. In other words, the estimates one obtain with MGREML using this approach will be the same when using the computationally more demanding expressions that arise when directly using unbalanced data.

**Scale and convergence.** Most terms in the log-likelihood function (Eq. 12) scale linearly with the number of individuals $N$, and therefore we optimize over the log-likelihood dived by $N$ in our software implementation of MGREML. This scaling by $N$ enhances the numerical stability of the optimization algorithm. Relatedly, the gradient also scales linearly with the number of individuals $N$. Therefore, we also divide the convergence criterion by $N$ because this criterion is based on the length of the gradient of the log-likelihood.

**Likelihood-ratio test for nested models.** Based on the model in Eq. 2, we can compare nested models using a likelihood ratio test. For example, one could compare the unconstrained model to a model without the genetic component. For two nested models, the likelihood-ratio statistic is given by $2\left(\log l\left(\boldsymbol{\theta}_{\boldsymbol{ML}}\right) - \log l\left(\boldsymbol{\theta}_{\boldsymbol{H_0}}\right)\right)$, with $\boldsymbol{\theta}_{ML}$ the parameters in the unconstrained model and $\boldsymbol{\theta}_{H_0}$ the parameters in the restricted model. The test statistic follows a $\chi^2(k)$ distribution, with $k$ the number of coefficients that is free in the unconstrained model but constrained to zero in the (restricted) null model.

### S2 Supplementary data description

Table S1 provides an overview of the data fields in the UK Biobank used to construct the traits analysed in this study.

Table S1: Overview of the traits analysed in this study, in alphabetical order: Trait, trait description, measurement unit, and UK Biobank data fields used to construct the trait.

| Trait | Trait description | Measurement unit | UK Biobank data field |
|---|---|---|---|
| BMI | Body mass index (logarithm) | $Kg/m^2$ | 21001 |
| Depression score | First principal component of depression intensity and frequency (logarithm) | NA | 2050, 2060, 4609, 4620, 5375, 5386, 2090, 2100 |
| Drinking | Alcoholic drinks consumed per week (logarithm) | Number of units of alcohol per week | 1558, 1568, 1578, 1588, 1598, 1608, 4407, 4418, 4429, 4440, 4451, 4462, 5364 |

| Trait | Trait description | Measurement unit | UK Biobank data field |
|---|---|---|---|
| Educational attainment | Highest self-reported schooling degree converted to US-schooling year equivalents using ISCED categories | years | 6138 |
| Grey matter in amygdala | Volume of grey matter in amygdala (left+right) | $mm^3$ | 25888, 25889 |
| Grey matter in angular gyrus | Volume of grey matter in angular gyrus | $mm^3$ | 25822, 25823 |
| Grey matter in brain-stem | Volume of grey matter in brain-stem | $mm^3$ | 25892 |
| Grey matter in caudate | Volume of grey matter in caudate (left+right) | $mm^3$ | 25880, 25881 |
| Grey matter in central opercular cortex | Volume of grey matter in central opercular cortex (left+right) | $mm^3$ | 25864, 25865 |
| Grey matter in cingulate gyrus, (ad) | Volume of grey matter in cingulate gyrus, anterior division (left+right) | $mm^3$ | 25838, 25839 |
| Grey matter in cingulate gyrus, (pd) | Volume of grey matter in cingulate gyrus, posterior division (left+right) | $mm^3$ | 25840, 25841 |
| Grey matter in crus I cerebellum | Volume of grey matter in crus I cerebellum (left+right) | $mm^3$ | 25900, 25902 |
| Grey matter in crus I cerebellum) | Volume of grey matter in crus I cerebellum (vermis) | $mm^3$ | 25901 |
| Grey matter in crus II cerebellum | Volume of grey matter in crus II cerebellum (vermis) | $mm^3$ | 25904 |
| Grey matter in crus II cerebellum | Volume of grey matter in crus II cerebellum (left+right) | $mm^3$ | 25903, 25905 |
| Grey matter in cuneal cortex | Volume of grey matter in cuneal cortex (left+right) | $mm^3$ | 25844, 25845 |
| Grey matter in frontal medial cortex | Volume of grey matter in frontal medial cortex (left+right) | $mm^3$ | 25830, 25831 |
| Grey matter in frontal operculum cortex | Volume of grey matter in frontal operculum cortex (left+right) | $mm^3$ | 25862, 25863 |
| Grey matter in frontal orbital cortex | Volume of grey matter in frontal orbital cortex (left+right) | $mm^3$ | 25846, 25847 |
| Grey matter in frontal pole | Volume of grey matter in frontal pole (left+right) | $mm^3$ | 25782, 25783 |
| Grey matter in Heschl's gyrus | Volume of grey matter in Heschl's gyrus (includes H1 and H2) (left+right) | $mm^3$ | 25870, 25871 |
| Grey matter in hippocampus | Volume of grey matter in hippocampus (left+right) | $mm^3$ | 25886, 25887 |

| Trait | Trait description | Measurement unit | UK Biobank data field |
|---|---|---|---|
| Grey matter in I-IV cerebellum | Volume of grey matter in I-IV cerebellum (left+right) | $mm^3$ | 25893, 25894 |
| Grey matter in inferior frontal gyrus, po | Volume of grey matter in inferior frontal gyrus, pars opercularis (left+right) | $mm^3$ | 25792, 25793 |
| Grey matter in inferior frontal gyrus, pt | Volume of grey matter in inferior frontal gyrus, pars triangularis (left+right) | $mm^3$ | 25790, 25790 |
| Grey matter in inferior temporal gyrus, (tp) | Volume of grey matter in inferior temporal gyrus, temporooccipital part (left+right) | $mm^3$ | 25812, 25813 |
| Grey matter in inferior temporal gyrus, (ad) | Volume of grey matter in Inferior temporal gyrus, anterior division (left+right) | $mm^3$ | 25808, 25808 |
| Grey matter in inferior temporal gyrus, (pd) | Volume of grey matter in inferior temporal gyrus, posterior division (left+right) | $mm^3$ | 25810, 25811 |
| Grey matter in insular cortex | Volume of grey matter in insular cortex (left+right) | $mm^3$ | 25784, 25785 |
| Grey matter in intracalcarine cortex | Volume of grey matter in intracalcarine cortex (left+right) | $mm^3$ | 25828, 25829 |
| Grey matter in IX cerebellum | Volume of grey matter in IX cerebellum (left+right) | $mm^3$ | 25915, 25917 |
| Grey matter in juxtapositional lobule cortex | Volume of grey matter in juxtapositional lobule cortex (formerly supplementary motor cortex) (left+right) | $mm^3$ | 25832, 25833 |
| Grey matter in lateral occipital cortex, (id) | Volume of grey matter in lateral occipital cortex, inferior division (left+right) | $mm^3$ | 25826, 25827 |
| Grey matter in lateral occipital cortex, (sd) | Volume of grey matter in lateral occipital cortex, superior division (left+right) | $mm^3$ | 25824, 25825 |
| Grey matter in lingual gyrus | Volume of grey matter in lingual gyrus (left+right) | $mm^3$ | 25852, 25853 |
| Grey matter in middle frontal gyrus | Volume of grey matter in middle frontal gyrus (left+right) | $mm^3$ | 25788, 25789 |
| Grey matter in middle temporal gyrus, (tp) | Volume of grey matter in middle temporal gyrus, temporooccipital part (left+right) | $mm^3$ | 25806, 25807 |

| Trait | Trait description | Measurement unit | UK Biobank data field |
| --- | --- | --- | --- |
| Grey matter in middle temporal gyrus, (ad) | Volume of grey matter in middle temporal gyrus, anterior division (left+right) | $mm^3$ | 25802, 25803 |
| Grey matter in middle temporal gyrus, (pd) | Volume of grey matter in middle temporal gyrus, posterior division (left+right) | $mm^3$ | 25804, 25805 |
| Grey matter in occipital fusiform gyrus | Volume of grey matter in occipital fusiform gyrus (left+right) | $mm^3$ | 25860, 25861 |
| Grey matter in occipital pole | Volume of grey matter in occipital pole (left+right) | $mm^3$ | 25876, 25877 |
| Grey matter in pallidum | Volume of grey matter in pallidum (left+right) | $mm^3$ | 25884, 25884 |
| Grey matter in paracingulate gyrus | Volume of grey matter in paracingulate gyrus (left+right) | $mm^3$ | 25836, 25837 |
| Grey matter in parahippocampal gyrus, (ad) | Volume of grey matter in parahippocampal gyrus, anterior division (left+right) | $mm^3$ | 25848, 25849 |
| Grey matter in parahippocampal gyrus, (pd) | Volume of grey matter in parahippocampal gyrus, posterior division (left+right) | $mm^3$ | 25850, 25851 |
| Grey matter in parietal operculum cortex | Volume of grey matter in parietal operculum cortex (left+right) | $mm^3$ | 25866, 25867 |
| Grey matter in planum polare | Volume of grey matter in planum polare (left+right) | $mm^3$ | 25868, 25869 |
| Grey matter in planum temporale | Volume of grey matter in planum temporale (left+right) | $mm^3$ | 25872, 25783 |
| Grey matter in postcentral gyrus | Volume of grey matter in postcentral gyrus (left+right) | $mm^3$ | 25814, 25815 |
| Grey matter in precentral gyrus | Volume of grey matter in precentral gyrus (left+right) | $mm^3$ | 25794, 25795 |
| Grey matter in precuneous cortex | Volume of grey matter in precuneous cortex (left+right) | $mm^3$ | 25842, 25843 |
| Grey matter in putamen | Volume of grey matter in putamen (left+right) | $mm^3$ | 25882, 25883 |
| Grey matter in subcallosal cortex | Volume of grey matter in subcallosal cortex (left+right) | $mm^3$ | 25834, 25835 |
| Grey matter in superior frontal Gyrus | Volume of grey matter in superior frontal gyrus (left) | $mm^3$ | 25786 |

| Trait | Trait description | Measurement unit | UK Biobank data field |
|---|---|---|---|
| Grey matter in superior parietal Lobule | Volume of grey matter in superior parietal lobule (left+right) | $mm^3$ | 25816, 25817 |
| Grey matter in superior temporal gyrus, (ad) | Volume of grey matter in superior temporal gyrus, anterior division (left+right) | $mm^3$ | 25798, 25799 |
| Grey matter in superior temporal gyrus, (pd) | Volume of grey matter in superior temporal gyrus, posterior division (left+right) | $mm^3$ | 25800, 25801 |
| Grey matter in supracalcarine cortex | Volume of grey matter in supracalcarine cortex (left+right) | $mm^3$ | 25874, 25875 |
| Grey matter in supramarginal gyrus, (ad) | Volume of grey matter in supramarginal gyrus, anterior division (left+right) | $mm^3$ | 25818, 25819 |
| Grey matter in supramarginal gyrus, (pd) | Volume of grey matter in supramarginal gyrus, posterior division (left+right) | $mm^3$ | 25820, 25821 |
| Grey matter in temporal fusiform cortex, (ad) | Volume of grey matter in temporal fusiform cortex, anterior division (left+right) | $mm^3$ | 25854, 25855 |
| Grey matter in temporal fusiform cortex, (pd) | Volume of grey matter in temporal fusiform cortex, posterior division (left+right) | $mm^3$ | 25856, 25857 |
| Grey matter in temporal occipital fusiform cortex | Volume of grey matter in temporal occipital fusiform cortex (left+right) | $mm^3$ | 25858, 25859 |
| Grey matter in temporal pole | Volume of grey matter in temporal pole (left+right) | $mm^3$ | 25796, 25797 |
| Grey matter in thalamus | Volume of grey matter in thalamus (left+right) | $mm^3$ | 25878, 25879 |
| Grey matter in V cerebellum | Volume of grey matter in V cerebellum (left+right) | $mm^3$ | 25895, 25896 |
| Grey matter in ventral striatum | Volume of grey matter in ventral striatum (left+right) | $mm^3$ | 25890, 25891 |
| Grey matter in VI cerebellum | Volume of grey matter in VI cerebellum (left+right) | $mm^3$ | 25897, 25899 |
| Grey matter in VI cerebellum) | Volume of grey matter in VI cerebellum (vermis) | $mm^3$ | 25898 |
| Grey matter in VIIb cerebellum | Volume of grey matter in VIIb cerebellum (left+right) | $mm^3$ | 25906, 25908 |

| Trait | Trait description | Measurement unit | UK Biobank data field |
|---|---|---|---|
| Grey matter in VIIb cerebellum) | Volume of grey matter in VIIb cerebellum (vermis) | $mm^3$ | 25907 |
| Grey matter in VIIIa cerebellum | Volume of grey matter in VIIIa cerebellum (left+right) | $mm^3$ | 25909, 25911 |
| Grey matter in VIIIa cerebellum) | Volume of grey matter in VIIIa cerebellum (vermis) | $mm^3$ | 25910 |
| Grey matter in VIIIb cerebellum | Volume of grey matter in VIIIb cerebellum (left+right) | $mm^3$ | 25912, 25914 |
| Grey matter in VIIIb cerebellum) | Volume of grey matter in VIIIb cerebellum (vermis) | $mm^3$ | 25913 |
| Grey matter in X cerebellum | Volume of grey matter in X cerebellum (left+right) | $mm^3$ | 25918, 25920 |
| Grey matter in X cerebellum) | Volume of grey matter in X cerebellum (vermis) | $mm^3$ | 25919 |
| IQ | Standardized fluid intelligence score | correct-answers | 20016, 20191 |
| Neuroticism | Neuroticism standardized score | NA | 1920, 1930, 1940, 1950, 1960, 1970, 1980, 1990, 2000, 2010, 2020, 2030 |
| Reaction time | Standardized reaction time | milliseconds | 20023 |
| Standing height | Standing height | cm | 50 |
| Subjective well-being | Subjective well-being: In general how happy are you? (Average value over time) | NA | 4526, 20458 |
| Visual memory | Standardized visual memory score (logarithm) | NA | 399, 20132 |
| Volume of brain | Volume of brain, grey+white matter | $mm^3$ | 25010 |

In our analysis sample, we removed individuals with brain diseases or surgical brain damage. Table S2 provides an overview of the the brain diseases and ICD10 codes used to make these exclusions.

Table S2: Overview of data fields in the UK Biobank used to exclude individuals with brain diseases or surgical brain damage.

| Trait | UK Biobank data field | ICD10 code |
|---|---|---|
| Dementia or Alzheimer's disease | 1263 | F01, F02, G30 |
| Parkinson's disease | 1262 | G20, G21 |
| Chronic degenerative neurological | 1258 | G23, G31, G32 |
| Guillain-Barré syndrome | 1256 | G610 |
| Multiple Sclerosis | 1261 | G35 |
| Other demyelinating disease | 1397 | G37 |
| Stroke or ischaemic stroke | 1081 | G463, G464, I64, I694 |
| Brain cancer | 1031 | C70, C71, D33 |
| Brain haemorrhage | 1491 | I60, I61, I62, I691, I692, I693 |
| Brain/intracranial abscess | 1245 | G060, G07 |
| Cerebral aneurysm | 1425 | I671, Q282, Q283 |
| Cerebral palsy | 1433 | G80, A521, A504, I64 |
| Encephalitis | 1246 | A83, A86, B011, B020, B262, A85, B004, B582, A84, B050, B941, G04, A321, G05 |
| Epilepsy | 1264 | G40, F803 |
| Head injury | 1266 | S07, T040 |
| Infections of the nervous system | 1244 | A80, A81, A82, A83, A84, A85, A86, A87, A88, A89 |
| Ischaemic stroke | 1583 | G45 |
| Meningeal cancer | 1031 | C70, C793 |
| Meningioma (benign) | 1659 | D33, D32 |
| Meningitis | 1247 | G03, A170, A171, A203, G01, G02, G00, G07 |
| Motor neuron disease (ALS) | 1259 | G122 |
| Neurological injury / trauma | 1240 | |
| Spina bifida | 1524 | Q05, Q760 |
| Subdural haematoma | 1083 | P100 |
| Subarachnoid haemorrhage | 1086 | I60, S066, P103 |
| Transient ischaemic attack | 1082 | G45 |

#### S3 Supplementary results

**Runtime analyses.** Supplementary Fig. S1 provides the results of a runtime comparison between MGREML and pairwise bivariate GREML (Lee et al., 2012b). To facilitate a fair comparison, the runtimes of bivariate GREML were obtained using our own MGREML implementation. The runtimes for pairwise bivariate GREML were extrapolated using the average runtime of five bivariate GREML analyses of randomly selected traits and the notion that one needs $(T \times (T-1))/2$ pairwise bivariate GREML analyses when analyzing $T$ traits. The results show that, given a number of individuals ($N$) and traits ($T$), MGREML is computationally faster than pairwise bivariate GREML. When increasing the number of traits $T$, the running time of pairwise bivariate GREML increases disproportionally as compared to MGREML. Increasing the number of individuals $N$ in the analyses increases the running time of both pairwise bivariate GREML and MGREML with approximately the same rate.

**Comparison with LDSC.** We investigated the statistical efficiency of MGREML by comparing the standard errors (Extended Data Table 2) of the estimated genetic correlations (Extended Data Table 1) to the standard errors one obtains by using bivariate LD-score regression (Bulik-Sullivan et al., 2015) to estimate genetic correlations. For this comparison, we used PLINK 1.9 (Chang et al., 2015) to run a GWAS for each of the 86 traits using the 1,384,830 SNPs that are also used for constructing the GRM, with the same covariates as those used in the MGREML analyses. The summary statistics of these GWASs were used to compute genetic correlations in a pairwise fashion using LDSC v1.0.1 (with the European individuals of the 1000 Genomes data as the reference panel).

The genetic correlations and corresponding standard errors are available in Extended Data Table 5 and Extended Data Table 6. The results show that the heritability estimates ($\rho = 0.95$) and genetic correlations ($\rho = 0.90$) obtained using MGREML and LDSC are strongly correlated. Importantly, the 95% confidence intervals obtained with MGREML are, compared to the LDSC results, on average 38.9% smaller for the heritability estimates and 46.0% smaller for the genetic correlation estimates.

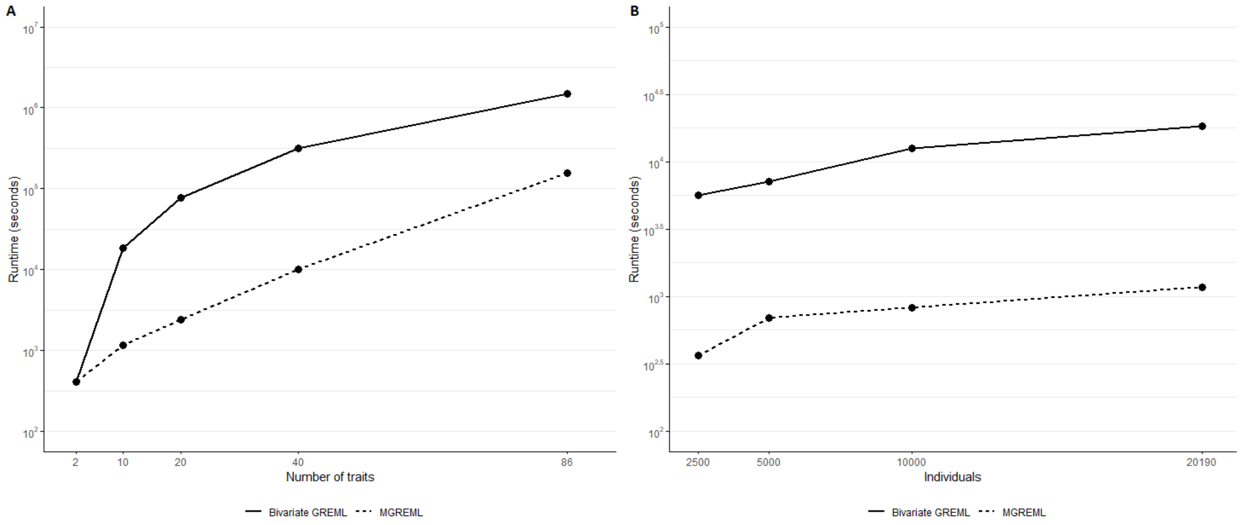


Figure S1: Runtime in seconds of MGREML and pairwise bivariate GREML as function of the number of individuals ($N$) and traits ($T$) in the analysis. $N$=20,190 in Figure S1A; $T$=10 in Figure S1B.

### S4 Analysis pipeline

In this section, we describe the analysis pipeline used to obtain the empirical results as reported in the main text:

1. Restrict to individuals of European ancestry.

2. Exclude the individuals with brain damage (see Table S2 for UK Biobank data cells used to exclude these individuals).

3. Restrict dataset to directly genotyped HapMap3 SNPs + SNPs with imputation quality of 0.9 or higher.

   - This step leaves a total of $1,796,892$ directly genotyped SNPs / SNPs with high imputation quality.

4. Apply regular quality control on SNP data: minor allele frequency (MAF) $< 0.01$, missingness per individual (MIND) $< 0.05$, missingness per SNP (GENO) $< 0.05$, Hardy-Weinberg equilibrium (HWE) $p$-value $< 0.001$.

   - This step leaves a total of $1,582,522$ SNPs.

5. Drop long-range LD (linkage disequilibrium) regions, following Linnér et al. (2019).

- This step leaves a total of $1,384,830$ SNPs.

6. Construct GRM, and apply relatedness cutoff of 0.025 using PLINK (Chang et al., 2015).
   - This step leaves a total of $N = 37,392$ individuals.
7. Curate phenotype data (including construction of genotyping platform dummy variable, which we use as control variable for each trait)
   - Exclude individuals with missing phenotypes: Our choice for fully balanced data leaves us with $N = 20,190$ individuals.
   - Curate covariates for the 74 brain volume phenotypes and the 2 global measures of brain volume: age, age$^2$, sex, sex$\times$age, head size, head motion while resting, head motion in task, date, date$^2$, UK Biobank Assessment Centre, the interactions between age-date$^2$ and UK Biobank Assessment Centre, and a constant.
   - Curate covariates for the 10 other traits: age, age$^2$, sex, sex$\times$age, head size, date, date$^2$, UK Biobank Assessment Centre, the interactions between age-date$^2$ and UK Biobank Assessment Centre, and a constant.
     - For IQ, instead of age we use age at the moment of the IQ assessment and we additionally include dummy variables for the number of IQ measurements used to construct this variable.
8. Run MGREML (with adjustment for the first 20 lead PCs).
